# Influence of the lab adopted natural diet and environmental and parental microbiota on life history and metabolic phenotype of *Drosophila melanogaster* larvae

**DOI:** 10.1101/2020.06.16.154823

**Authors:** Andrei Bombin, Owen Cunneely, Kira Eickman, Sergei Bombin, Abigail Ruesy, Mengting Su Abigail Myers, Rachael Cowan, Laura Reed

## Abstract

Obesity is an increasing worldwide epidemic and contributes to physical and mental health losses. The development of obesity is caused by multiple factors including genotype, hormonal misregulation, psychological stress, and gut microbiota. Our project investigated the effects produced by microbiota community, acquired from the environment and horizontal transfer, on traits related to obesity. The study applied a novel approach of raising *Drosophila melanogaster* from ten, wild-derived genetic lines (DGRP) on naturally fermented peaches, thereby preserving genuine microbial conditions. Our results indicated that larvae raised on the natural and standard lab diets were significantly different from each other in every tested phenotype. In addition, sterilized larvae raised on the autoclaved peach diet, therefore exposed to natural nutritional stress but lacking natural microbiota community, were associated with adverse phenotypes such as low survival rate, longer developmental time, smaller weight, and elevated triglyceride and glucose levels. Our findings suggested that frozen peach food provided nutritional conditions similar to the natural ones and preserved key microbial taxa necessary for survival and development of *Drosophila* larvae. The presence of parental microbiota did not produce a significant effect on any of the tested phenotypes when larvae were raised on the lab diet. Contrarily, on the peach diet, the presence of parental microbiota increased the weight and development rate, even if the original peach microbiota were still present. In addition, we found that larvae raised on the peach diet formed a microbial community distinctive from larvae raised on the lab or peach autoclaved diets. The effect that individual microbial taxa produced on the host varied significantly with changing environmental and genetic conditions, occasionally to the degree of opposite correlations.

## Introduction

The holobiont theory states that a host and its commensal microbiota possess a metagenome that expresses a synergistic phenotype, which is subjected to evolutionary forces as one complex organism (Rosenberg et al., 2010). A phenotype of this unit could be varied by genome modifications of the host, as well as its commensal bacteria. Metagenomic changes induced by bacteria have more potential for genetic variability and could arise by altering dominant species of bacteria, as well as acquisition of new strains of microorganisms from an environment (Bordenstein and Theis, 2015).

Obesity is a worldwide epidemic that continues to grow and contributes to the development of various diseases, including but not limited to: type two diabetes mellitus, stroke, asthma, arthritis, coronary heart disease, arterial hypertension, all components of metabolic syndrome (Seganfredo et al., 2017, Wahba and Mak, 2007). Obesity does not only induce health risks but also is an important social factor. Stereotypically, the major cause of obesity is considered to be overeating and lack of exercise. Obese individuals are often stigmatized as lazy and unsuccessful, making them vulnerable to discrimination (Puhl and Heuer, 2010). Contrary to the stereotypic view, the development of obesity and metabolic syndrome is caused not only by excessive calorie intake and lack of physical activity but also by genotype, epigenetic factors, sleep deprivation, malfunction of endocrine system, psychological stress, and gut microbiota (Seganfredo et al., 2017, Han and Lean, 2016).

Gut microbiota is one of the most important factors shaping metabolic phenotype and, as the consequence is a key element in the development of metabolic and autoimmune diseases, cancer, and asthma (Read and Holmes, 2017, Leitão-Gonçalves et al., 2017). Alterations in gut microbiota biodiversity and community structure are correlated with the development of the obese phenotype (Turnbaugh et al., 2006, Flint et al., 2017). Transfer of the microbiota from an obese to a lean, axenic (microbiota free) individual significantly increases weight gain and adiposity, compared to axenic mice colonized with microbiota community from a lean organism (Ridaura et al., 2013, Tilg and Moschen, 2016, Turnbaugh et al., 2006). These results suggest that obesity can be transferred from one individual to another; therefore, exhibiting some characteristics of an infectious disease (Tilg and Moschen, 2016).

During colonization of a fruit, *Drosophila* inoculates the substrate with their microbiota (Morais et al., 1995). In addition, when females deposit embryos on a substrate, parental microbiota resides on the chorion of the egg (Ryu et al., 2008). The initial microbiota is gained by the first instar larvae through consumption of the chorion (Ryu et al., 2008). Later, during feeding, larvae acquire additional microbiota from the environment, representatives of which remain in the larval and pupal intestine until eclosion of the adult fly (Ridley et al., 2012). Wong et. al (2015) showed that parental microbiota transferred with the chorion of the egg could modify the microbial community composition in a food substrate and in the offspring. In addition, the transfer of axenic *Drosophila* on food substrate would change the food microbial community to resemble the symbiotic microbiota composition that would develop in the host (Wong et al., 2015).

Symbiotic microbiota play an important role in *D. melanogaster* development and metabolic phenotype. Axenic flies have a longer development time, lower weight, protein, and glycogen content but higher free glucose and triglyceride levels (Newell and Douglas, 2014b, Dobson et al., 2015, Huang and Douglas, 2015). There are similar negative phenotypic effects on axenic *D. melanogaster* and on those being raised on harmful diets such as high sugar and high fat (Newell and Douglas, 2014a, Huang and Douglas, 2015, Reed et al., 2014, Birse et al., 2010, Dew-Budd et al., 2016). Although axenic *Drosophila* consume less food, their energy storage indices (triglyceride, glucose, glycogen, and trehalose levels) stay significantly higher than that of conventional flies (Wong et al., 2014). In addition, the presence of the commensal microbiota allows *Drosophila* to maximize their lifespan and reproductive output (Leitão-Gonçalves et al., 2017). Dobson et. al (2015) showed that the abundance of specific bacterial taxa is associated with a change in metabolic phenotype: *Acetobacter, Gluconobacter* and *Komagataibacter* are negatively correlated with a fly’s energy storage index. *Lactobacillus* is positively associated with triglyceride concentration and *Achromobacter* and *Xanthomonadaceae* have a positive correlation with glycogen levels (Chaston et al., 2014, Chaston et al., 2016). In addition, *Drosophila* microbiota varies across genetic backgrounds which makes it possible to establish associations between a host’s genes and microbiota dependent metabolic responses as well as particular symbiotic species (Chaston et al., 2016, Dobson et al., 2015, Early et al., 2017).

Symbiotic microbiota allow *Drosophila* to overcome the nutritional limitation of their diet. Shin et. al (2011) showed that axenic larvae raised on a casamino acid diet experienced a 90% body size reduction and were not able to survive to form pupae. However, the presence of only one bacterial species, *Acetobacter pomorum*, could restore survival and the normal rate of larval development via induction of the hosts’ insulin-like growth factor signaling (Shin et al., 2011). Leitão-Gonçalves et. al (2017) demonstrated that axenic flies express a strong preference for yeast-rich food due to their demand for essential amino acids. *A. promorum* and several *Lactobacilli* species are able to suppress the yeast appetite in *Drosophila* and shift the flies’ nutritional preference toward high-sugar concentration diets (Leitão-Gonçalves et al., 2017). Their change in nutritional preference may be explained by competition between the host and its symbiotic microbiota for available sugars and through production of essential amino acids by the microbial community (Leitão-Gonçalves et al., 2017).

Microbiota composition of lab dwelling *Drosophila* primarily consists of Acetobacteraceae, Enterococcaceae, and Lactobacillaceae families (Chandler et al., 2011, Early et al., 2017). Within these families, the influence of *Acetobacter tropicalis, Enterococcus faecalis, Lactobacillus brevis*, and *Lactobacisul plantarum* on a host’s metabolic phenotype have been studied more than others (Early et al., 2017, Shin et al., 2011, Huang and Douglas, 2015). However, the microbiota of wild fly populations may differ in the diversity and the abundance of dominant species (Chandler et al., 2011). In the lab, *Drosophila* raised on fruits still exhibit a more complex and diverse community of symbiotic microbes compared to conventionally raised flies (Vacchini et al., 2017). Lab food preservatives, especially methylparaben sodium salt (moldex), largely contribute to the difference between natural and laboratory associated microbiota communities (Tefit et al., 2017). Therefore, studying the evolutionary relationship of *Drosophila* and its microbiota, as well as the symbiont’s influence on fly’s metabolic phenotype only on standard lab microbiota, may be insufficient to understand the natural relationship and co-evolution of the fly and its microbiota.

With this work we wanted to address a series of specific questions and hypotheses:

1. How does a natural diet with a naturally occurring and/or maternally inherited community of microbes influence the life history and metabolic phenotypes of flies relative to a standard lab diet? Is there genetic variation in the phenotypic response to nutritional change and dietary and parental microbiota availability?
  1. We hypothesized that larvae raised on a natural diet will exhibit different life history traits and metabolic phenotypes, comparing with larvae raised on a standard lab diet.
  2. We hypothesized that the presence of a maternally transmitted microbiota will significantly impact larvae phenotypes, and that this impact may vary across dietary treatments.
  3. Given prior findings on the roles of genetic variation on metabolic phenotypes, we hypothesized that there is genetic variation in phenotype that interacts with the dietary conditions and the availability of maternally transmitted microbiota.
2. Will the symbiotic microbiota community of the larvae raised on the natural diet be different from the lab food raised larvae? Will the presence of maternally inherited microbiota influence the formation of microbial communities? Is the microbiota community variable across host genotypes?
  1. We hypothesized that the gut microbial community composition and diversity will vary substantially across both dietary and parental microbiota conditions.
  2. We hypothesized that the maternally transmitted microbiota will have “founder effects” in the formation of the larval gut microbiome.
  3. We hypothesized that the composition of the microbial community will exhibit variation with host genotype.
3. Will specific microbial taxa and/or microbiota communities as a whole influence the larvae phenotypes differently across diets and genotypes?
  1. We hypothesized that some microbial taxa will have consistent correlations with host phenotype across diets and treatments while others will have a diet or treatment specific relationship.

## Materials and Methods

### Diets preparation

On August 28^th^, 2017, we put approximately 200 peaches outdoors and allowed them to decay for six days. On September 3^rd^ 2017, the fruits were collected, manually ground, and stored in freezers at −20 °C. Peach food (PR) preparation protocol was the following: we allowed approximately one liter of the peach food to thaw, homogenized it with an immersion blender, and distributed it into vials, with approximately 10 ml of food per vial. In order to prepare autoclaved peach food (PA), vials containing peach food were autoclaved for 25 min at 121 °C. Regular *Drosophila* lab food (R) was cooked according to the protocol described in previous works (Dew-Budd et al., 2016, Mendez et al., 2016).

To ensure that the autoclaved peach food did not contain any live microorganisms, we used 1g of autoclaved and non-autoclaved peach materials and diluted them in 9 ml of sterile Phosphate-buffered saline solution (PBS) (Leitão-Gonçalves et al., 2017, Carvalho et al., 2005). Then, we performed a serial dilution in PBS to get food dilutions (Leboffe and Pierce, 2012). We mixed 1 ml of each dilution with standard methods agar (Criterion) via the pour plate method (Leboffe and Pierce, 2012). The agar was prepared according to the manufacturer’s directions. Samples were incubated at 35 °C for 48 hours (Maturin et al., 2001). The independent variable for the diet component will be referred to as D.

### Drosophila stocks and husbandry

We used 10 naturally derived genetic lines created by the DGRP2 project: 142, 153, 440, 748, 787, 801, 802, 805, 861, and 882 (Mackay et al., 2012, Huang et al., 2014). Stocks were maintained at constant temperature, humidity and light/dark cycle on a molasses-based lab diet as described in previous works (Reed et al., 2014, Dew-Budd et al., 2016, Mendez et al., 2016). The independent variable for the genetic component will be referred to as G.

### Drosophila embryos sterilization

In order to remove parental microbiota, we sterilized ∼12-hour old embryos with subsequent two-minute washes in 2.5% active hypochlorite solution, 70% ethanol solution, and sterilized distilled water (Leitão-Gonçalves et al., 2017, Carvalho et al., 2005). After sterilization, embryos were placed on the apple agar plates, and incubated until the first instar stage under fly-rearing conditions described above (Ashburner, 1989). The non-sterilized control embryos (NS) were allowed to develop for ∼24 hours (until the 1^st^ instar larvae stage) on the apple agar plates, on which they had been deposited. In order to demonstrate that sterilized embryos did not possess parental microbiota, 20 sterilized 1^st^ instar larvae were collected and grinded in 200 ul of the sterile PBS using a mechanical homogenizer. The resulting mixtures were plated on nutrient and standard method agars (Criterion). The agars were prepared according to the manufacturer’s directions. Plates were incubated for 48 hours at 35 °C (Maturin et al., 2001). NS larvae were used as the positive control according to the same procedure. The independent variable for the sterilization treatment component will be referred to as T.

### Larvae rearing and collection

In three separate time periods (∼ 30 days apart), we put 50 sterilized and non-sterilized larvae of each genetic line in at least three vials of PA, PR and R food, each. The independent variable for the time period (round) component will be referred to as R. Larvae were allowed to develop until the late third instar wandering stage (when they stopped moving but before the pupation started) and then collected in micro centrifuge tubes with sterile Ringer’s solution (Ashburner, 1989). Each vial was checked for the presence of larvae, at the right developmental stage, four times per day at 9 am, 11 am, 2:30pm, and 5 pm for 18 days after larvae colonization. Larvae were inspected for the presence of any damage (damaged ones were sorted out), cleaned with at least two washes in a sterile Ringer’s solution, and stored in the Ringer’s solution at −20 °C in Eppendorf tubes with 10 larvae per tube.

### Measuring Experimental Phenotypes

**Survival**. The number of larvae collected per vial was summed and used to evaluate the number of larvae that survived till the late 3^rd^ instar stage. **Developmental rate**: The developmental rate, in days, was calculated for each larva individually, from the day it was put in the food vial to the collection date. We then calculated median developmental time per vial and used it in our statistical analysis (Ridley et al., 2012). **Weight**: In order to measure the dry weight, larvae were taken from the −20 °C freezer, allowed to reach room temperature and placed in a VWR standard oven at 37 °C overnight. After drying, each larva was weighed individually using Mettler Toledo XS 105 microbalance. Weights were recorded with LabX direct software v. 2.2. **Triglyceride**: With the exception of four samples (due to low survival of larvae of certain D/G/T/R combinations), we homogenized 10 larvae per sample to determine total triglyceride concentration using the Sigma Triglyceride Determination Kit (Clark and Keith, 1988, De Luca et al., 2005, Reed et al., 2010, Dew-Budd et al., 2016). Results were adjusted to represent the average triglyceride level per mg of dry larval weight. **Protein**: Protein levels were quantified using the Bradford’s method with 10 homogenized larvae per sample (with the exception of 3% of the samples in which we used one to nine larvae, due to especially low survival rates of the specific groups) (Bradford, 1976, Dew-Budd et al., 2016). Protein levels were averaged to represent the protein concentration per mg of dry larvae weight. **Glucose**: For most of the samples, combined trehalose and glucose concentrations were quantified via homogenization of 10 larvae (with the exception of 5% of the samples in which we used four to nine larvae) with subsequent overnight incubation in 1 μg/mL trehalase solution and further application of the Sigma Glucose Determination Kit (Rulifson et al., 2002, Reed et al., 2014, Dew-Budd et al., 2016). Glucose levels were averaged and adjusted to represent the amount of glucose per mg of dry weight.

To assess the triglyceride, protein and glucose levels in the diets, we used freshly unfrozen food and unfrozen food that was incubated in the food vials for seven days at 25 °C, in the same incubator as the experimental fly stocks, where larvae development was taking place. For the food assays, we used the same procedure as for the larvae but with ∼12.35 mg of the food sample. For the glucose assay, instead of incubating samples in trehalase, we incubated them in 100μl of 1 mg/μl solution of invertase (to convert sucrose into glucose) (Ward’s Natural Science) overnight at 37 °C. For analysis, the results were adjusted to represent the amount of the measured compound per mg of the sample.

### DNA extraction and sequencing

DNA was extracted from 10 larvae with the Qiagen blood and tissue DNA extraction kit according to the standard protocol, with overnight incubation of the samples in proteinase K at 56 °C. DNA extractions were used for sequencing the V4 region of the 16S ribosomal RNA subunit (16S rRNA), which was performed in the Microbiome Core Facility of The University of Alabama in Birmingham, AL according to the previously published method on the Illumina MiSeq platform (Kumar et al., 2014).

Trimmomatic-0.36 (Bolger et al., 2014) was used to process demultiplexed DNA sequences. Standard Illumina-specific barcode sequences, all sequences with less than 36 bases, and leading and trailing low-quality bases were all removed using the default settings of the Trimmomatic-0.36 program. The USEARCH-fastq_mergepairs tool was used to combine forward and reverse readings. All reads with an expected error greater than 1 were removed, as well as chimeric reads and singletons. Combined readings without a merging pair were filtered using fastq_filter command. The -cluster_otus tool was used to cluster readings into operational taxonomic units (OTUs) with 97% identity. The -unoise3 tool was used to cluster readings into zero-radius OTU (ZOTU) 100% identity. OTUs were then designated with the lowest taxonomic rank using the UCULT algorithm implemented in QUIME 1.9.1 (Caporaso et al., 2010b, Edgar, 2010) along with SILVA reference database version 132 (Quast et al., 2012). Using SILVA v. 132 database, PyNAST (Caporaso et al., 2010a) with default options was used for sequence alignment. The phylogenetic trees of ZOTUs and OTUs were assembled using the default options of QUIIME 1.9.1 with the FastTree program (Price et al., 2009). Alpha and beta diversities were rarefied with QUIIME 1.91 -single_rarefaction.py using the --subsample_multinomial option in order to subsample the replacements. Rarefaction for all samples was performed to the depth of 4,500 readings. This was the lowest possible number of readings between samples.

### Statistical Analysis

#### Data transformation

Normality tests, data transformations and statistical models were done with JMP Pro 14.0. Phenotype measurements were tested for normality with the Shapiro-Wilk test and an outlier box plot. Only larvae survival (**ST. 1**) showed normal distribution. Therefore, all other phenotypic measurements data were transformed. We performed a cube root transformation on the data for development rate and glucose by weight, a square root transformation on data for weight and protein by weight, and a log transformation on data for triglyceride by weight levels. The microbial abundance was log(x+1) transformed for all parametric analyses.

#### Microbial Diversity

Alpha and beta diversities were computed in QIIME v. 1.9.1. To estimate alpha diversity, we used Shannon, Simpson, and PD Whole Tree metrices. As all of the alpha diversity indices were not normally distributed, we performed a pairwise comparison of them with a Wilcoxon rank-sum test in R v. 3.5.1 with “matrixTests” package v. 0.17 (Bruno et al., 2019). Beta diversity was estimated with Bray-Curtis and Weighted Unifrac distances. The similarity between each sample’s beta diversity distance was evaluated via hierarchical clustering, applying a ward method for distance calculation, and visualized with a constellation plot in JMP v. 14.0.

#### Statistical modeling

In order to evaluate the contribution of each variable and their interactive effect on each phenotypic development, we used standard least squares model with model effects to include Diet (D), Genotype (G), Sterilization Treatment (T) and their specific interactive effect: D*G (diet by genotype), D*T (diet by treatment), G*T (genotype by treatment), D*G*T (diet by genotype by treatment). In order to verify that the built model fits the data, a Lack of Fit test was performed. If the time period of the experiment (R) and/or the variance between the colorimetric assay runs (triglyceride, protein, and glucose) (P) produced a significant effect, these variables were included in the model’s effects, unless their addition caused the model to fail the lack of fit test. Thus, the models for larvae survival, development time, and weight were the following:

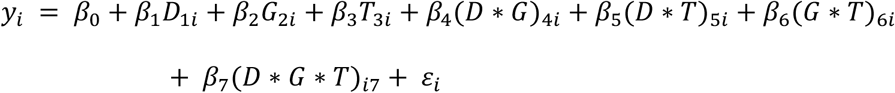

For triglyceride by weight:

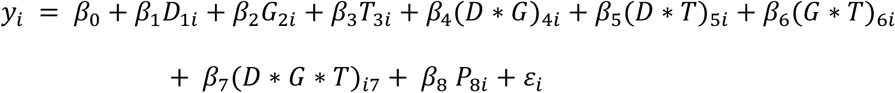

And for protein and glucose by weight:

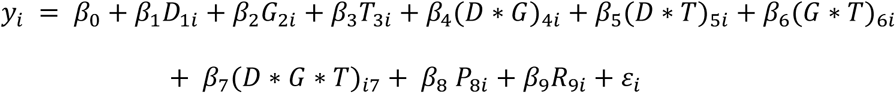

Where *y*_*i*_ is the response, *β* values are constants, and *εi* is a random error term.

All models for interactive effects of diet, genotype, and treatment were done with all 10 genetic lines, with the exception of glucose which was done without 861 due to the low survival rate of this genetic line.

In order to verify that the built model fits the data, a Lack of Fit test was performed. Only the models with non-significant lack of fit p-value were kept and used for an evaluation. To assess the pairwise difference between diets (R vs PR and PR vs PA) and treatments (NS vs S), we used the least square model with one main explanatory variable of interest (Diet or Treatment). We also included time period and assay plate variance as additional explanatory variables if they produced a significant effect. We then used Post-Hoc pairwise comparisons to evaluate the Student’s t-test.

#### Microbial abundance

In order to evaluate if the diet and treatment could serve as categorical predictors for classification of the larvae microbial samples, we performed discriminant analysis at phylum, class, order, family, and genus taxonomic levels, as well as at the level of individual ZOTUs. The results were visualized with a canonical plot in JMP v. 14.0 (JMP manual). For the 10 most abundant representatives of each taxonomic level, we applied the linear covariance method for the discriminant analysis which allowed us to visualize the covariates in the form of rays. Using this method allowed us to represent which of them drove the separation of the clusters (JMP manual). When we ran the analysis with all identified taxa, we applied a wide linear method for the discriminant analysis. To compare the abundance of microbial taxa between the diets and treatments, we performed a Wilcoxon test as described above. The p values were adjusted for the false discovery rate with Benjamini-Hochberg correction and added to data tables as FDR p. The threshold for the value of FDR p that should be considered significant could be subjective and vary from 0.25 to 0.05 among microbiology studies (Wu et al., 2011, Bruce-Keller et al., 2015). To evaluate the interactive effect of the variables on the abundance of each identified microbial taxa, we used the three-way interaction model.

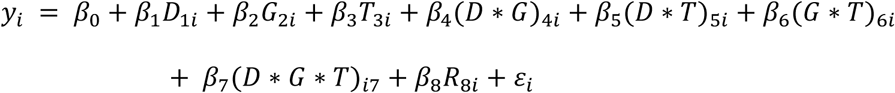

In order to identify the correlations between phenotypes and microbial abundances, we found the average phenotype for each combination of diet, genetic line, treatment, and round and aligned it with the microbial sequences of the corresponding combination of independent variables. Spearman’s rank correlation between the microbial abundances and tested phenotypes was calculated with Hmisc v. 4.3-0 in R v. 3.5.1, with the adjustment of p values for FDR p as described above. We also tested the possible interactive effect of each identified microbial taxa and one of the independent variables on the formation of the tested phenotypes according to the formula:

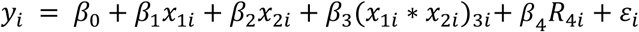

where, *x*_1_ was the abundance of the microbial taxa, *x*_2_was one of the independent variables (D, G, or T) and R was time component. Development, weight, triglyceride, protein, and glucose were normalized with log, square root, log, log, and cube root, transformations respectively.

## Results

### 1) The influence of a natural diet with a naturally occurring and/or maternally inherited community of microbes on the life history and metabolic phenotypes of flies relative to a standard lab diet

**1.1** Larvae raised on a natural diet exhibited different life history traits and metabolic phenotypes compared to larvae raised on a standard lab diet. **Survival**. The number of larvae that survived on the lab diet was significantly higher than the larval survival on a natural peach diet regardless of sterilization (NS: p= 0.001, S: p= 0.0079) (Fig. 1A, Table 1.1, Sup. Table 1.1). The undisturbed microbial community of the peach diet produced a significant positive effect on larvae survival, when compared with the autoclaved peach diet for both sterilized and non-sterilized larvae (NS: p= 0.0001, S: p< 0.0001) (Fig. 1A, Table 1.1, Sup. Table 1.1).

**Table 1.1:** The influence of diet on larvae phenotypes. Comparison of larvae life history traits and metabolic phenotypes between larvae, from all 10 genetic lines, raised on regular lab diet (R), peach diet (PR), and autoclaved peach diet (PA). NS stands for non-sterilized larvae, S stands for sterilized larvae. Asterisks indicate the significance of comparisons p< 0.001 ***, p< 0.01 **, and p< 0.5 *

| Treatment | Survival | Development | Weight | Triglyceride | Protein | Glucose |
| --- | --- | --- | --- | --- | --- | --- |
| NS | R> PR ** | R< PR *** | R> RP *** | R< PR *** | R> PR ** | R> PR *** |
| S | R> PR ** | R< PR *** | R> PR *** | R< PR *** | R> PR *** | R> PR *** |
| NS | PR> PA *** | PR< PA *** | PR> PA *** | PR< PA *** | PR< PA | PR< PA ** |
| S | PR> PA *** | PR< PA *** | PR> PA *** | PR< PA *** | PR< PA *** | PR< PA *** |

**Figure 1:**
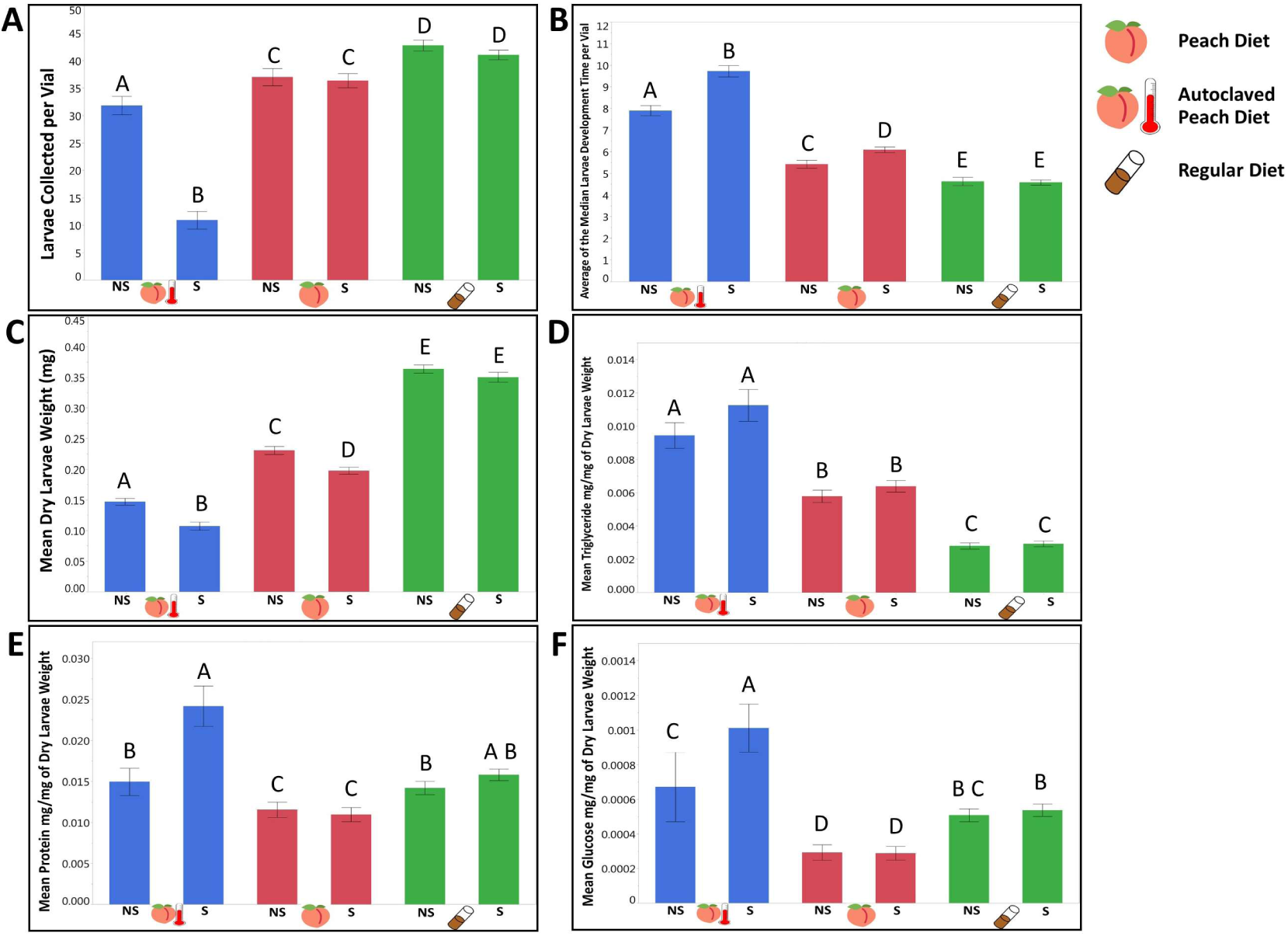
The Influence of diet and treatment on larvae mean. A) survival until late 3^rd^ instar stage B) development time C) weight D) Triglyceride E) Protein F) Glucose per mg of dry larvae weigh. Error bars indicated standard error.

**Development rate**. Within both controlled and sterilized treatments, larvae developed faster on the regular lab diet compared to the natural diet (NS: p= 0.0002, S: p< 0.0001) (Fig. 1B, Table 1.1, Sup. Table 1.1) and faster on the original peach food compared to the autoclaved diet (NS: p< 0.0001, S: p< 0.0001) (Fig. 1B, Table 1.1, Sup. Table 1.1). **Weight**. Larvae raised on the lab food were significantly heavier than those that were raised on the natural diet (NS and S: p< 0.0001) (Fig. 1C, Table 1.1, Sup. Table 1.1). Among the peach diets, larvae consuming the autoclaved diet were significantly lighter (NS and S: p< 0.0001) (Fig. 1C, Table 1.1, Sup. Table 1.1), suggesting that under natural nutritional conditions, microbiota in the food substrate facilitate growth and weight gain of the larvae. **Triglyceride**. Although fresh and incubated R food had higher triglyceride levels than the PR food (both p< 0.0001) (Sup. Table 1.2, 1.3), larvae raised on the PR diet had significantly higher triglyceride concentrations by weight than those that were raised on a lab diet, independent of sterilization treatment (NS and S: p< 0.0001) (Fig. 1D, Table 1.1, Sup. Table 1.1). Incubated autoclaved peach food had significantly higher triglyceride content (p< 0.0001) comparing with regular peach food (Sup. Table 1.3) and produced larvae with higher triglyceride by weight levels (NS and S p< 0.0001) (Fig. 1D, Table 1.1, Sup. Table 1.1). **Protein**. Independent of the treatment, larvae raised on the regular food had higher protein by weight levels compared to larvae raised on peach food (NS: p< 0.0045, S: p< 0.0001) (Fig. 1E, Table 1.1, Sup. Table 1.1). Larvae raised on the PA diet had significantly higher protein by weight levels compared to PR raised larvae, but only in the absence of parental microbiota (NS: p= 0.099, S: p< 0.0001) (Fig. 1E, Table 1, Sup. Table 1.1). Evaluating the difference in fresh food protein content, we found significantly higher protein concentration in the regular lab diet compared to the peach diet (p< 0.0001) (Sup. Table 1.2). Between fresh peach diets there was no significant difference (Sup. Table 1.2). However, after incubation, autoclaved peach food had significantly less protein than regular peach food (p= 0.0389) (Sup. Table 1.3). **Glucose**. Larvae raised on a lab food diet had significantly higher glucose levels than larvae raised on the peach food diet (NS: p< 0.0001, S: p< 0.0001) (Fig. 1F, Table 1, Sup. Table 1.1). Larvae that consumed PR food had lower glucose by weight levels compared with larvae raised on the autoclaved version, which was consistent with our 1.1 hypothesis (NS: p= 0.0089, S: p< 0.0001) (Fig. 1F, Table 1, Sup. Table 1.1). Interestingly, the fresh peach food had a significantly higher glucose concentration than the lab food (p< 0.0001) (Sup. Table 1.2). However, after incubating the peach food, the concentration of glucose was lower in PR than in both R (p<0.0001) and PA (p< 0.0001) diets (Sup. Table 1.3), suggesting a strong impact from the live microbial community.

**1.2** We observed that the presence of a maternally transmitted microbiota significantly impacted larvae phenotypes, and that impact varied across dietary treatments. **Survival**. The parental microbiota did not produce a significant effect on the larvae’s survival on the lab diet or the non-autoclaved peach diet (Fig. 1A, Table 1.2, Sup. Table 1.4), but its presence significantly enhanced the overall survival of larvae on the autoclaved peach diet (p< 0.0001) (Fig. 1A, Table 1.2, Sup. Table 1.4). This suggested the existence of microbial taxa that were necessary for a successful transition of the larvae through the instar stages under natural nutritional conditions, and that these microbial species could be picked up from the food substrate if available and/or inherited maternally. **Development rate**. Presence of parental microbiota on the peach diet reduced the number of days necessary for larvae to reach the third instar stage (p= 0.0002) and autoclaved peach diet (p< 0.0001) but not on a regular lab diet (Fig. 1B, Table 1.2, Sup. Table 1.4), suggesting that under natural nutritional conditions, maternal microbes might influence the developmental rate independent of the microbiota acquired from the food substrate. **Weight**. Maternally inherited microbiota produced a significant positive effect on larval weight on all of the tested diets (R: p= 0.005, PR: p= 0.0003, PA: p< 0.0001) (Fig. 1C, Table 1.2, Sup. Table 1.4). This indicated the universality of their influence on larval growth across food substrates. **Triglyceride**. Parental microbiota did not influence the triglyceride levels significantly on any diet (Fig. 1D, Table 1.2, Sup. Table 1.4). **Protein**. Evaluating the role of parental microbiota, we observed that sterilized larvae had higher protein by weight levels but only on the PA diet (p= 0.0014) (Fig. 1E, Table 1.2, Sup. Table 1.4). This suggested that the core microbiota involved in a natural metabolic phenotype formation might be inherited or acquired from the environment. **Glucose**. The parental microbiota reduced the glucose by weight levels only on PA food (p= 0.001) (Fig. 1F, Table 1.2, Sup. Table 1.4), indicating that both parental and environmental microbial taxa might be sufficient to reduce glucose levels in larvae.

**Table 1.2:**
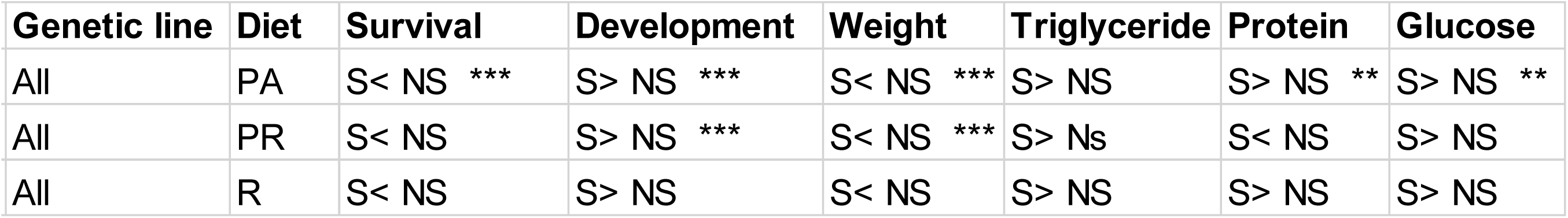
The influence of parental microbiota on larvae phenotypes. Comparison of larvae life history traits and metabolic phenotypes between larvae, from all 10 genetic lines, raised on regular lab diet (R), peach diet (PR), and autoclaved peach diet (PA). NS stands for non-sterilized larvae, S stands for sterilized larvae. Asterisks indicate the significance of comparisons p< 0.001 ***, p< 0.01 **, and p< 0.5 *

**1.3** Evaluating the contribution of tested independent variables on larvae phenotypes, we observed a genetic variation in most of the tested life history traits and phenotypes that interacted with the dietary conditions and the availability of maternally transmitted microbiota. **Survival**. All of the independent variables included in the model produced a significant effect on the larvae’s ability to survive until the late third instar stage (for diet, genetic line, treatment, diet by genotype, and the interactive effect of diet by genetic line p< 0.0001, genetic line by treatment p= 0.0374, and the interactive effect of diet, genetic line and treatment p< 0.0009). Out of the tested variables, diet was the strongest predictor of survival (variance explained (VE) 28.4%), followed by the interactive effect of the diet by treatment (VE= 8.02%) and genetic line (VE= 5.13%) (Table 1.3, Sup. Table 1.5). **Development**. The development rate of the larvae was significantly influenced by diet (p< 0.0001), genotype (p< 0.0001), and treatment (p< 0.0001) (Table 1.3). Among the specific interaction of these variables, only D*T (p< 0.0001) and G*T (p= 0.0374) produced a significant effect on development (Table 1). Diet was the key factor that influenced the time necessary for the larvae to reach the late third instar stage and explained almost half of all variance (VE= 47.5%) followed by the genetic line (VE= 4.98 %, Table 1). The combination of the rest of the variables was responsible only for 8.42 % of variation in developmental time (Table 1.3, Sup. Table 1.5).

**Table 1.3:** The contribution of diet, genotype, treatment, and their interactive effects on formation of larvae life history traits and metabolic phenotypes. VE stands for variance explained, by each independent variable. Asterisks indicate the significance of comparisons p< 0.001 ***, p< 0.01 **, and p< 0.5 *

| Independent variable | Survival | Development | Weight | Triglyceride | Protein | Glucose |
| --- | --- | --- | --- | --- | --- | --- |
| **Diet** | VE=28.4% *** | VE= 47.5% *** | VE= 31.4% *** | VE= 41.4% *** | VE= 4.24% *** | VE= 17.8% *** |
| **Genetic line** | VE=5.13% *** | VE=4.98% *** | VE=7.44% *** | VE=2.36% ** | VE=5.71% *** | VE=3.15% |
| **Treatment** | VE=4.69% *** | VE=2.41% *** | VE=0.74% *** | VE= 0.36% | VE=0.40% * | VE= 2.61% *** |
| **Diet\*Genetic line** | VE=3.13% *** | VE=0.96% | VE=2.88% *** | VE=5.03% *** | VE=5.09% *** | VE=5.52% |
| **Diet\*Treatment** | VE=8.02% *** | VE=1.41% *** | VE=0.10% | VE=0.07% | VE=0.45% | VE= 3.55% *** |
| **Genetic line\*Treatment** | VE=0.98% * | VE=2.82% *** | VE=1.06% *** | VE= 1.81% * | VE=3.53% *** | VE=0.72% |
| **Diet\*Treatment\*Genetic line** | VE= 2.37% *** | VE=0.82% | VE=2.75% *** | VE=4.81% *** | VE= 4.92% *** | VE= 2.37% |

**Weight**. All of the independent variables, with the exception of D*T, produced a significant effect on dry larval weight (p< 0.0001) with diet being the best predictor (VE= 31.4%), followed by genotype (VE= 7.44%), and D*G interaction (VE= 2.88%) (Table 1.3, Sup. Table 1.5). **Triglyceride**. Once again, diet explained the largest portion of variance (VE= 41.4%) across all independent variables. The interactive effect of D*G was a better predictor of triglyceride levels than the genotype (VE= 5.03% and 2.36%, respectively) (Table 1.3, Sup. Table 1.5). **Protein**. In contrast with other measured phenotypes, the variance explained by the model was predominantly evenly distributed across the independent variables, with genotype having the highest predicting power (VE= 5.71%) followed by D*G interactive effect (VE= 5.09%) (Table 1.3, Sup. Table 1.5). **Glucose**. Diet was the strongest predictor of larvae glucose levels (VE= 17.8 %, p< 0.0001) (Table 1.3, Sup. Table 1.5). Other variables that produced a significant effect on glucose levels were treatment (p= 0.0009), D*G (p = 0.05), and D*T (p= 0.0005) (Table 1.3, Sup. Table 1.5).

### 2) The symbiotic microbiota community composition of the larvae raised on the natural diet was different from the lab food raised larvae and was influenced by maternally inherited microbiota and the host’s genotype

**2.1** The gut microbial community composition and diversity varied substantially across dietary and treatment conditions. **Alpha diversity**. We characterized a total of 6,763 unique ZOTUs across the whole dataset with the number of ZOTUs per sample ranging from 55 to 886. The total number of reads per sample ranged from 4,685 to 908,308. Of the ZOTUs that could be assigned a taxonomic classification, we identified 134 classes, 218 families, and 394 genera represented across the samples. The response of alpha diversity to changing diets varied with the larval sterilization treatment. For NS larvae, we found that the Shannon index of larvae raised on PR and R diets was significantly higher than those raised on the PA diet (p= 0.0478 and p= 0.0252) (Sup. Table 2.1). All other comparisons were not significant. However, if the embryos were subjected to sterilization, the microbial species richness of larvae raised on the PA diet was significantly higher than larvae raised on the PR diet (p= 0.0371) (Sup. Table 2.1). In addition, there was no significant difference in microbial species richness between larvae raised on regular or peach regular diets (Sup. Table 2.1). We observed the exact same pattern for the PD whole tree index. Larvae raised on any diet were not significantly different in Shannon’s index (Sup. Table 2.1).

**Beta diversity**. The hierarchical clustering of the Bray-Curtis distances indicated that the most distant microbial communities were formed between larvae raised on the R and PR diet (Fig. 2A). This pattern held true for both sterilized and non-sterilized larvae (Fig. 2A). Clustering Weighted Unifrac distances suggested that PR and R diets may produce symbiotic microbial communities that were phylogenetically distant from each other, especially if the parental microbiota had been removed (Fig. 2B).

**Figure 2:**
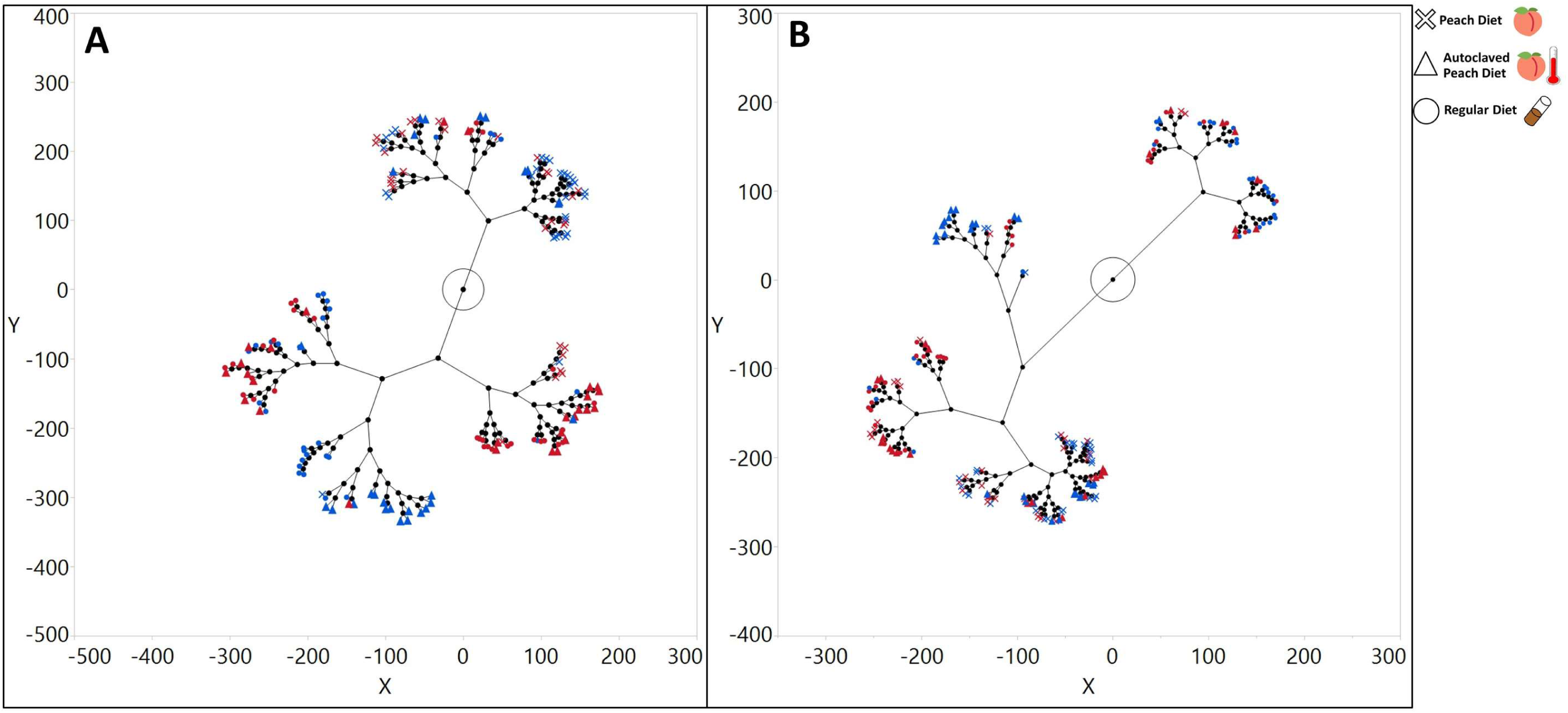
The influence of the diet on beta diversity distances between larvae symbiotic microbiota communities. Constellation plot based on hierarchal clustering of larvae microbiota community beta diversity distances A) Bray-Curtis distances B) Weighed Unifrac distances. Samples from sterilized larvae are marked with a blue color and samples from non-sterilized larvae are marked with the red color. Samples raised on regular lab diet are marked with circled, on a peach diet with crosses and on peach autoclaved diet with triangles.

**Taxa composition**. Applying discriminant analysis on the ten most abundant microorganisms at each taxonomic level revealed which organisms were largely responsible for the differentiation of the microbial composition on the canonical plot, based on diet. Thus, PR food is largely defined by the abundance of Cyanobacteria at the phylum level (Sup. Fig. 1), Epsilonproteobacteria at the class level (Sup. Fig. 2), Streptophyta at the order level (Sup. Fig. 3A, 3B), Leuconostocaceae sequences at the family level (Sup. Fig. 4), and *Leuconostoc* at the genera level (Fig. 3A, Sup. Fig. 5). In turn, the lab diet was defined by Firmicutes, Bacilli, Lactobacillales, Lactobacillaceae, and *Lactobacillus* respectively (Sup. Fig. 1-5, Fig. 3A). Interestingly, when we considered only the 10 most abundant organisms at each taxonomic level, we did not see a full separation between R and PA diets unless the larvae were sterilized (Sup. Fig. 1-5). If parental microbiota were removed, the differentiation of the PA diet was led by Actinobacteria, Actinobacteria and Alphaproteobacteria, Clostridiales and Rickettsiales, Lachnospiraceae and Rickettsiaceae, and *Bacteroides* and *Wolbachia* (Sup. Fig. 1B-5B, Fig. 3A). Including all identified bacterial groups in the discriminant analysis revealed that diet was a good predictor of bacterial taxa composition at phylum, class, order, family, genus, and even individual ZOTU levels (Fig 3B, Sup. Fig.6-11).

**Figure 3:**
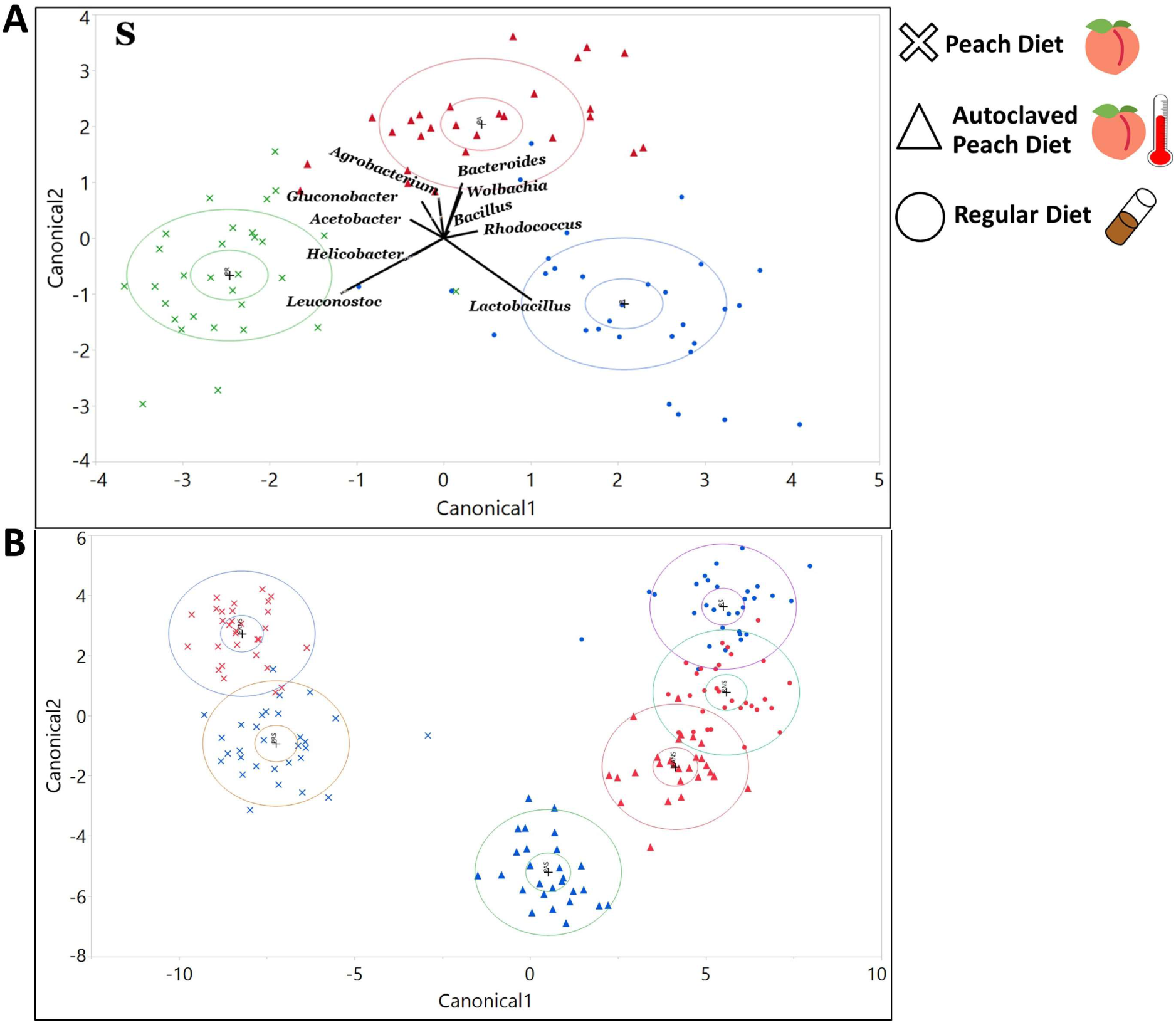
The influence of the diet on larvae symbiotic microbiota communities’ composition. Discriminant analysis of symbiotic microbiota community based on taxa relative abundances in larvae from non-sterilized (red) and sterilized (blue) treatments. A) 10 most abundant microbial genera found in sterilized larvae. The length of the vector is correlated with the strength of the impact that it produced for the samples to be separated, in the vector direction, on the canonical plot B) 100 most abundant microbial zotu found in both sterilized and non-sterilized larvae. Non sterilized samples are marked with red color and sterilized larvae are marked with blue color.

Overall, for non-sterilized larvae the abundance of eight phyla, 12 classes, 20 orders, 27 families, 40 genera, and 141 ZOTUs were significantly different between PR and R food (Sup. Tables 3.1-3.6). Comparing PR and PA food, we found that the abundance of four phyla, six classes, nine orders, 16 families, 20 genera, and 76 ZOTUs were significantly different (Sup. Tables 3.1-3.6). Lastly, we observed the significant difference for the abundance of four phyla, three classes, five orders and families, three genera, and 27 ZOTUs between R and PA food (Sup. Tables 3.1-3.6). This indicated the minimal difference between microbial communities of these diets to be consistent with the discriminant analysis.

In larvae lacking parental microbiota, we observed that the abundance of six phyla, 10 classes, 18 orders, 34 families, 37 genera, and 114 ZOTUs were significantly different between lab and peach diets (Sup. Tables 3.1-3.6). Comparing PR and PA diets, we found a significant difference in abundance of seven phyla, 14 classes, 32 orders, 48 families, 67 genera, and 200 ZOTUs (Sup. Tables 3.1-3.6). R and PA diets were significantly different in the abundance of five phyla, six classes, nine orders, 13 families, 18 genera, and 87 ZOTUs (Sup. Tables 3.1-3.6).

**2.2** The maternally transmitted microbiota influenced the composition of the larvae’s symbiotic microbiota communities. **Alpha diversity**. S larvae had higher values for species richness, Shannon, and PD whole tree indexes on the PA diet (Sup. Table 2.2). NS larvae had a significantly higher Simpson index value on the PR diet (Sup. Table 2.2). All other comparisons were not significantly different. **Beta diversity**. When comparing the difference between beta diversity metrics in NS and S treatments for each diet, we observed a distinctive clustering, based on the treatment of samples that were raised on PA food for Bray-Curtis Distance (Fig. 4A) and Weighted Unifrac distance (Fig. 4D). For the samples that were raised on R food, we observed the clustering for Weighted Unifrac distance only (Fig 4F).

**Figure 4:**
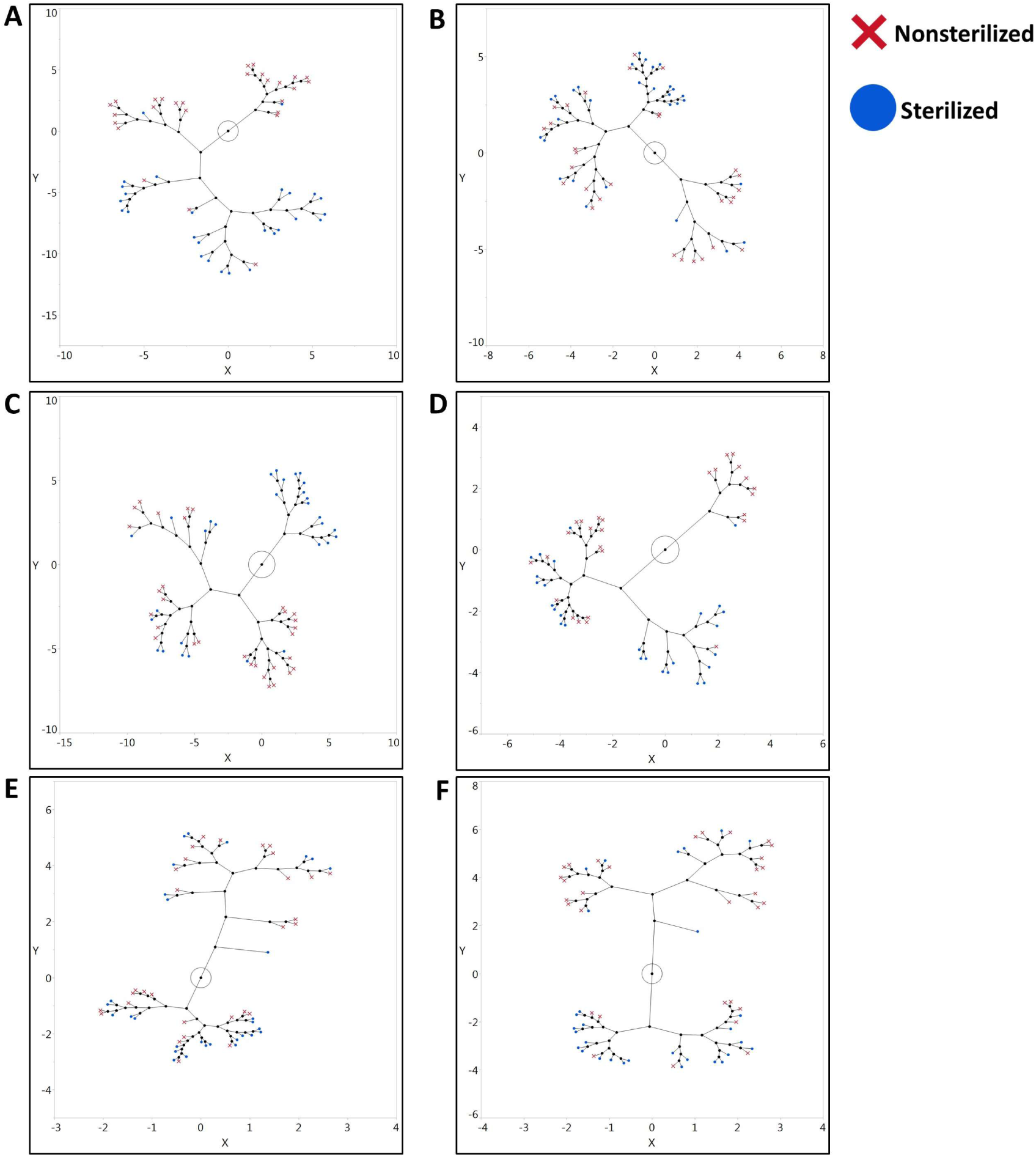
The influence of the treatment on beta diversity distances between larvae symbiotic microbiota communities. Hierarchal clustering of larvae microbiota community beta diversity distances A-C) Bray-Curtis distances between NS and S larvae raised on PA, PR, and R diets, respectively D-F) Weighed Unifrac distances between NS and S larvae raised on PA, PR, and R diets, respectively.

**Taxa composition**. The discriminant analysis indicated that the status of inheritance of the parental microbiota could serve as a good predictor for differentiation of the bacterial community as indicated with the canonical plot on all taxonomic levels (phylum, order, class, family, genus, and ZOTU) (Fig.5 A-C, Sup. Fig. 12A-17C). Among the 10 most abundant phyla that defined the differentiation of the NS community were Firmicutes and Bacterioidetes on the PA diet (Sup. Fig. 18A), Firmicutes on the PR diet (Sup. Fig. 18B), and Proteobacteria, Tenericutes, and Flusobacteria on the R diet (Sup. Fig. 18C). Phyla that were influential for differentiation of the S community were Actinobacteria, Flusobacteria and Cyanobacteria on the PA diet (Sup. Fig. 18A), Actinobacteria, Tenericutes, and Flusobacteria on the PR food (Sup. Fig. 18B), and Planctomycets and Bacterioides on the R diet (Sup. Fig. 18C). Considering bacterial classes, the NS treatment was strongly defined by Bacilli on the PA (Sup. Fig. 19A) and the PR diets (Sup. Fig. 19B), and Alphaproteobacteria and Bacteroida on the R food diet (Sup. Fig. 19C). The sterilization treatment was mostly separated due to Actinobacteria on the PA diet, Alpha and Beta proteobacteria on the PR diet, and Gammaproteobacteria on the R diet (Sup. Fig. 19A-C). On R food, microbial communities from the sterilized and non-sterilized larvae were not fully separated on the canonical plot (Sup. Fig. 19C). At the order level, the NS community was defined by Lactobcillales on the PA and the PR diets, and Actinomycetales and Rhodospirillales on the R food (Sup. Fig. 20A-C). Sterilized larvae were associated with abundances of Streptophyta on the PA food, as well as Burkholderiales and Rhodospirillales on the PR diet (Sup. Fig. 20A-B). On the R diet, four out of ten tested orders were strongly associated with the S treatment (Sup. Fig. 20C). At the family level, the NS larvae were correlated with Lactobacillaceae on the PA and the PR diets and Acetobacteraceae on the R food diet (Sup. Fig. 21A-C). Sterilized larvae were defined by the abundance of Nocardiaceae on the PA food and Leuconostocaceae and Caulobactereceae on the R diet (Sup. Fig. 21A-C). On the genera level, NS was primarily separated by *Lactobacillus* on the PA diet and *Acetobacter* and *Agrobacterium* on the R food (Sup. Fig. 22A, C). On the PR food diet, 95% confidence ellipses almost overlapped, indicating that sterilization status might not be the decisive predictor for abundance of the 10 most common genera (Fig. 5E). The S treatment was primarily defined by the abundance of *Leuconostoc* and *Gluconobacter* on the PA diet and *Lactobacillus* and *Leuconostoc* on the R diet (Fig. 5 A, C, Sup. Fig. 22A, C).

**Figure 5:**
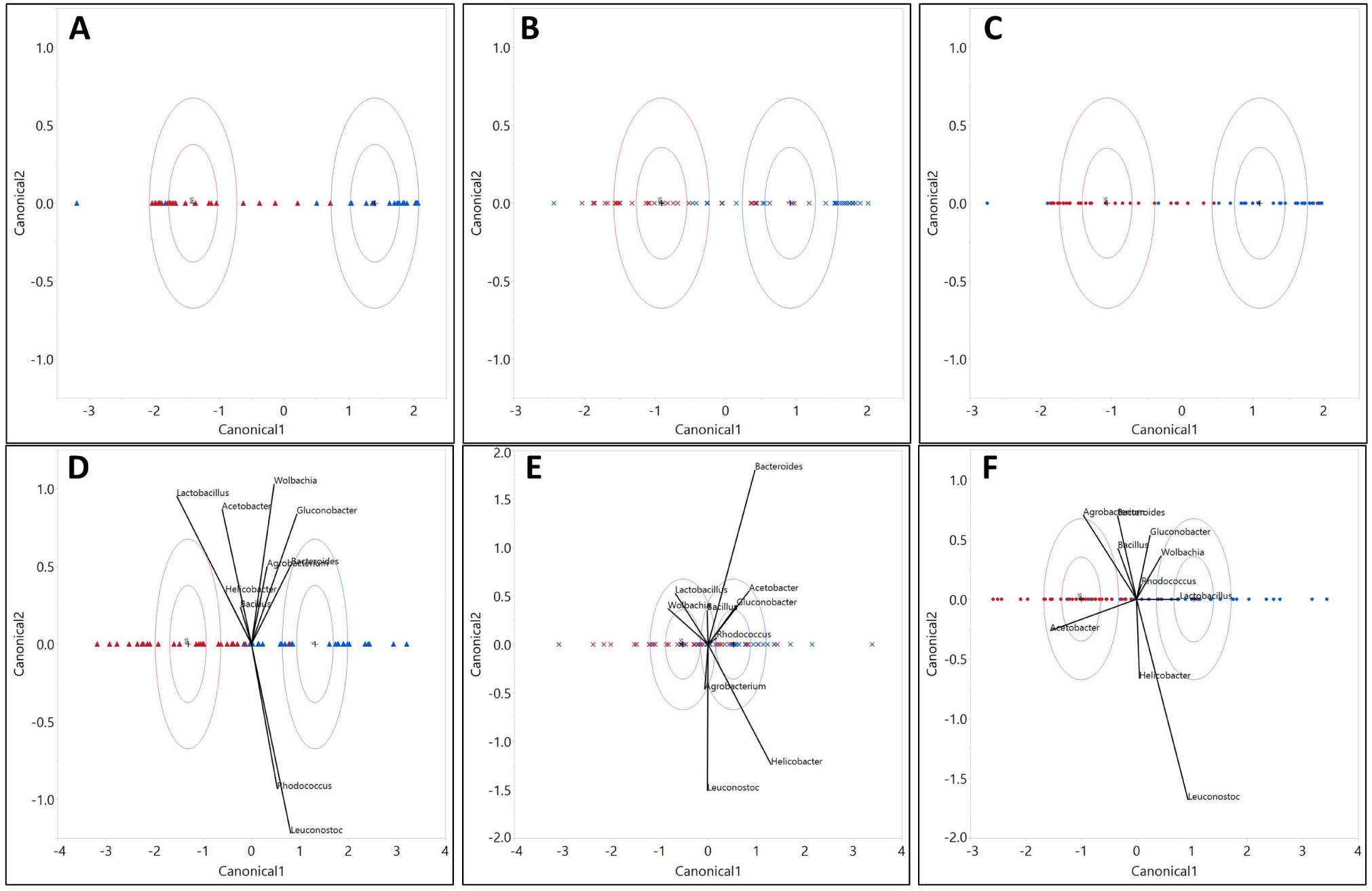
The influence of the treatment on larvae symbiotic microbiota communities’ composition. Discriminant analysis of symbiotic microbiota community based on taxa relative abundances in larvae from non-sterilized (red) and sterilized (blue) treatments. The analysis includes A) All identified ZOTUs on a PA diet B) All identified ZOTUs on a PR diet C) All identified ZOTUs on a R diet D) 10 dominant genera on a PA diet E) 10 dominant genera on a PR diet F) 10 dominant genera on a R diet. In figures D-F, the length of the vector is correlated with the strength of the impact that it produced for the samples to be separated in the vector direction, on a canonical plot.

Multiple microbial groups were significantly different in their distribution between NS and S treatments across the diets on all taxonomic levels. On regular food, there existed an abundance of four phyla (Sup. Table 4.1), four classes (Sup. Table 4.2), five orders (Sup. Table 4.3), six families (Sup. Table 4.4), nine genera (Sup. Table 4.5), and 41 ZOTUs (Sup. Table 4.6). On the PR diet we saw a significant difference in the abundance of one phylum (Sup. Table 4.1), two classes (Sup. Table 4.2), six orders (Sup. Table 4.3), nine families (Sup. Table 4.4), 11 genera (Sup. Table 4.5), and 61 ZOTUs (Sup. Table 4.6). The highest number of significantly different taxa was observed on the PA diet with seven phyla (Sup. Table 4.1), 12 classes (Sup. Table 4.2), 21 orders (Sup. Table 4.3), 34 families (Sup. Table 4.4), 43 genera (Sup. Table 4.5), and 148 ZOTUs (Sup. Table 4.6).

**2.3** The composition of the microbial community exhibited variation with host genotype, which further exhibited a significant interactive effect with diet and treatment. We also tested the influence of genotype and other variables’ interactive effect on the abundance of microbiota. At the phyla level, 14 were significantly influenced by genotype, three by D*G interaction, five by G*T, five by D*T, and six by D*G*T (Sup. Table 5.1). Abundances of 30 classes were significantly influenced by genotype, eight by D*G, G*T, and D*T, and 10 by D*G*T interaction (Sup. Table 5.2). Among the orders, an abundance of 46 was significantly influenced by genotype, 13 by D*G, 15 by G*T, 16 by D*T, and 15 by D*G*T (Sup. Table 5.3). The abundance of 72 families was significantly influenced by genotype, 20 by D*G, 15 by G*T, 18 by D*T, and 23 by D*G*T (Sup. Table 5.4). Lastly, genotype significantly influenced 94 genera, D*G influenced 44, G*T influenced 46, D*T influenced 30, and D*G*T influenced 45 genera (Sup. Table 5.5).

**3.1** We identified microbial taxa that exhibited correlations with host phenotype across diets and treatments, with many that had a diet, treatment or genotype specific relationship. Across the NS larvae, we found four significant interactions on the R food, 14 on PR diet, and 14 on PA diet at the phylum level (Sup. Table 6.1). For the S larvae at the same taxonomic level, we found eight significant interactions on the R diet, five on the PR and two on PA food (Sup. Table 6.1). At the class taxonomic level, nine significant interactions were found on R food, 35 on PR, and 24 on PA diet (Sup. Table 6.2). Considering S larvae, there were 23 significant correlations on the R diet, 11 on PR, and four on the PA diet (Sup. Table 6.2). For NS larvae at the order level, we found 23 significant correlations on the R diet, 57 on PR, and 46 on PA diet (Sup. Table 6.3). For S larvae we found 29 significant correlations on R food, 22 on PR, and nine on PA diet (Sup. Table 6.3). At the family level, we found 34 significant correlations on R food, 88 on PR, and 67 on PA diet for NS larvae (Sup. Table 6.4). For S larvae, we observed 51 significant interactions on R food, 29 on PR, and 14 on PA diets (Sup. Table 6.4). Across the genera, we found 40 significant interactions on R, 105 on PR, and 64 on PA diets, for NS larvae (Fig. 6-7, Sup. Table 6.5). Considering S larvae, we found 76 significant interactions between tested taxa and phenotypes on R, 46 on PR, and 33 on PA diets (Fig. 6-7, Sup. Table 6.5). At the level of individual ZOTUs, for NS larvae, we found 226 significant interactions on R, 283 on PR, and 225 on PA diets (Sup. Table 6.6). For S larvae the number of significant interactions between ZOTUs abundances and larvae phenotypes were as follow 313 on R, 164 on PR, and 230 on PA diets (Sup. Table 6.6).

**Figure 6:**
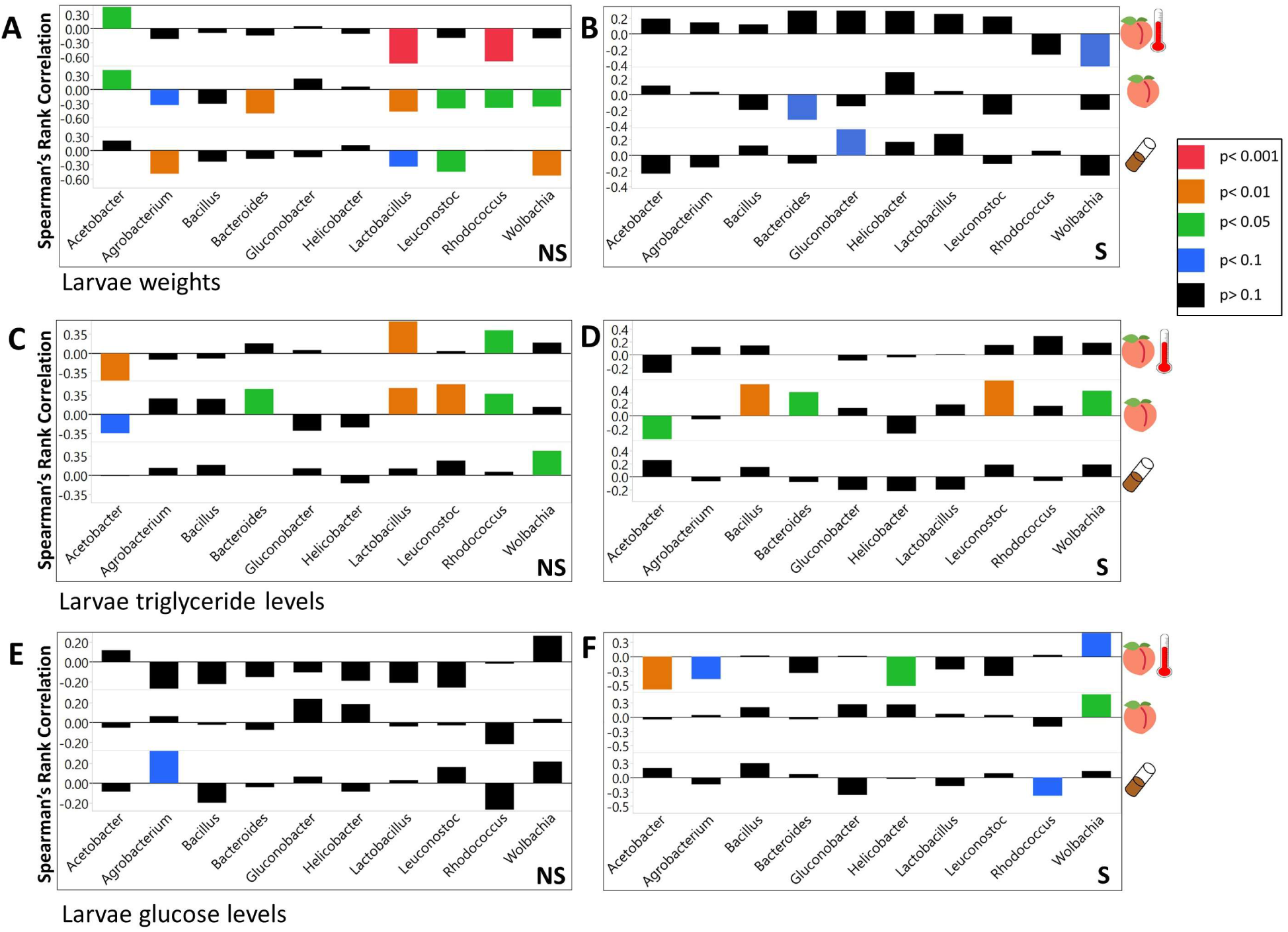
The influence of symbiotic microbial taxa on larvae metabolic phenotypes. Spearman’s rank correlation coefficients between the abundance of 10 dominant symbiotic microbiota genera and larvae phenotypes, on each diet. The color of the bars corresponds to the level of significance for each correlation. A) Weight of NS larvae B) Weight of S larvae C) Triglyceride levels of NS larvae D) Triglyceride levels of S larvae E) Glucose levels of NS larvae F) Glucose levels of S larvae

**Figure 7:**
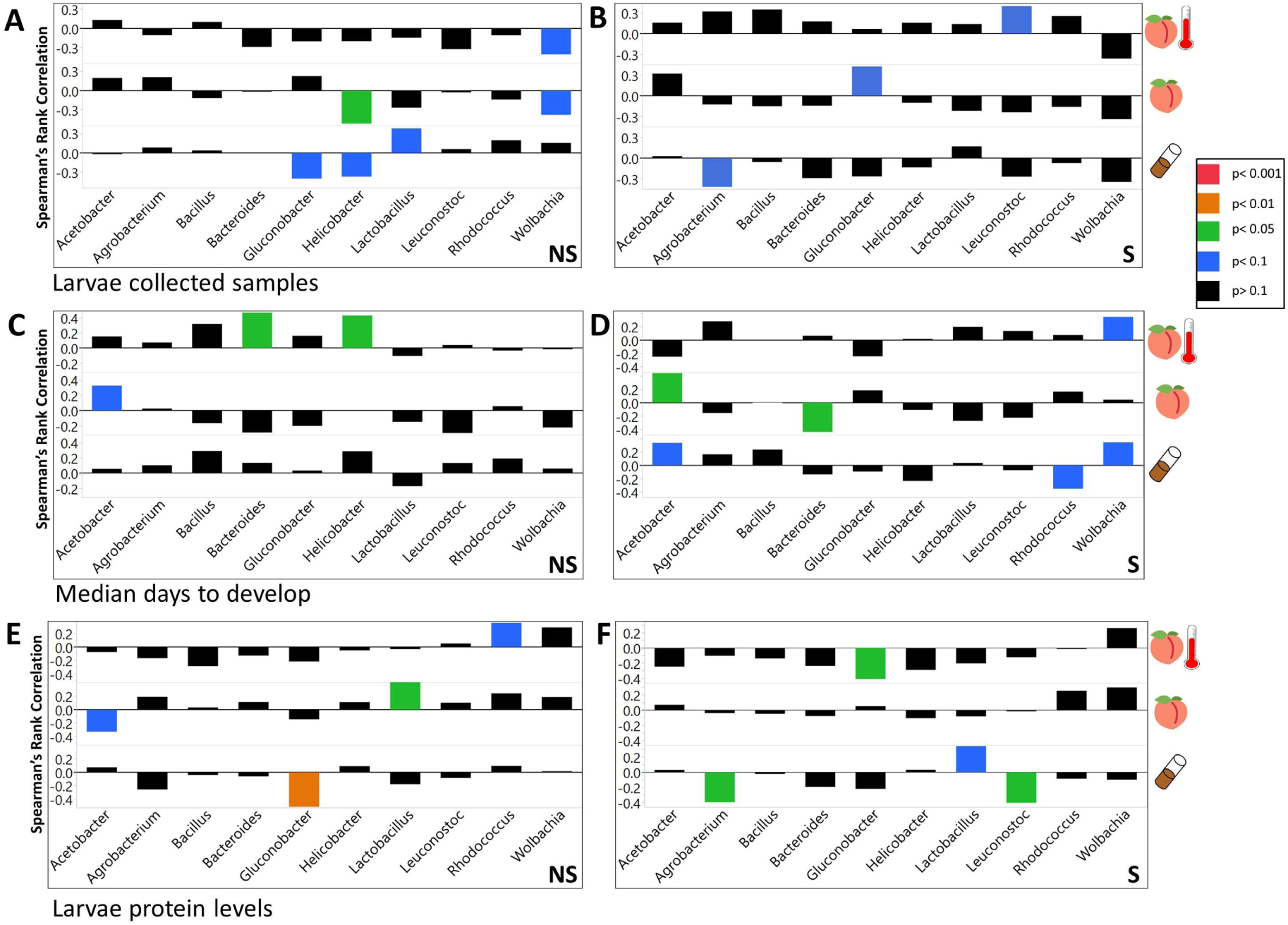
The influence of symbiotic microbial taxa on larvae life history traits and metabolic phenotypes. Spearman’s rank correlation coefficients between the abundance of 10 dominant symbiotic microbiota genera and larvae phenotypes on each diet. The color of the bars corresponds to the level of significance for each correlation. A) Total number of collected NS larvae B) Total number of collected S larvae C) Median number of days to reach pre-pupation stage for NS larvae D) Median number of days to reach pre-pupation stage for S larvae E) Protein levels of NS larvae F) Protein levels of S larvae

We evaluated the interactive effect of the abundance of microbial taxa and diet, genotype, and treatment on forming the tested phenotypes. D*A produced a significant effect in 11 cases at the phylum level (Sup. Table 7.01), in 26 at the class level (Sup. Table 7.02), in 52 at the order level (Sup. Table 7.03), in 72 at the family level (Sup. Table 7.04), and in 95 cases at the genus level (Sup. Table 7.05). We found a significant G*A interaction in eight cases at the phylum level (Sup. Table 7.06), in 10 at the class level (Sup. Table 7.07), in 39 at the order level (Sup. Table 7.08), in 57 at the family level (Sup. Table 7.09), and in 105 cases at the genus level (Sup. Table 7.10). T*A produced a significant effect in 13 cases at the phylum level (Sup. Table 7.11), in 27 at class level (Sup. Table 7.12), in 36 at the order level (Sup. Table 7.13), in 60 at the family level (Sup. Table 7.14), and in 87 cases at the genus level (Sup. Table 7.15).

## Discussion

### 1.1: Overall, it appeared that frozen peach food was capable of providing nutritional conditions similar to the natural ones and can preserve key microbial taxa necessary for survival and development of *Drosophila* larvae

The reduction in survival, increase in development time and increase in triglyceride levels, as well as decreased weight and protein levels of the larvae raised on the natural food compared with the larvae raised on the R lab food resembles the phenotype generated by a reduced protein diet. These findings correlate with our evaluation of the protein concentrations in different diets. (Klepsatel et al., 2018, Bing et al., 2018, Skorupa et al., 2008, Sang, 1956). In addition, the adaptation of *Drosophila* to the lab environment was connected to increased weight and reduced stress tolerance (Sgro and Partridge, 2000, Hoffmann et al., 2001, Russell et al., 2012). Therefore, nutritional and pathogenic stresses associated with the natural food conditions could further contribute to the decrease in survival and development rate of larvae raised on the PR food compared to the standard diet (Staubach et al., 2013, Bing et al., 2018, Pais et al., 2018, Sang, 1956).

The pattern regarding glucose concentration was more interesting. Freshly unfrozen peach food had a higher glucose concentration, but larvae raised on the PR diet had the lowest concentration compared to larvae raised on any other diet. This pattern was likely caused by the activity of naturally acquired microbes since it was shown that the presence of several microbial taxa that naturally associate with *Drosophila*, such as *Acetobacter*, is correlated with decreased sugars in fly food and *Drosophila* itself (Huang and Douglas, 2015, Dobson et al., 2015). In addition, incubation of the PR food, even without the larvae, led to a drastic reduction of glucose levels compared to the R and PA diets. Furthermore, the difference between all phenotypes (with the exception of glucose) increased if the peach food diet was autoclaved and even more (with the exception of triglyceride) if the parental microbiota were not transferred to the autoclaved diet. This suggested that symbiotic microbiota drove the phenotypic change between PR and PA raised larvae.

These findings are consistent with previous studies which showed that the presence of naturally associated microbiota was advantageous for *Drosophila melanogaster* and *Drosophila suzukii* on fresh fruit diets, which are also poor in protein content (Pais et al., 2018, Bing et al., 2018). In fact, larvae raised on a PA diet closely resembled the phenotype of axenic larvae and axenic larvae raised under low protein nutritional conditions. Examples of this similar phenotype include lower survival and body size/weight (Shin et al., 2011, Wong et al., 2015, Dobson et al., 2015), longer development time (Shin et al., 2011, Newell and Douglas, 2014a), elevated glucose (Huang and Douglas, 2015) and triglyceride levels (Newell and Douglas, 2014a, Dobson et al., 2015).

### 1.2 Maternally deposited microbes produced positive effects on larvae that were raised on the peach diets

Interestingly, the presence of parental microbiota did not produce a significant effect on any of the tested phenotypes, when larvae were raised on the lab diet. Contrarily, on the peach diet, the presence of parental microbota increased the weight and development rate even if the original peach microbiota were still present. These findings are consistent with the reports of beneficial effects, of the maternally deposited microbiota, for larvae on a fruit diet. These results also indicate the importance of considering an organism’s natural environmental conditions when addressing the questions about symbiotic relationships and evolutionary patterns (Pais et al., 2018, Bing et al., 2018).

### 1.3 Genotype was one of the key factors that influenced larvae phenotypes

It is important to note that although the described patterns were observed for the total experimental population of larvae, the genetic component still played a significant role in generating all but the glucose phenotype. In addition, consistent with previous research, we observed that D*G interaction played a significant role in forming metabolic phenotype as well as contributed to the survival of the organism (Reed et al., 2010, Reed et al., 2014). Furthermore, most of the tested phenotypes were significantly correlated with G*T and even D*G*T, indicating the importance of considering multiple factors to understand the development of complex traits.

### 2.1 Microbiota of the larvae raised on PR food exhibit a distinct community structure and might remind microbiota of wild flies

Multiple studies were performed to evaluate the gut microbiota composition of lab and wild populations of *Drosophila* (Chandler et al., 2011, Adair et al., 2018, Douglas, 2018, Pais et al., 2018, Wong et al., 2013). Although most of them consistently report the prevalence of different members of Alphaproteobacteria, Bacilli, or Gammaproteobacteria in lab and wild populations, the relative abundance of the taxa, especially at lower taxonomic levels, often varies between studies (Douglas, 2018). In our work, larvae raised on the PR food diet formed a distinct community clearly separated from the larvae raised on the R food diet, as displayed on the canonical plot. We observed a higher prevalence of *Gluconobacter* and *Leuconostoc* and lower abundance of *Lactobacillus* in larvae raised on the PR diet compared with the R food diet (Pais et al., 2018, Staubach et al., 2013, Corby-Harris et al., 2007, Chandler et al., 2011), which is consistent with previous findings performed on natural populations of *Drosophila*.

However, it is difficult to judge how well the microbial community of our experimental larvae represent the microbial community of wild population, as the variety of factors could influence gut microbiota composition in flies which certainly complicates the comparison between studies (Douglas, 2018, Staubach et al., 2013, Jehrke et al., 2018). As such, it was shown that the gut microbiota composition of lab reared flies may vary with diet (and even among the standard diets with the major carbohydrate source), genetic line, development stage temperature, and etc. (Jehrke et al., 2018, Douglas, 2018, Wong et al., 2011, Moghadam et al., 2018). The wild populations of gut microbiota in *Drosophila* was shown to vary with collection location and diet (Adair et al., 2018, Wong et al., 2013, Staubach et al., 2013, Martinez-Porchas et al., 2017). In other insects and wild populations of vertebrates, gut microbiota was shown to change even with seasonality (Behar et al., 2008, Ferguson et al., 2018, Tong and Zhang, 2019, Maurice et al., 2015).

In addition, the relationship between *Drosophila* gut microbiota during the developmental and adult stages is a subject of controversy between a few studies that compared those relationships (Jehrke et al., 2018, Wong et al., 2011, Vacchini et al., 2017). Furthermore, to the best of our knowledge, the gut microbiota of the larvae from the natural populations was not assessed at all. This is likely due to the complexity of identifying *Drosophila* species during the larval stage.

Therefore, we hope to provide the methodology for the possibility of exploring the effects of a natural diet, and the microbial community associated with it, in a controlled lab environment. This setting provides the opportunity to work not only with adult flies but also with larvae.

### 2.2-2.3 Community structure of symbiotic microbiota were correlated with diet, treatment, host genotype and their specific interactive effects

The development of symbiotic microbiota populations was shown to be correlated with the available nutrients present in the diet, the host’s genotype and parental microbiota left on the chorion of the egg (Douglas, 2018, Jehrke et al., 2018, Wong et al., 2015). Complementary to the results reported by Jehkre (2018), we also observed that the genotype of the host may influence the abundance of bacterial taxa more than the diet. Wong (2015) reported that the bacterial population deposited on the *Drosophila* embryo may shift the symbiotic microbiota population of the offspring, even in the presence of bacteria that previously colonized the food substrate. We observed similar results in most cases.

However, among the 10 most abundant genera on the PR diet, the full separation of the S and NS larvae microbial community compositions was not present on the canonical plot indicating the possibility of a difference in the response of the lab and the natural microbial population to the presence of *Drosophila* parental microbiota. This differentiation was not likely caused by the nutrition composition of the food since the PA separation, represented on the canonical plot, between S and NS treatments was obvious in all cases. In addition, for beta diversity distances, the abundance of individual microbial taxa, as well as the correlations between the abundances of microbial taxa within the microbial community, represented patterns found in PA food that resembled the ones in the R food raised larvae, if parental microbiota were not removed (Fig 4, 5, Sup. Fig.12-23). Overall, consistently with previous studies, our findings indicated the dependency of relative microbial abundances on all of the tested variables and additionally on the interactive effect between them (Wong et al., 2015, Jehrke et al., 2018, Douglas, 2018).

### 3.1 The influence of individual microbial taxa as well as the influence of the whole microbial community on the host may vary with the diet and other environmental and genetic conditions

Genotype and gut microbiota composition are among the major factors that control the development of obesity traits (Parks et al., 2013). Changes in some key microbiota populations are associated with the rapid expansion in the prevalence of metabolic syndrome (Zhang et al., 2010). Alterations in the gut microbiota community can modulate insulin secretion and sensitivity, thus contributing to diabetes susceptibility (Kreznar et al., 2017). Moreover, previous research indicates that genetic variation considerably influences the gut microbiota composition (Zhang et al., 2010, Kreznar et al., 2017, Jehrke et al., 2018). However, most of the studies mentioned above have used less than ten genotypes to study the correlation between gut microbiota and the pathogenesis of obesity in mice. The challenges of using a mouse model involve relatively high expenses for husbandry and logistics (Rosenthal and Brown, 2007, Berger et al., 2005, Martin et al., 2016).

*Drosophila melanogaster* is an exceptional model to study the effect of genotype on the phenotype formation, due to the variety of established tools such as Drosophila Genetic Reference Panel and The Drosophila Synthetic Population Resource. These resources offer a variety of diverse genotypes, with sequenced parental genomes, that allow for testing the microbiota effects across various genetic backgrounds and provides potential for studying genetic interaction between host and its symbionts, and even mapping the specific genetic loci responsible for the interactions (Mackay et al., 2012, King et al., 2012, Chaston et al., 2016). The phenotypic response to a diet modification often varies with the genotype (Reed et al., 2010, Reed et al., 2014). In fact, diet by genotype (DxG) interaction may explain more variance than diet alone in the metabolic response of such traits, such as triglyceride and carbohydrate concentrations (Reed et al., 2010). In addition, recent findings showed that genotype by diet interactions significantly influences metabolomic profiles; hence, laying the foundation for explaining the mechanism through which DxG influences metabolic traits (Reed et al., 2014, Williams et al., 2015). Our results are consistent with previous research, in that phenotypic response varied significantly between genetic lines (Reed et al., 2014, Williams et al., 2015). Genotype had a significant effect on survival, development rate, and triglyceride levels, and was the second-best predictor of weight and the best predictor of protein levels. Obesity and type two diabetes are associated with elevated weight, high blood glucose concentrations, and excess accumulation of adipose tissue (Martyn et al., 2008, Akter et al., 2017). Consistent with recent studies linking *Lactobacillus* and *Coprococcus* to obesity in humans, our results show that these genera are positively associated with glucose levels (Million et al., 2012, Ignacio et al., 2016, Murugesan et al., 2015, Armougom et al., 2009). In addition, previous research has shown an overall decrease in the abundance of *Firmicutes* in obese humans (Schwiertz et al., 2010). Similarly, we observed that the total abundance of *Firmicutes* is negatively associated with triglyceride levels. It should be noted that the correlation between metabolic phenotype and particular microbial taxa could vary between studies (Ley et al., 2006, Furet et al., 2010, Turnbaugh et al., 2006).

Consistently with previous *Drosophila* research, we observed that the abundance of microbial taxa was correlated with measured phenotypes. Such as *Acetobacteraceae* is negatively correlated with larvae glucose levels (Chaston et al., 2016, Douglas, 2018, Chaston et al., 2014). Additionally, *Acetobacter* increases development time while *Lactobacillus* and *Firmicutes* decreases it (Chaston et al., 2014, Chaston et al., 2016, Newell and Douglas, 2014a, Storelli et al., 2011). Previous work showed that *Acetobacter* species reduced triglyceride levels while most *Lactobacillus* species had no effect (Chaston et al., 2016, Newell and Douglas, 2014a). In contrast, our data shows that *Acetobacter* did not significantly affect triglyceride levels, and *Lactobacillus* showed a negative correlation. Consistent with Newell and Douglas (2014b), we found that *L. brevis* and *L. plantarum* had no significant effect on protein levels, but in addition, our results indicated that the abundance *Acetobacter* was negatively correlated with the protein levels.

Consistently with Jehrke (2018), we observed that most of the correlations between the tested phenotype and abundance of microbiota are relatively weak. Weaker correlations observed with large sample sizes in microbiome research, while significant, fail to hold up to the use of stricter FDR values or other conservative p adjustment methods (Jehrke et al., 2018, Wu et al., 2011). Expanding the analysis of the bacterial species abundance for each phenotype beyond the most dominant species, while providing a more complete overview of the correlation between tested phenotypes and microbial abundance, also raises FDR values as a result of increasing sample sizes (Wu et al., 2011). Previous microbiome studies have dealt with high FDR values by accepting higher thresholds, so as to not miss possible correlations (Wu et al., 2011). Since the level of FDR that should be tolerated is poorly defined and often widely variable compared to accepted p-values, its value is often seen as arguable (Pawitan et al., 2005). Considering the large sample sizes used in our analysis, using a low FDR value may obscure important correlations between the tested phenotypes and their abundance of microbiota (Wu et al., 2011).

Some of the inconsistencies between our work and previous studies on the correlation of the abundance of microbial taxa and measured metabolic phenotype, perhaps may be addressed to the interactive effect between the variables included in the experiments. Several studies showed that the contribution of the symbiotic microbiota to the host may be observed only in a diet dependent manner. Such as Shin et al. (2011), showed that axenic *Drosophila* larvae would not be able to develop on a protein poor diet without activation of the insulin signaling pathway by its symbiotic microbe. Wong et al. (2014) found diet-dependent differences in microbiota produced effect, including differences in vitamin microbial sparing on a low-yeast diet and suppression of lipid and carbohydrate storage on a high-sugar diet. Bing et. al (2018) found that symbiotic microbiota of *D. suzukii* are critical for providing proteins for development of flies raised on fresh fruit, but that these microbial proteins are not essential for development of flies raised on a nutrient sufficient diet.

In our study, we also observed that the effect that microbial abundance at the level of individual taxa produced on larvae phenotype varied with the diet. In few cases even the direction of the correlation between the abundance of microbial taxa and tested phenotype was opposite on different diets. In addition, using PCA, we observed that correlational effects that microbial abundance (as an example at the family level) produced on measured metabolic (Sup. Fig. 24A) and fitness phenotypes (development rate and survival) (Sup. Fig 24B) varied between the diets. The correlation coefficients for the influence of all microbial taxa on measured metabolic phenotypes clustered together for the PR diet but not for other diets (Sup. Fig 24A). The correlation coefficients between microbiota abundances and fitness phenotypes clustered for all but PA diets (Sup. Fig 24B). Taking into account all of the described findings, perhaps in future studies that will aim to understand the mechanisms for formation of metabolic phenotypes, we will see a consistent control for the gut microbiota composition in a similar fashion as now we can see it for genetic lines and nutritional composition of the food.

## Acknowledgements

We appreciate the research assistance of L. Griffin, C. Hart, V. Oza, C. Scott, R. O’Rourke K. MacIntyre, K. Lowman C. Tunckanat, J. Jarnigan, Y. Nam, and all of the members of the Reed lab. Helpful guidance was provided by J. Yoder, J. Olson, S. Chtarbanova-Rudloff, C. Morrow, and J. Lopez-Bautista. Special thanks for research assistance, guidance, and moral support to S. Yan. Funding sources included University of Alabama Graduate School, Department of Biological Sciences, Graduate Student Association, University of Alabama at Birmingham Research Voucher Program, National Institutes of Health: 5R01GMO98856.

