## Supplemental figures for "Influence of the lab adopted natural diet and environmental and parental microbiota on life history and metabolic phenotype of *Drosophila melanogaster* larvae"

### Slide 1
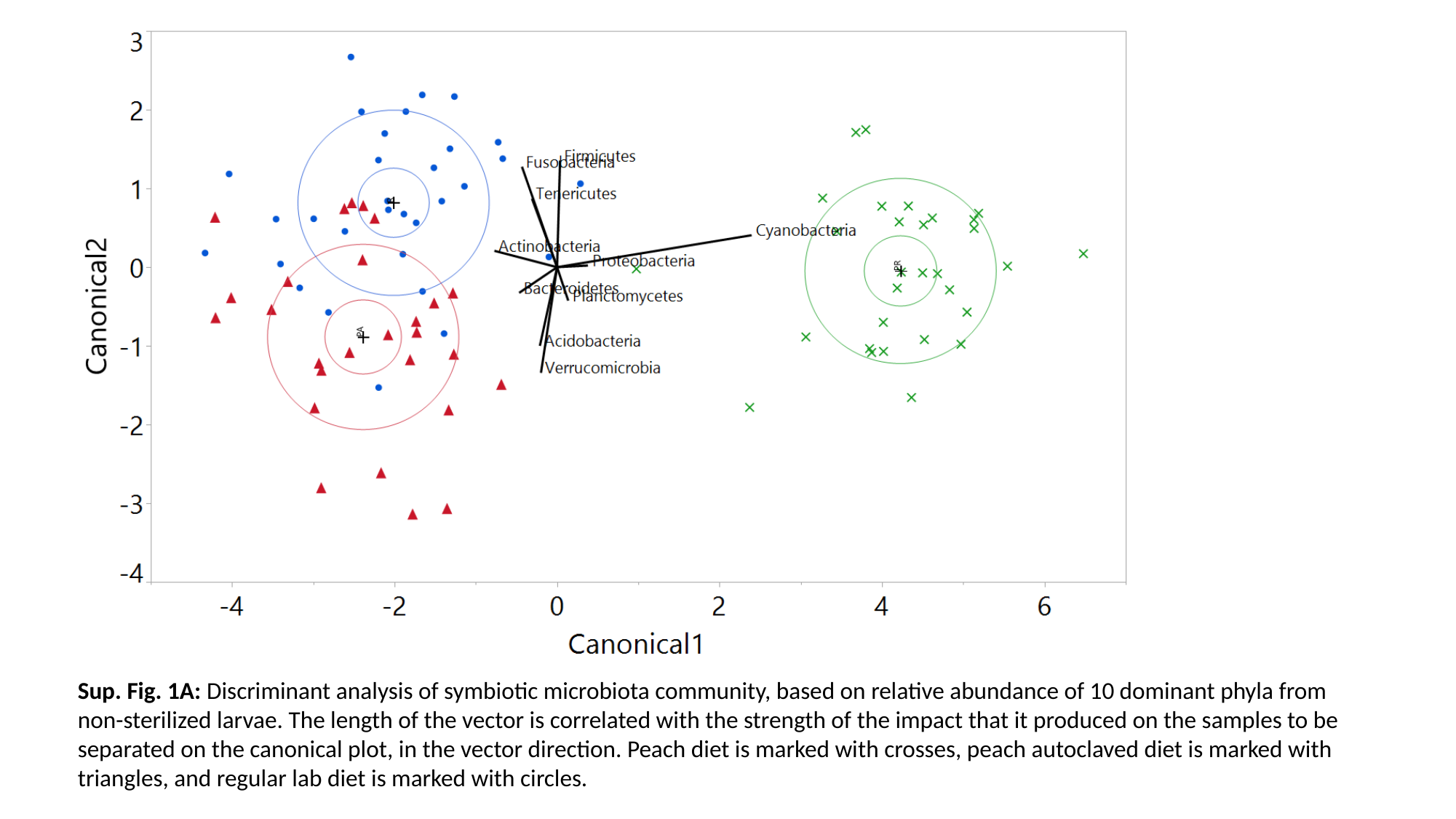

Sup. Fig. 1A: Discriminant analysis of symbiotic microbiota community, based on relative abundance of 10 dominant phyla from non-sterilized larvae. The length of the vector is correlated with the strength of the impact that it produced on the samples to be separated on the canonical plot, in the vector direction. Peach diet is marked with crosses, peach autoclaved diet is marked with triangles, and regular lab diet is marked with circles.

### Slide 2
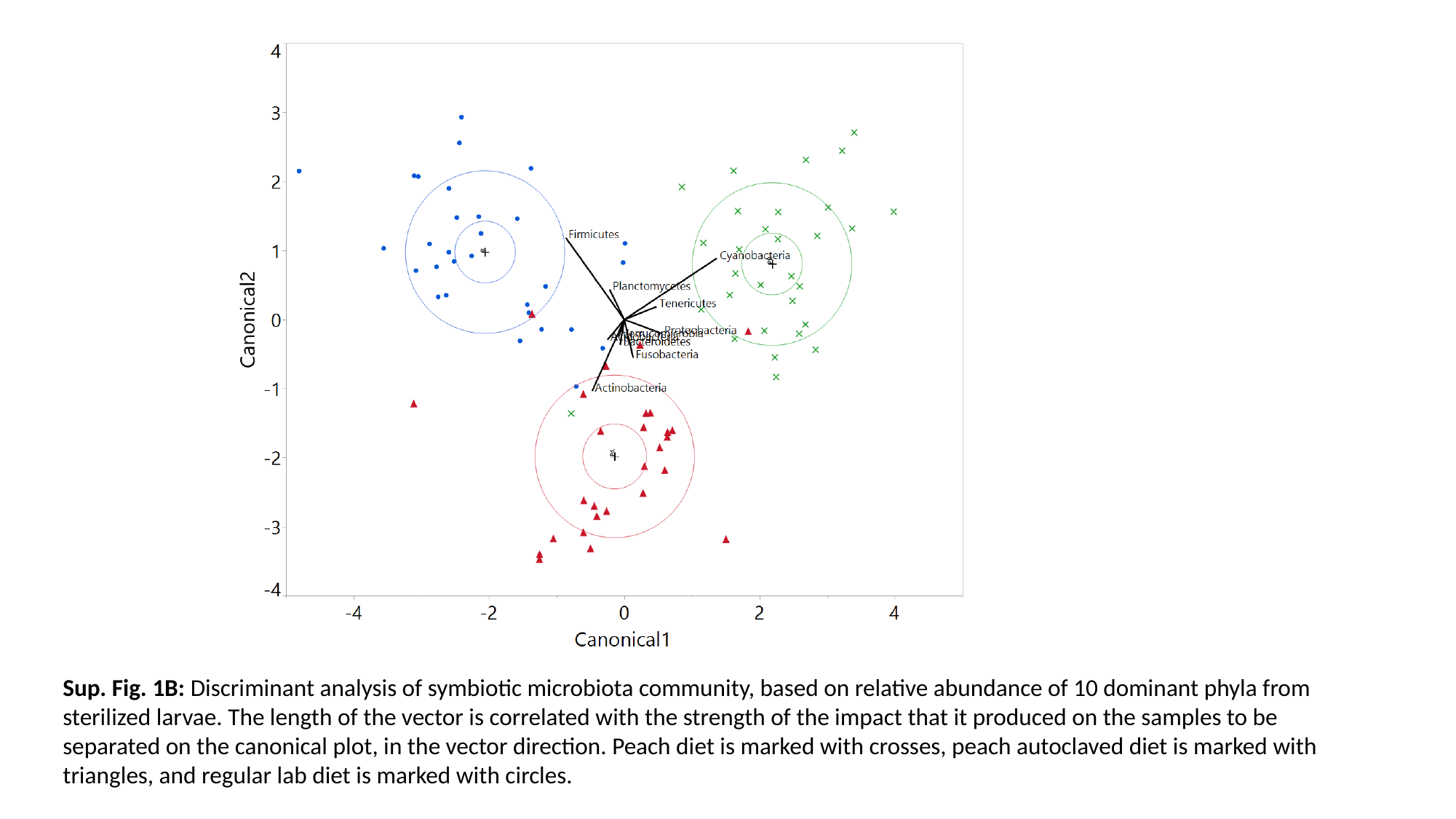

Sup. Fig. 1B: Discriminant analysis of symbiotic microbiota community, based on relative abundance of 10 dominant phyla from sterilized larvae. The length of the vector is correlated with the strength of the impact that it produced on the samples to be separated on the canonical plot, in the vector direction. Peach diet is marked with crosses, peach autoclaved diet is marked with triangles, and regular lab diet is marked with circles.

### Slide 3
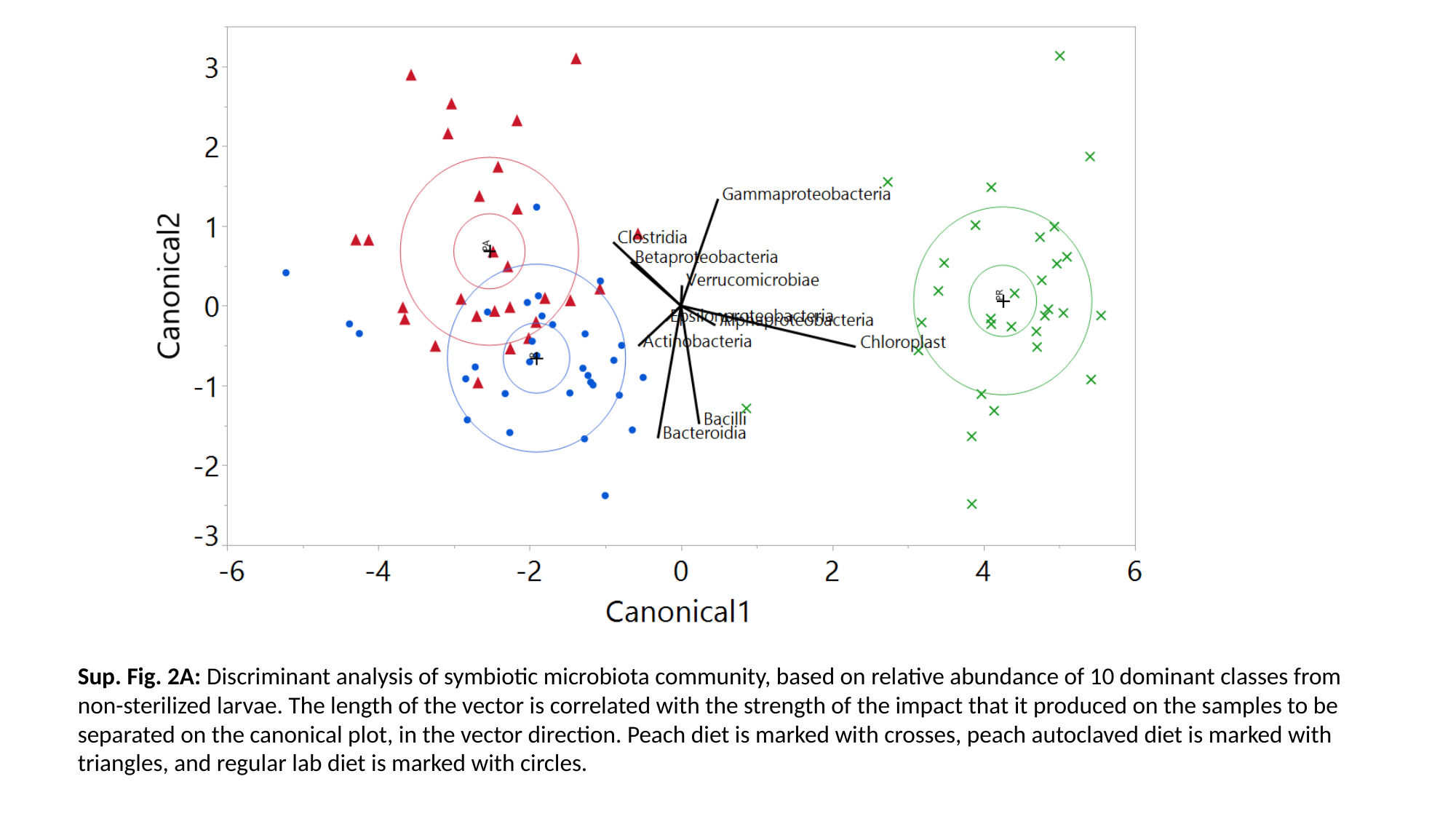

Sup. Fig. 2A: Discriminant analysis of symbiotic microbiota community, based on relative abundance of 10 dominant classes from non-sterilized larvae. The length of the vector is correlated with the strength of the impact that it produced on the samples to be separated on the canonical plot, in the vector direction. Peach diet is marked with crosses, peach autoclaved diet is marked with triangles, and regular lab diet is marked with circles.

### Slide 4
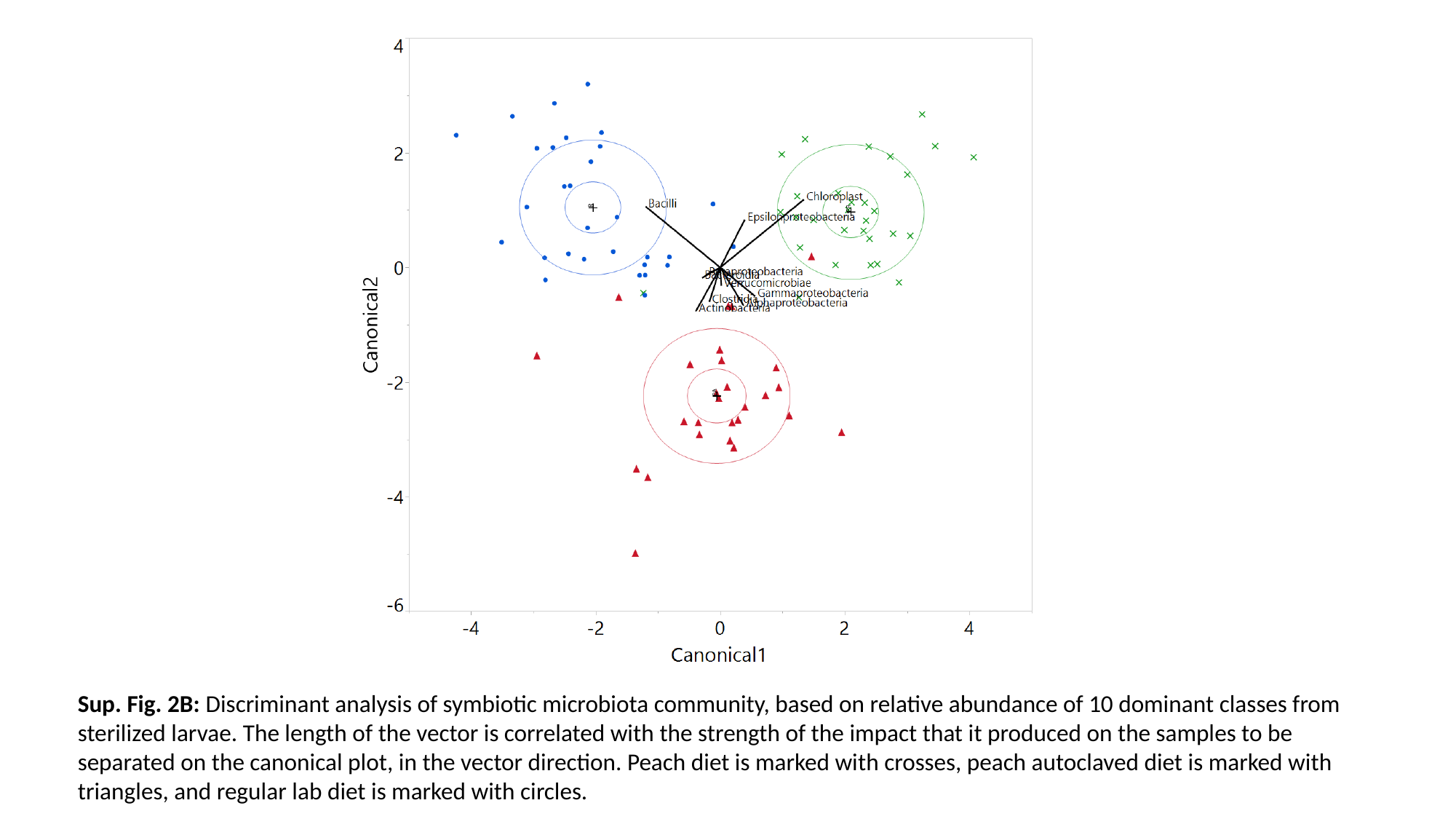

Sup. Fig. 2B: Discriminant analysis of symbiotic microbiota community, based on relative abundance of 10 dominant classes from sterilized larvae. The length of the vector is correlated with the strength of the impact that it produced on the samples to be separated on the canonical plot, in the vector direction. Peach diet is marked with crosses, peach autoclaved diet is marked with triangles, and regular lab diet is marked with circles.

### Slide 5
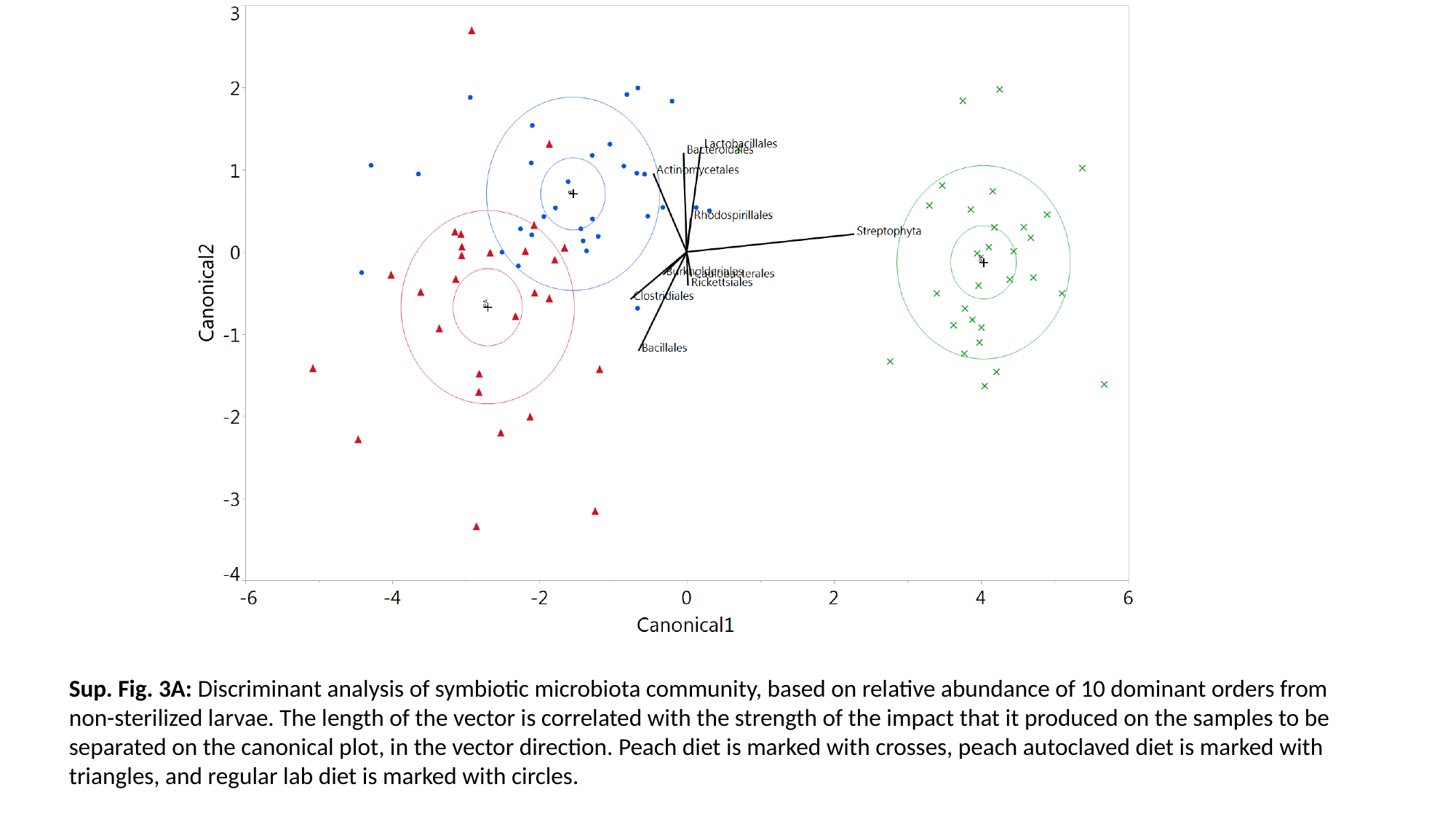

Sup. Fig. 3A: Discriminant analysis of symbiotic microbiota community, based on relative abundance of 10 dominant orders from non-sterilized larvae. The length of the vector is correlated with the strength of the impact that it produced on the samples to be separated on the canonical plot, in the vector direction. Peach diet is marked with crosses, peach autoclaved diet is marked with triangles, and regular lab diet is marked with circles.

### Slide 6
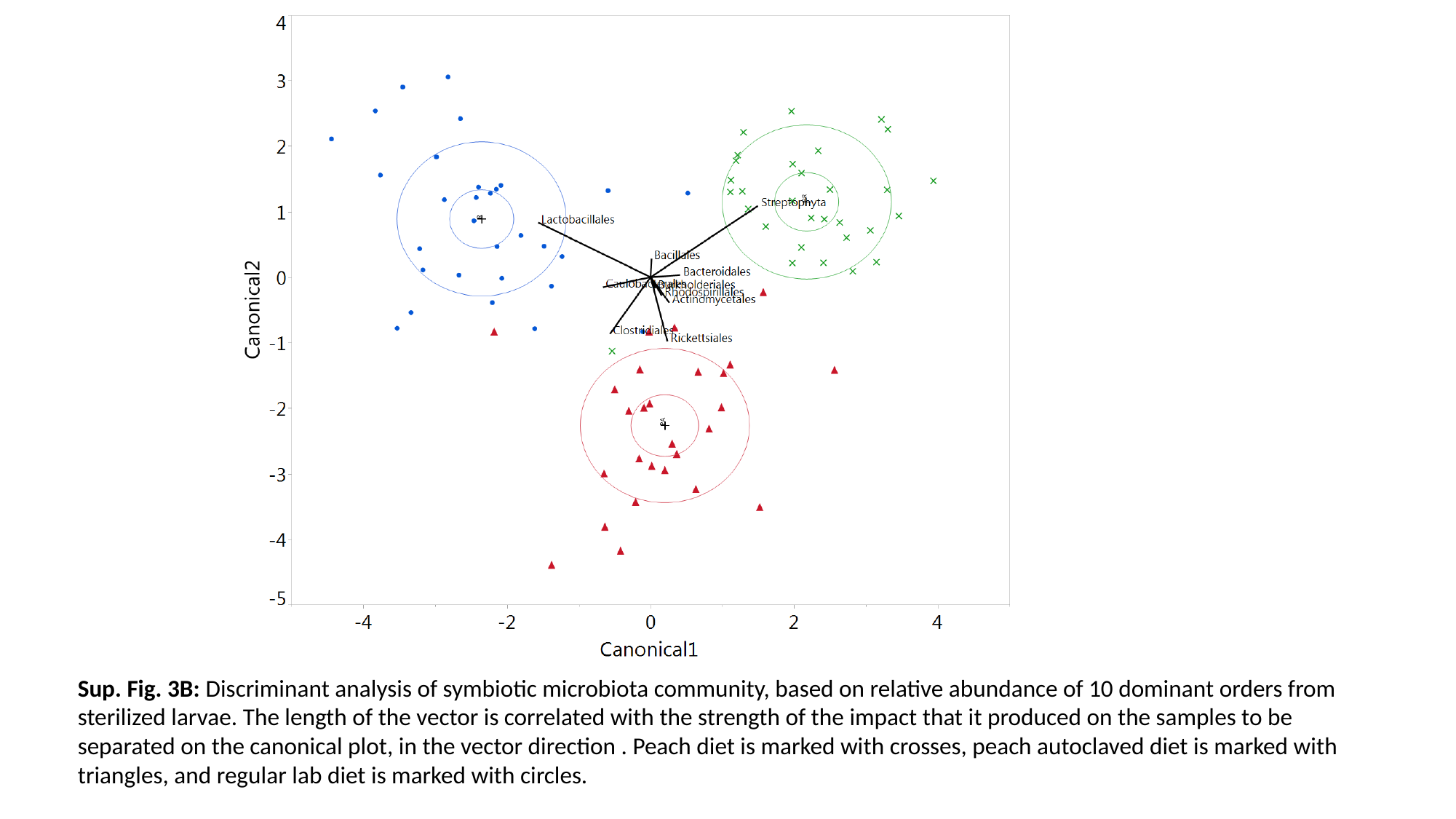

Sup. Fig. 3B: Discriminant analysis of symbiotic microbiota community, based on relative abundance of 10 dominant orders from sterilized larvae. The length of the vector is correlated with the strength of the impact that it produced on the samples to be separated on the canonical plot, in the vector direction . Peach diet is marked with crosses, peach autoclaved diet is marked with triangles, and regular lab diet is marked with circles.

### Slide 7
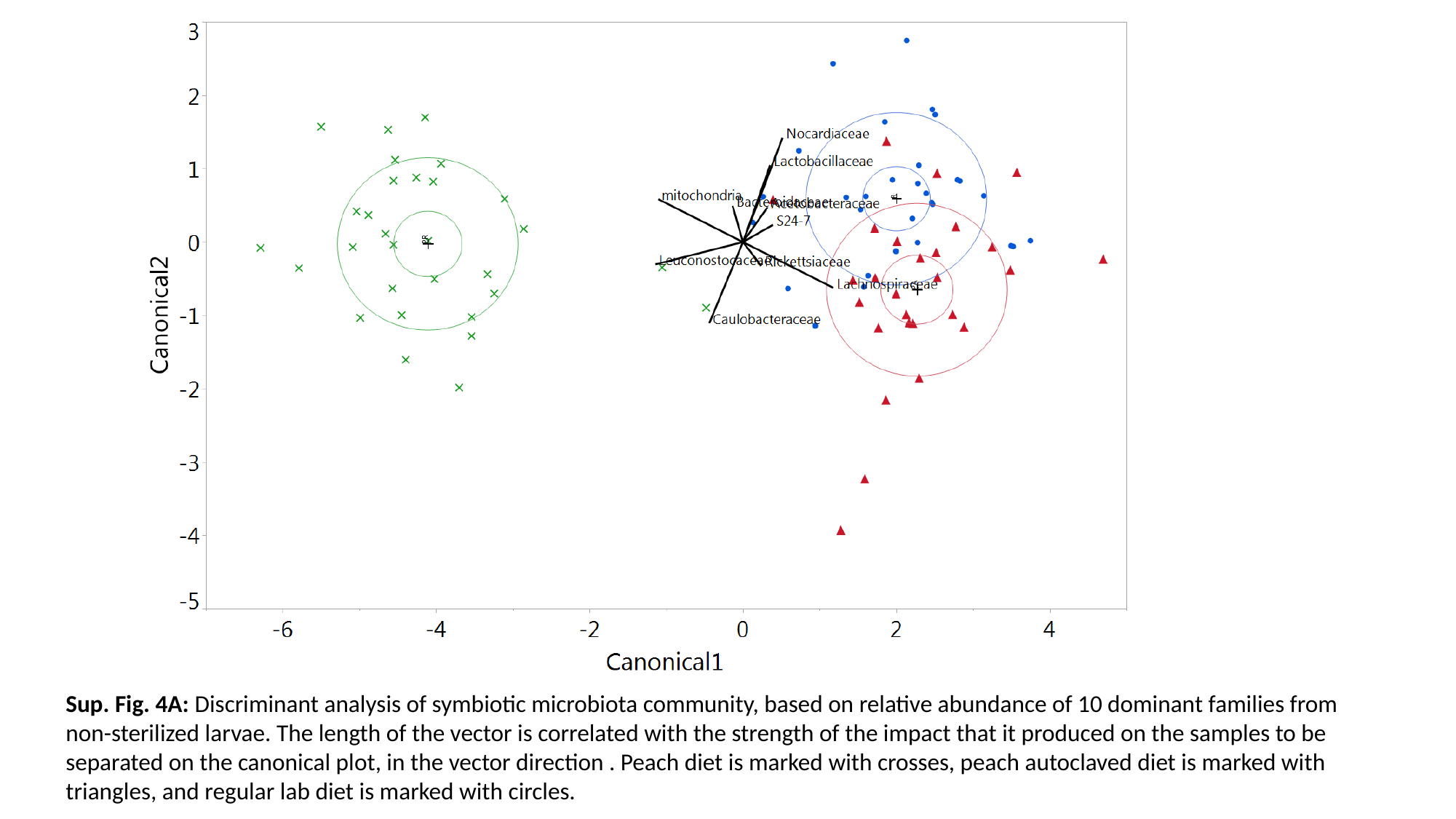

Sup. Fig. 4A: Discriminant analysis of symbiotic microbiota community, based on relative abundance of 10 dominant families from non-sterilized larvae. The length of the vector is correlated with the strength of the impact that it produced on the samples to be separated on the canonical plot, in the vector direction . Peach diet is marked with crosses, peach autoclaved diet is marked with triangles, and regular lab diet is marked with circles.

### Slide 8
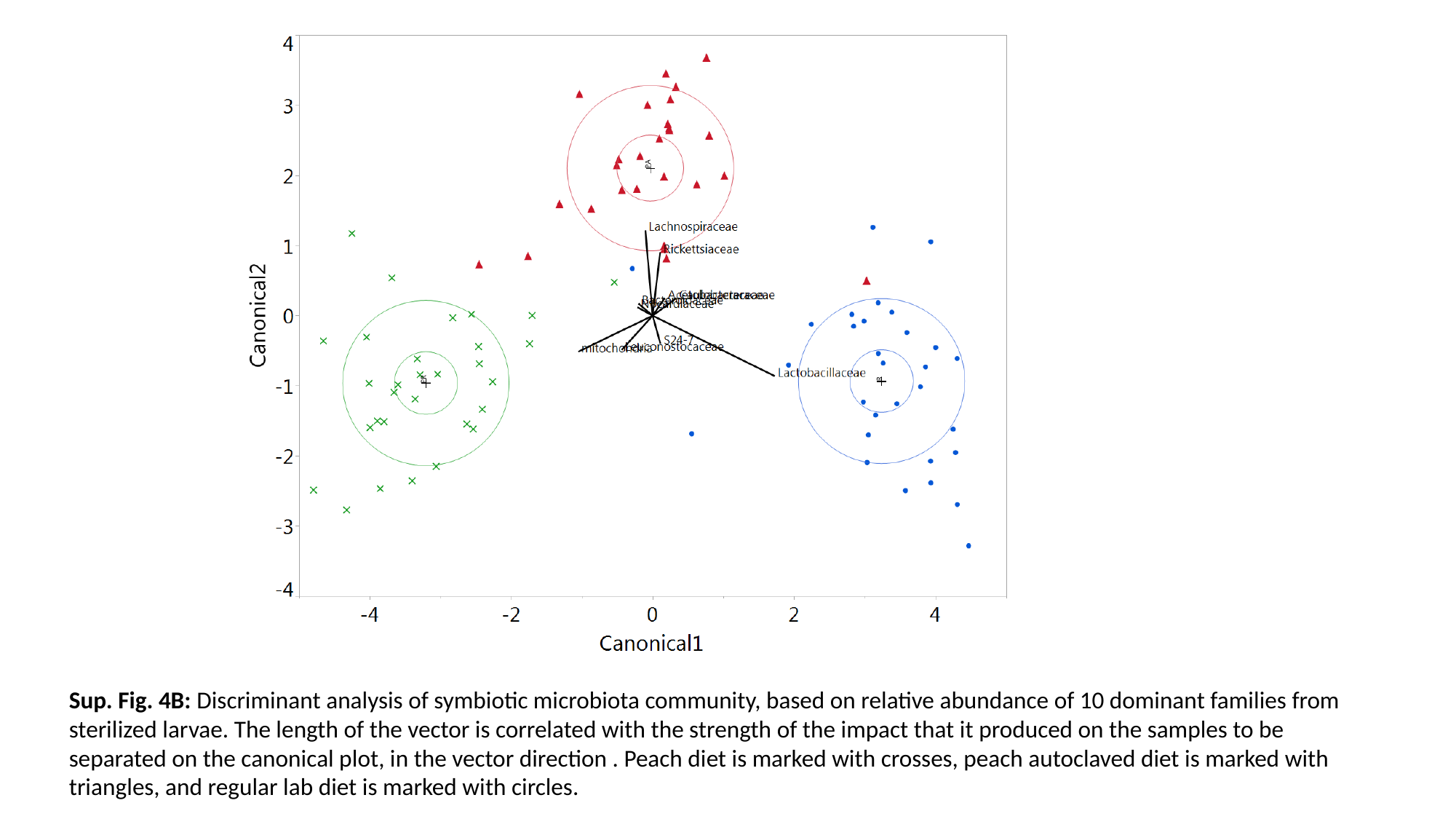

Sup. Fig. 4B: Discriminant analysis of symbiotic microbiota community, based on relative abundance of 10 dominant families from sterilized larvae. The length of the vector is correlated with the strength of the impact that it produced on the samples to be separated on the canonical plot, in the vector direction . Peach diet is marked with crosses, peach autoclaved diet is marked with triangles, and regular lab diet is marked with circles.

### Slide 9
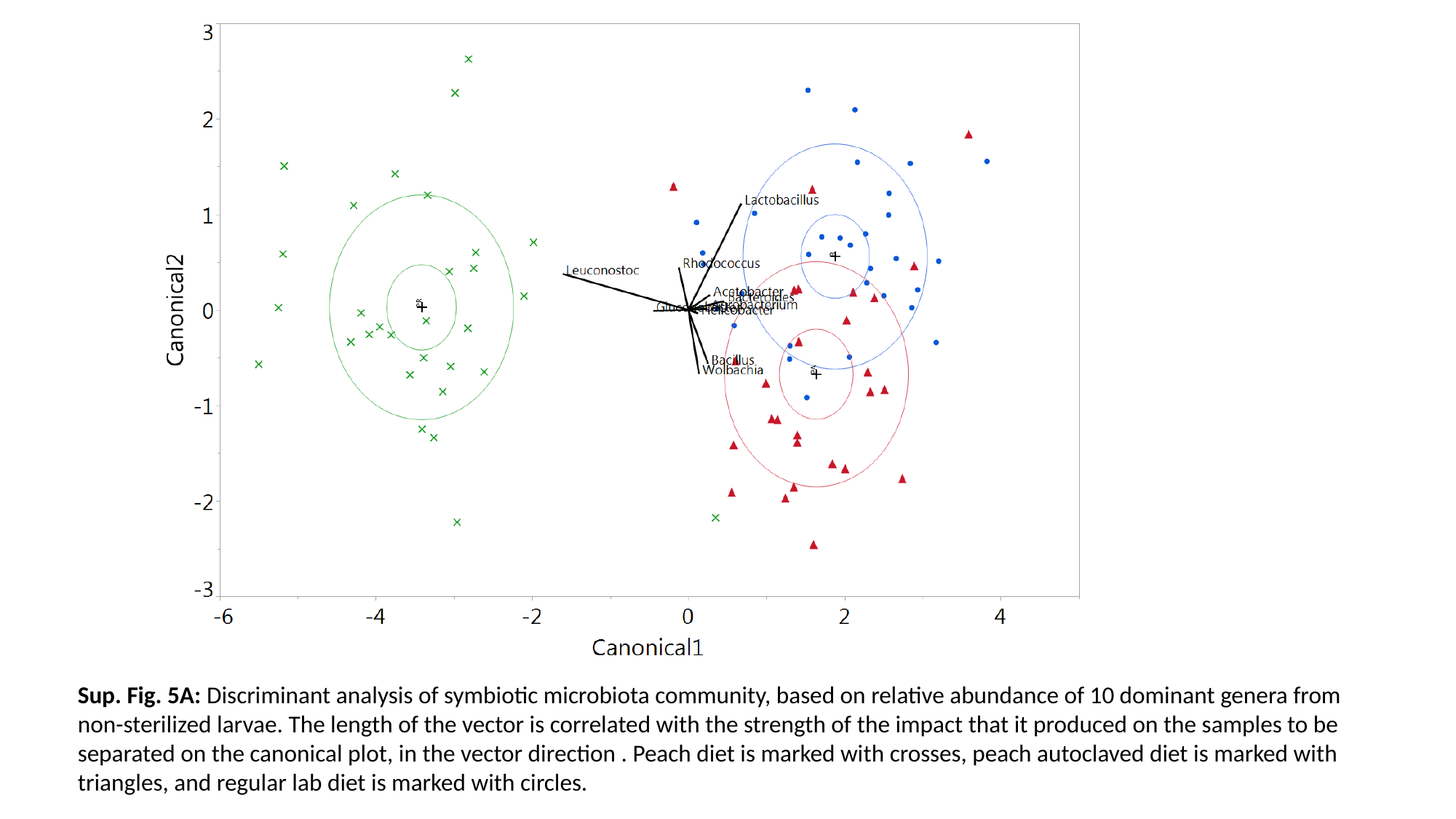

Sup. Fig. 5A: Discriminant analysis of symbiotic microbiota community, based on relative abundance of 10 dominant genera from non-sterilized larvae. The length of the vector is correlated with the strength of the impact that it produced on the samples to be separated on the canonical plot, in the vector direction . Peach diet is marked with crosses, peach autoclaved diet is marked with triangles, and regular lab diet is marked with circles.

### Slide 10
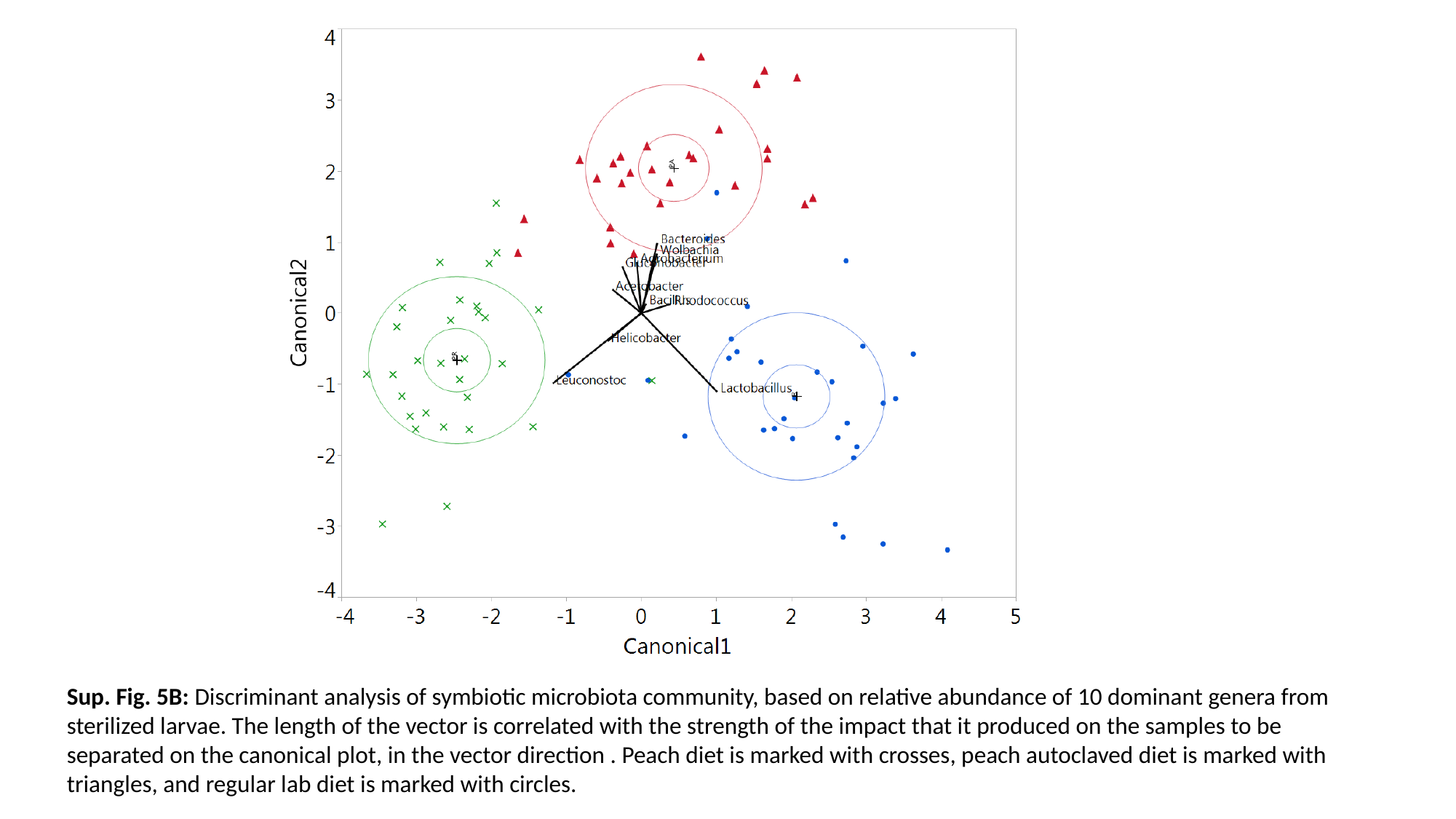

Sup. Fig. 5B: Discriminant analysis of symbiotic microbiota community, based on relative abundance of 10 dominant genera from sterilized larvae. The length of the vector is correlated with the strength of the impact that it produced on the samples to be separated on the canonical plot, in the vector direction . Peach diet is marked with crosses, peach autoclaved diet is marked with triangles, and regular lab diet is marked with circles.

### Slide 11
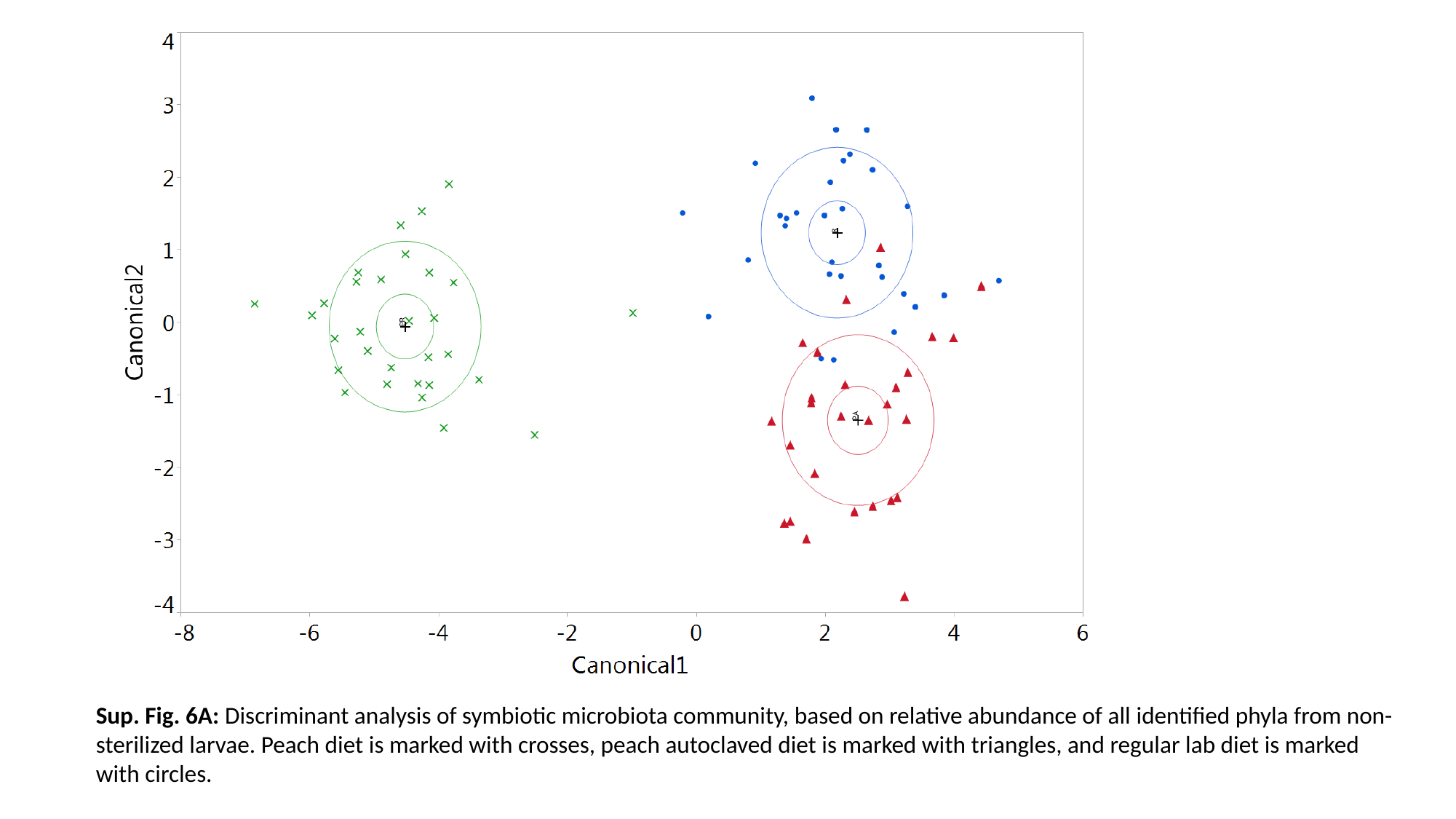

Sup. Fig. 6A: Discriminant analysis of symbiotic microbiota community, based on relative abundance of all identified phyla from non-sterilized larvae. Peach diet is marked with crosses, peach autoclaved diet is marked with triangles, and regular lab diet is marked with circles.

### Slide 12
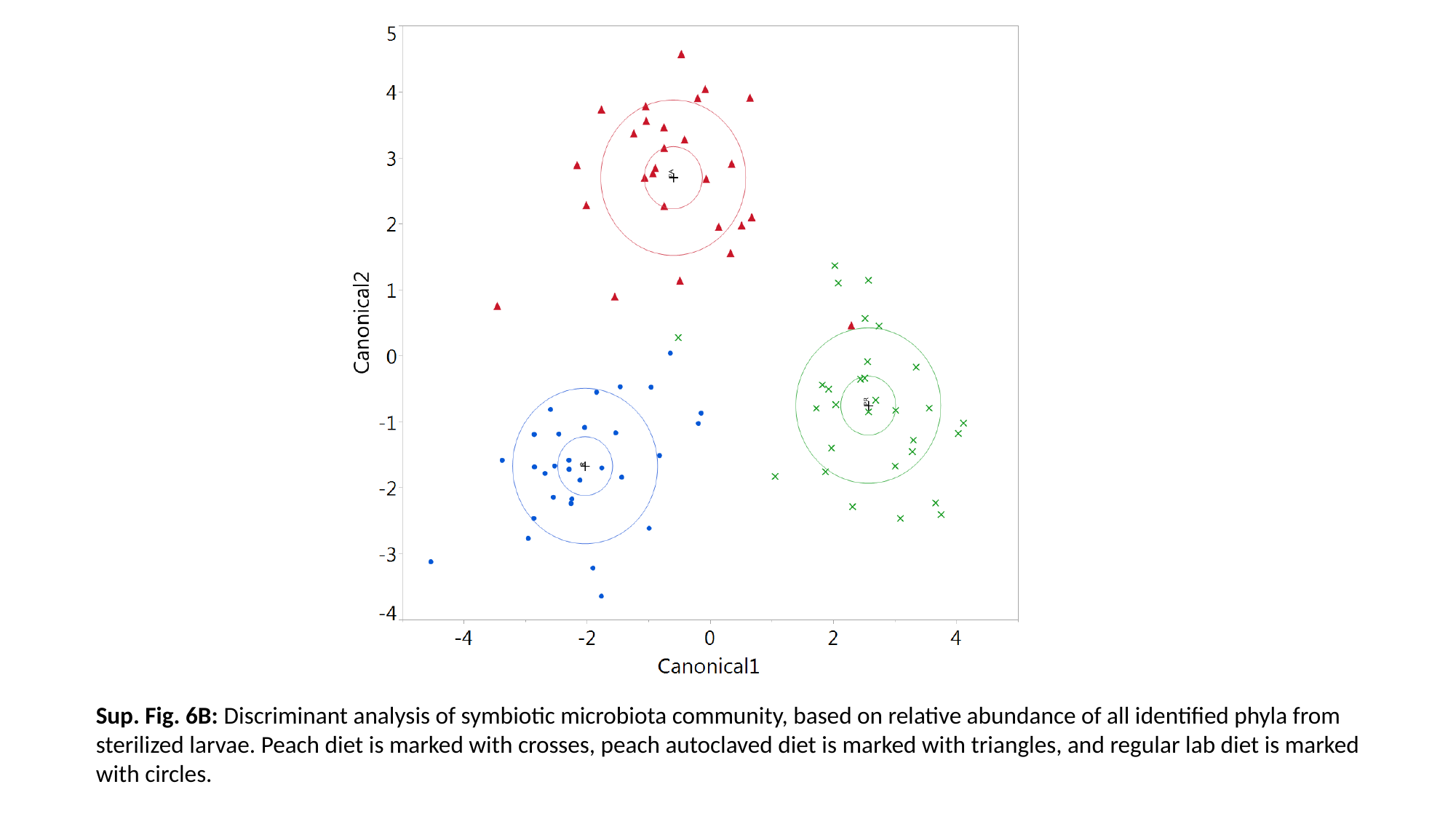

Sup. Fig. 6B: Discriminant analysis of symbiotic microbiota community, based on relative abundance of all identified phyla from sterilized larvae. Peach diet is marked with crosses, peach autoclaved diet is marked with triangles, and regular lab diet is marked with circles.

### Slide 13
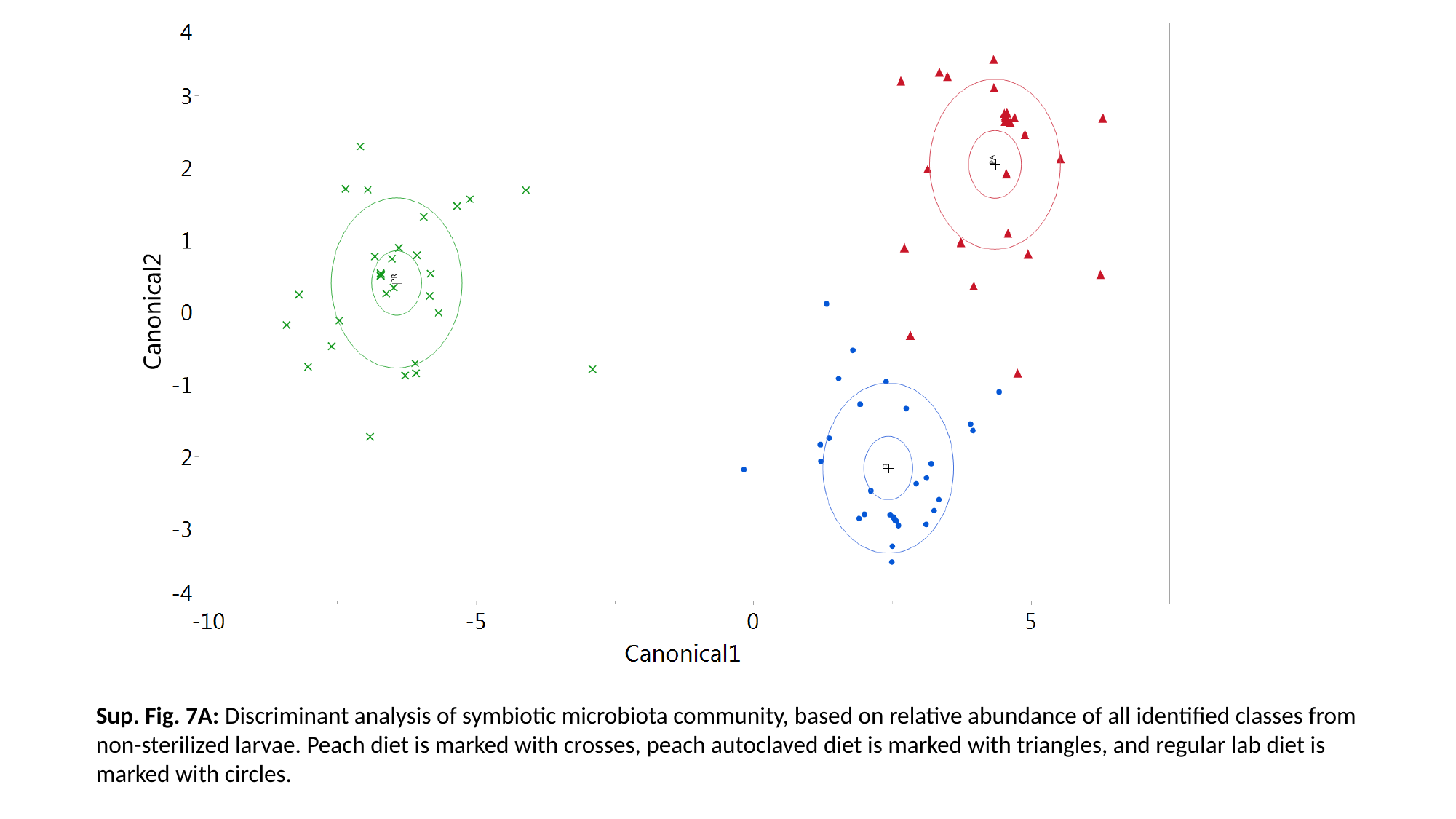

Sup. Fig. 7A: Discriminant analysis of symbiotic microbiota community, based on relative abundance of all identified classes from non-sterilized larvae. Peach diet is marked with crosses, peach autoclaved diet is marked with triangles, and regular lab diet is marked with circles.

### Slide 14
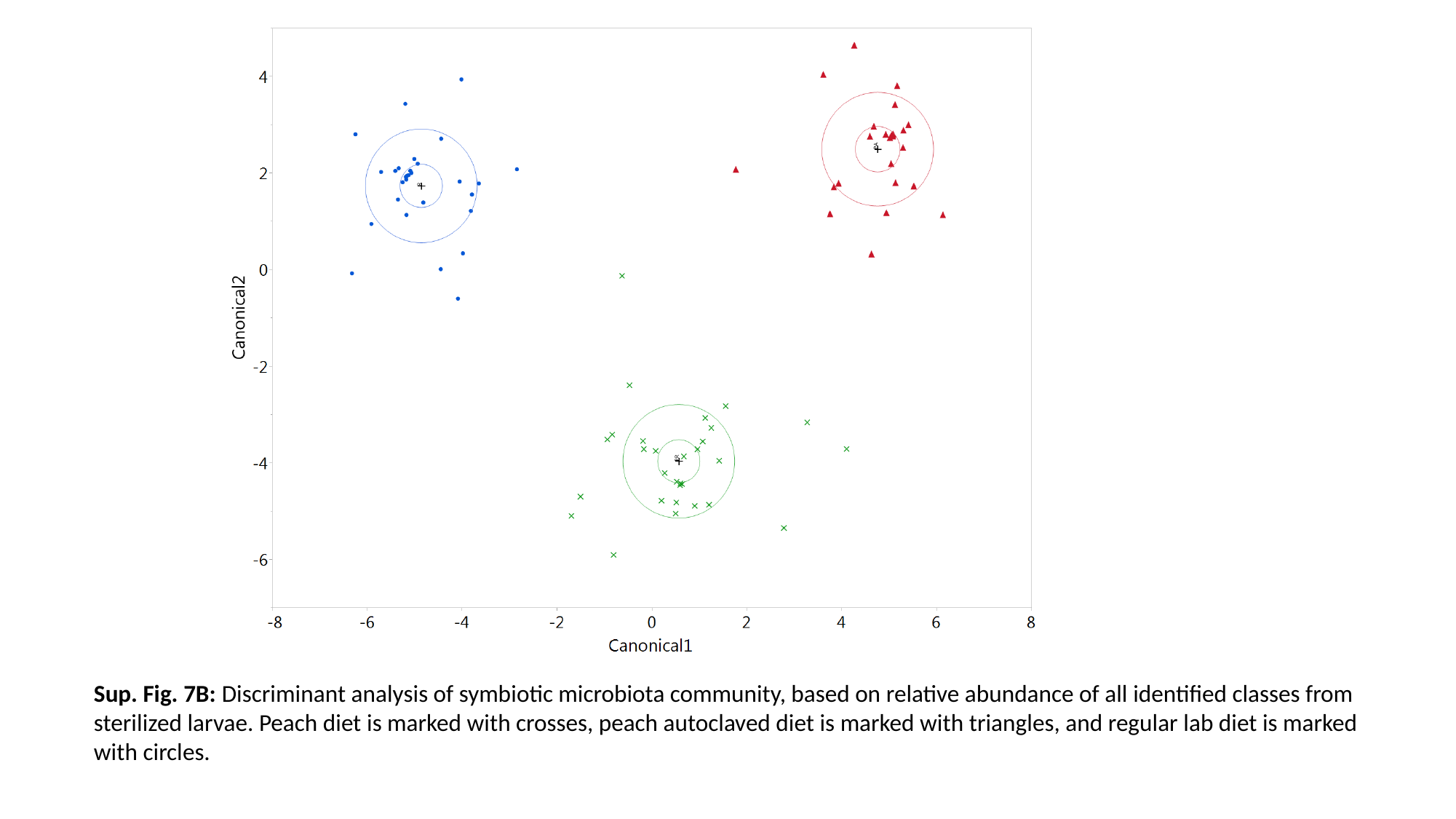

Sup. Fig. 7B: Discriminant analysis of symbiotic microbiota community, based on relative abundance of all identified classes from sterilized larvae. Peach diet is marked with crosses, peach autoclaved diet is marked with triangles, and regular lab diet is marked with circles.

### Slide 15
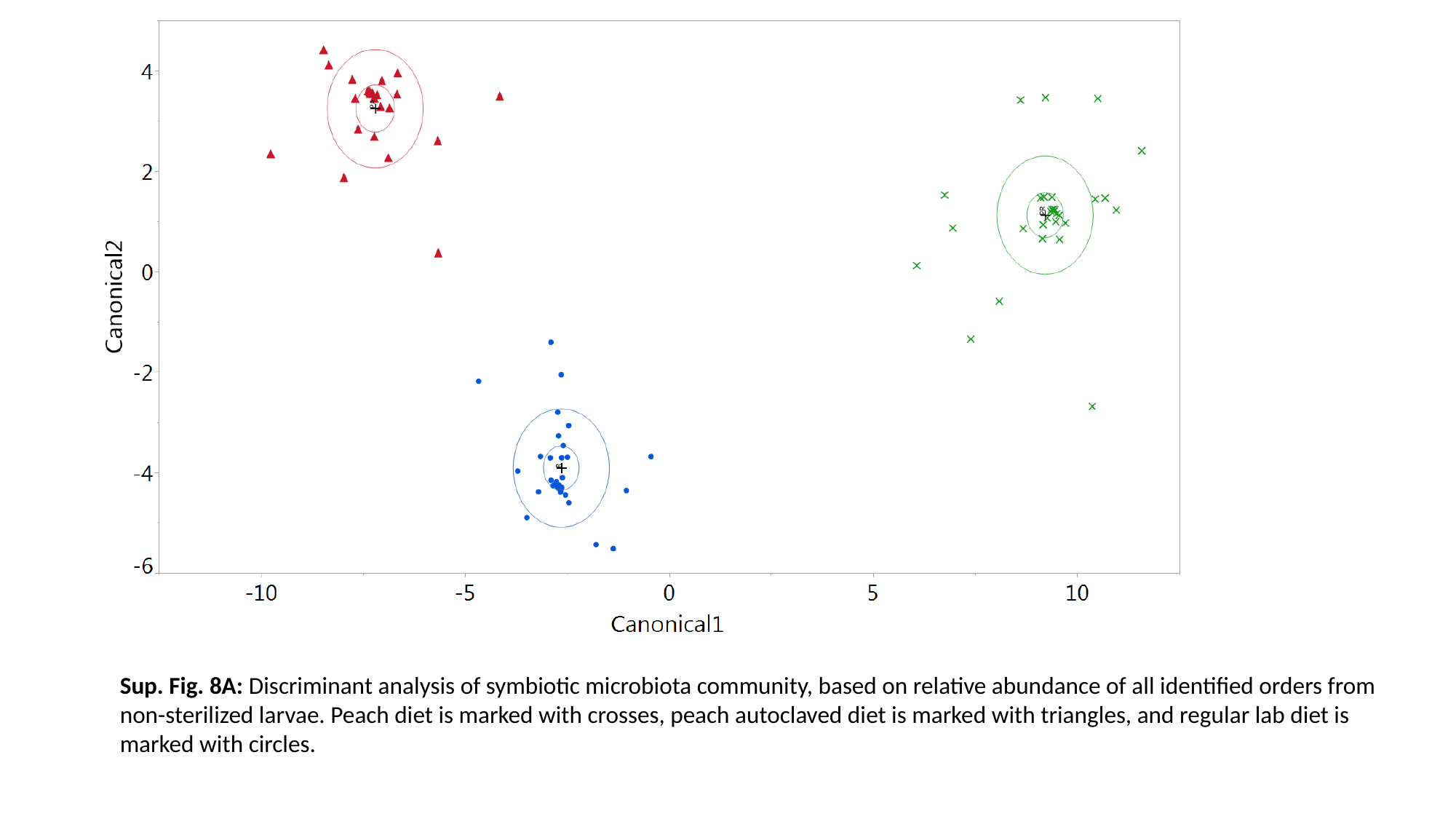

Sup. Fig. 8A: Discriminant analysis of symbiotic microbiota community, based on relative abundance of all identified orders from non-sterilized larvae. Peach diet is marked with crosses, peach autoclaved diet is marked with triangles, and regular lab diet is marked with circles.

### Slide 16
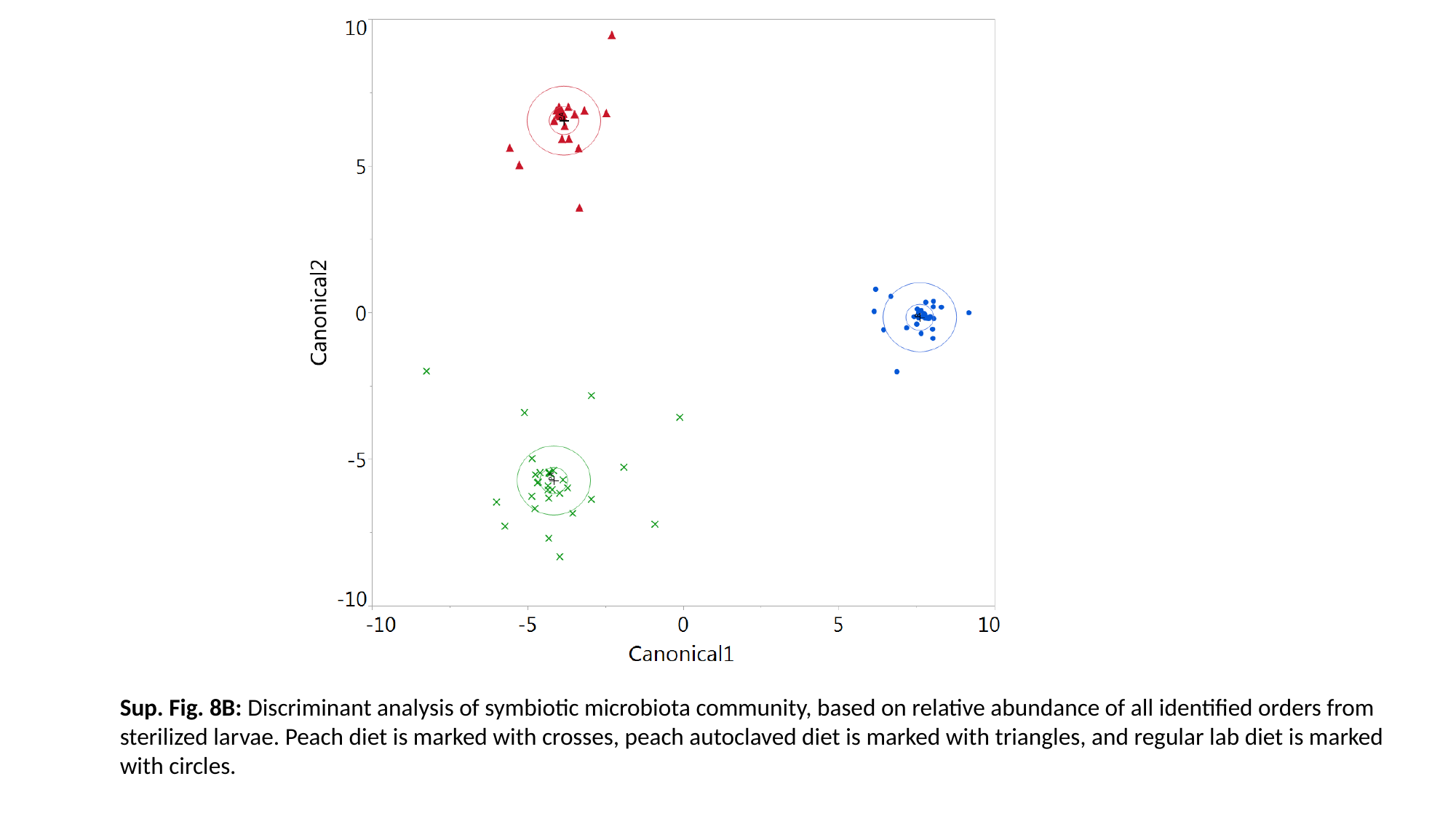

Sup. Fig. 8B: Discriminant analysis of symbiotic microbiota community, based on relative abundance of all identified orders from sterilized larvae. Peach diet is marked with crosses, peach autoclaved diet is marked with triangles, and regular lab diet is marked with circles.

### Slide 17
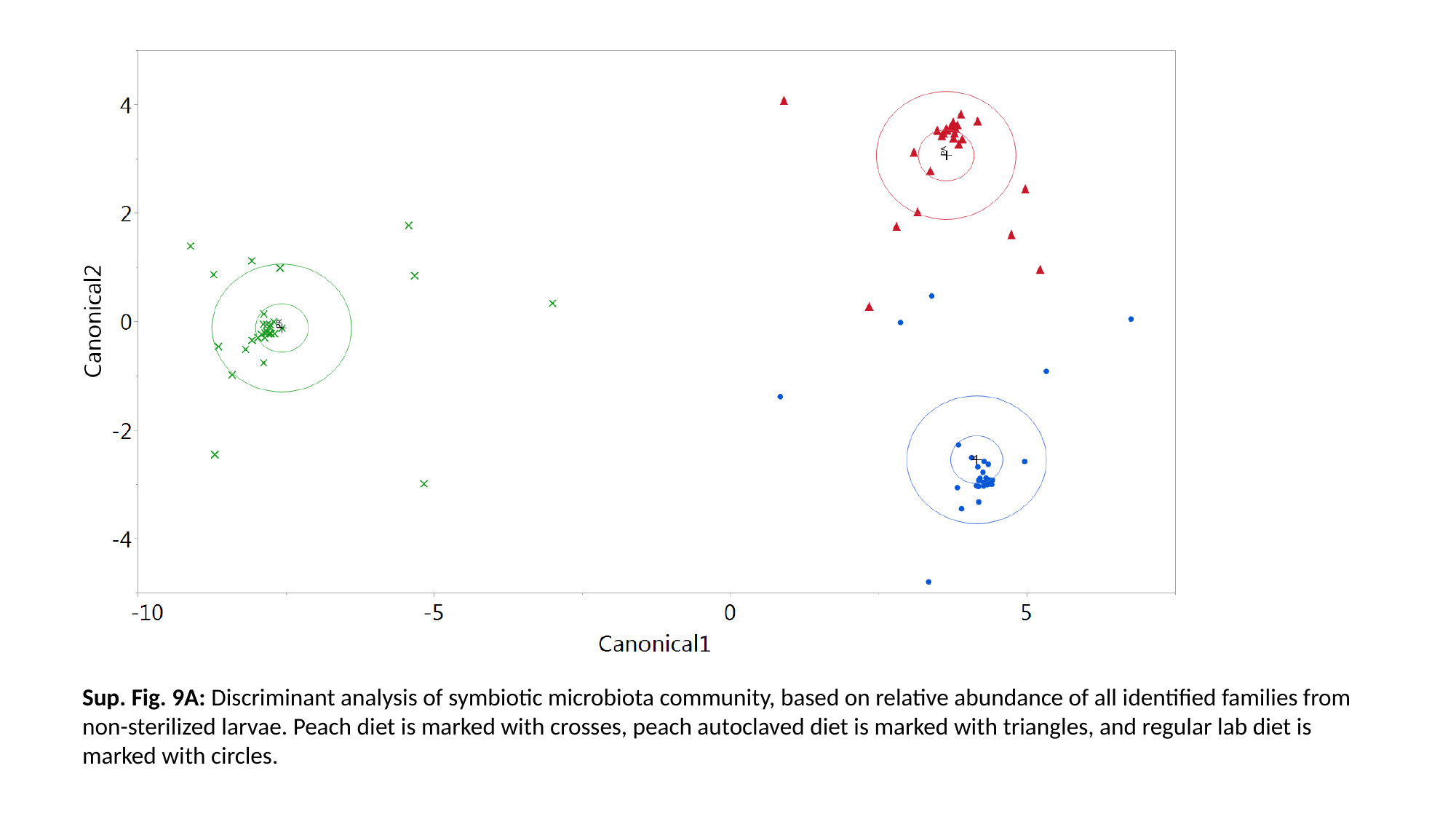

Sup. Fig. 9A: Discriminant analysis of symbiotic microbiota community, based on relative abundance of all identified families from non-sterilized larvae. Peach diet is marked with crosses, peach autoclaved diet is marked with triangles, and regular lab diet is marked with circles.

### Slide 18
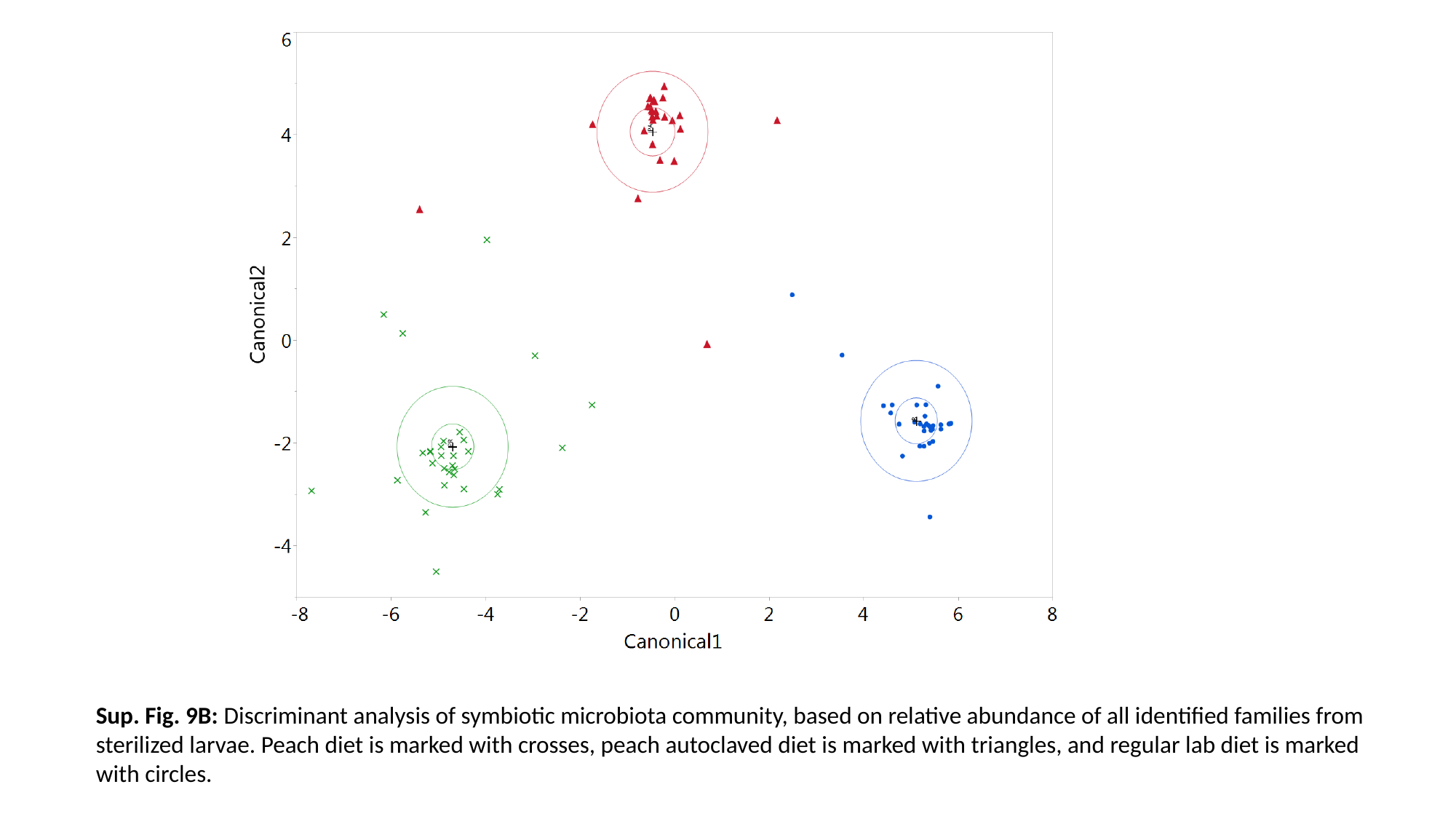

Sup. Fig. 9B: Discriminant analysis of symbiotic microbiota community, based on relative abundance of all identified families from sterilized larvae. Peach diet is marked with crosses, peach autoclaved diet is marked with triangles, and regular lab diet is marked with circles.

### Slide 19
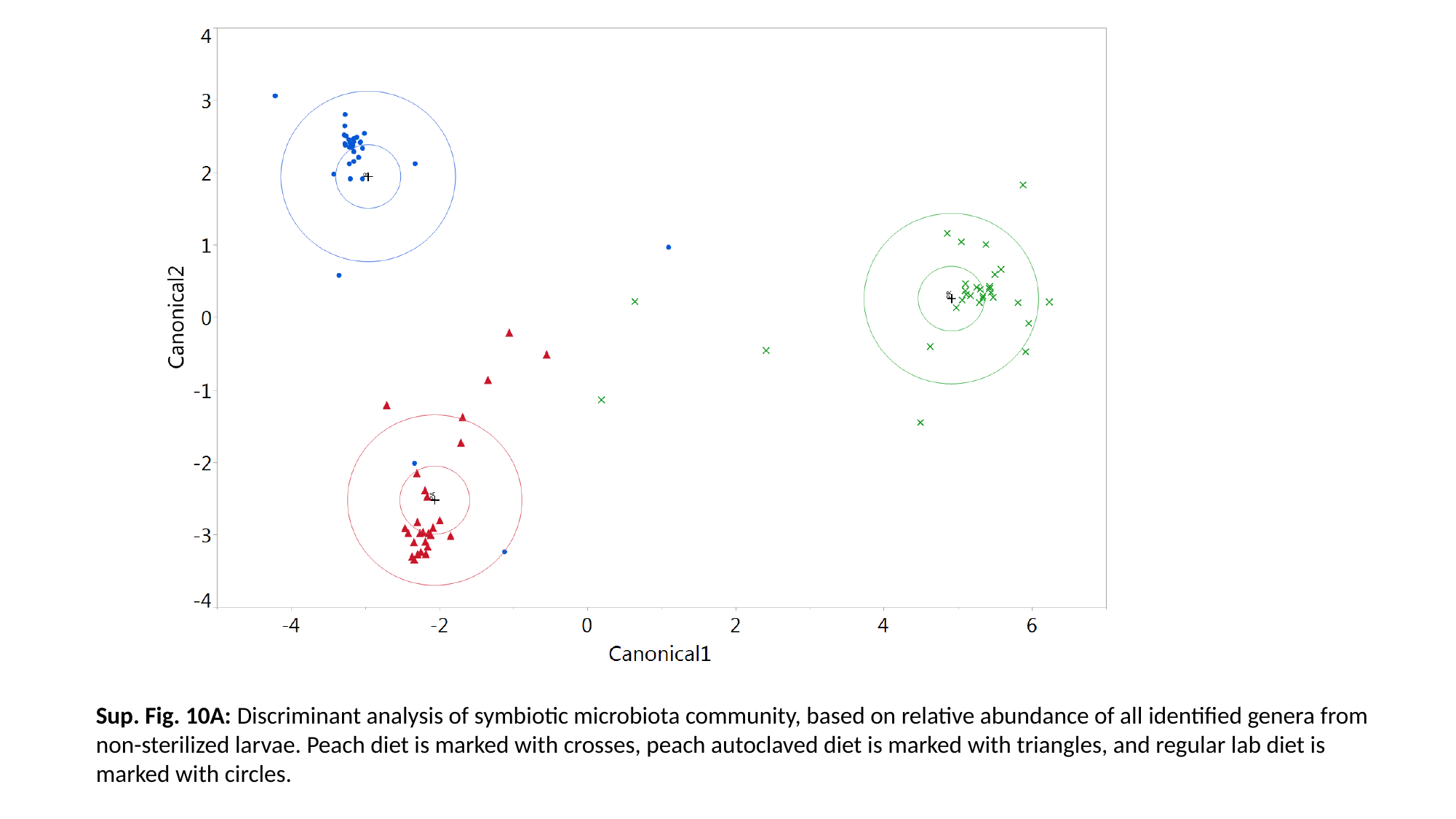

Sup. Fig. 10A: Discriminant analysis of symbiotic microbiota community, based on relative abundance of all identified genera from non-sterilized larvae. Peach diet is marked with crosses, peach autoclaved diet is marked with triangles, and regular lab diet is marked with circles.

### Slide 20
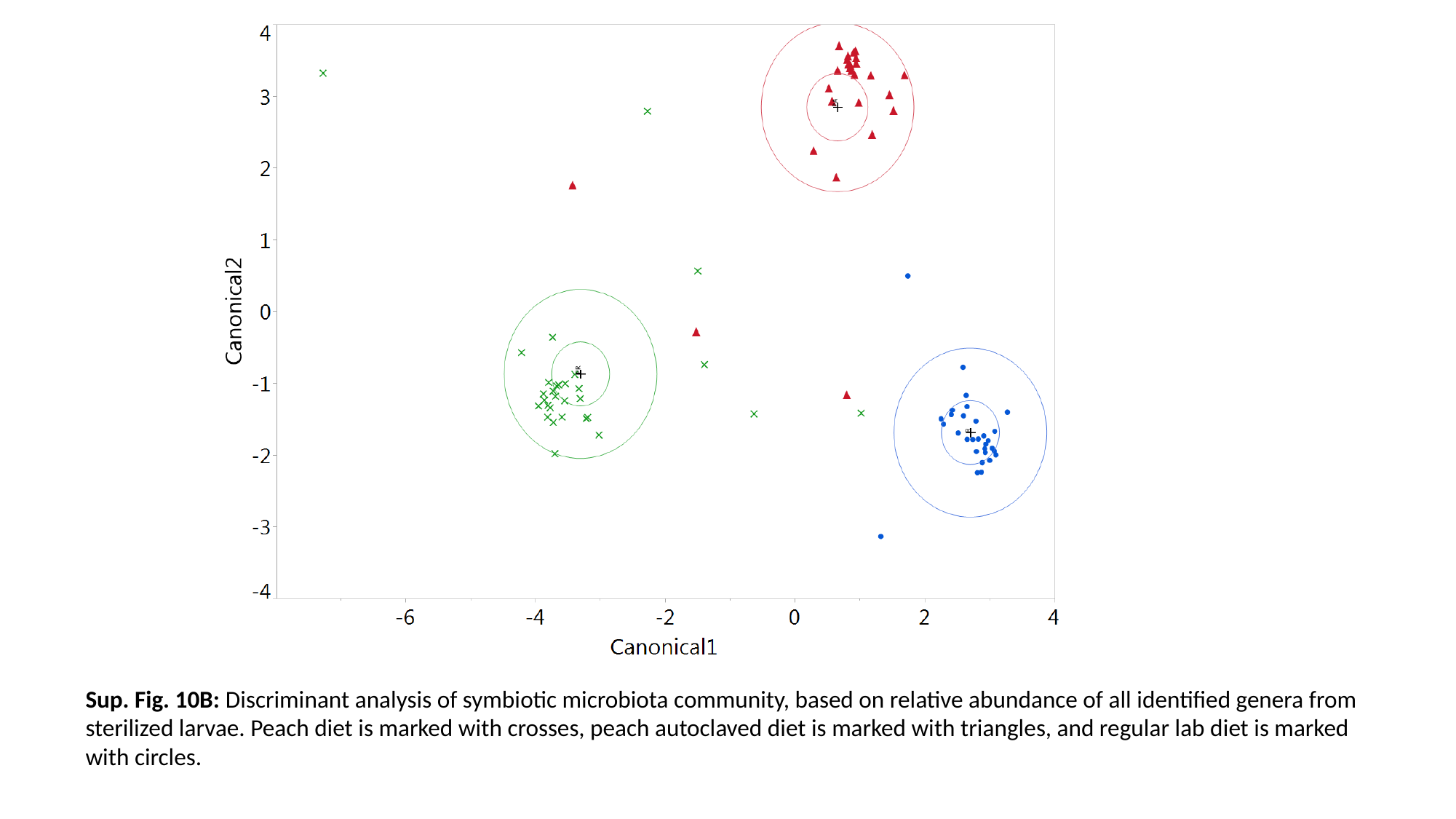

Sup. Fig. 10B: Discriminant analysis of symbiotic microbiota community, based on relative abundance of all identified genera from sterilized larvae. Peach diet is marked with crosses, peach autoclaved diet is marked with triangles, and regular lab diet is marked with circles.

### Slide 21
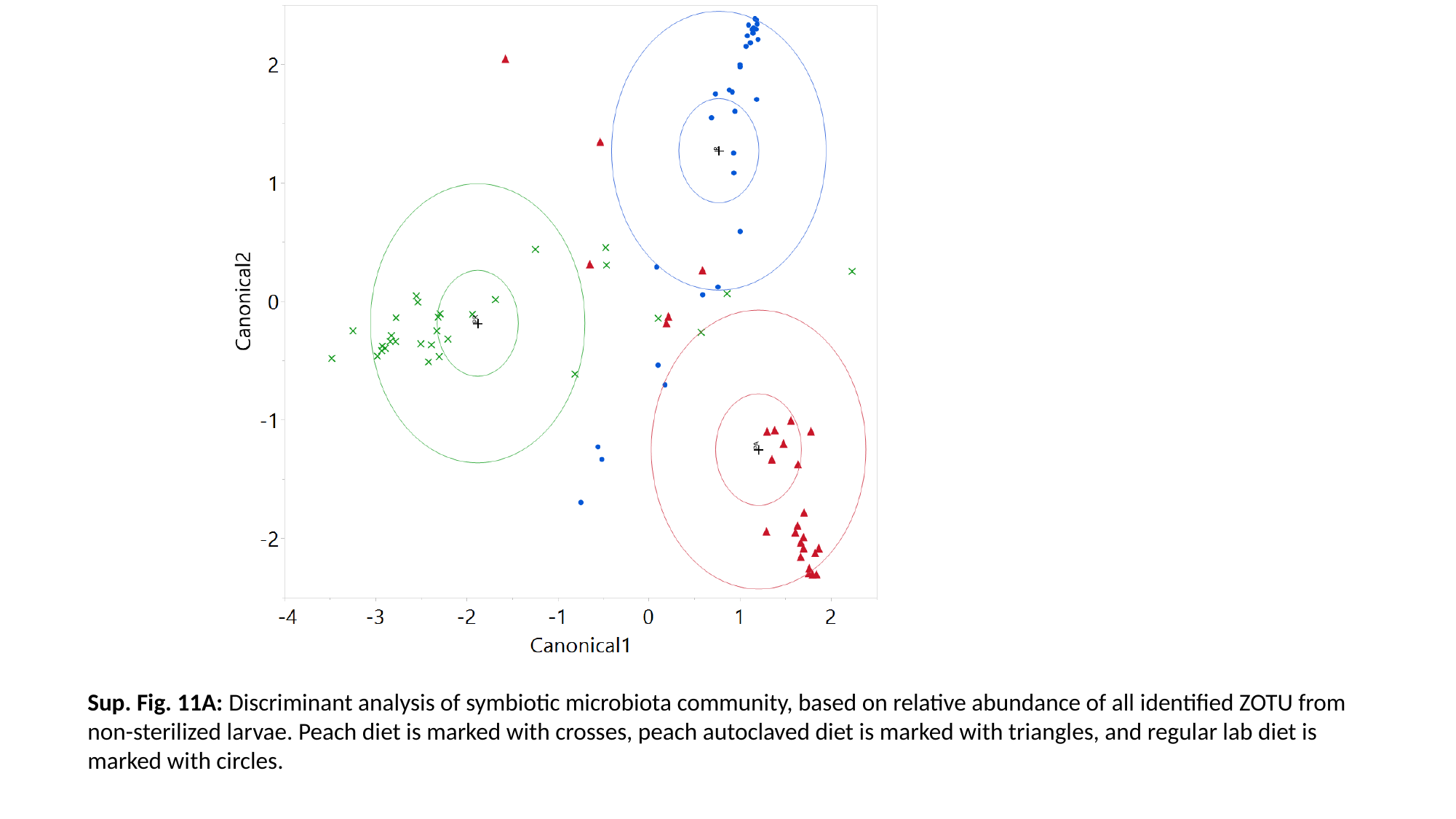

Sup. Fig. 11A: Discriminant analysis of symbiotic microbiota community, based on relative abundance of all identified ZOTU from non-sterilized larvae. Peach diet is marked with crosses, peach autoclaved diet is marked with triangles, and regular lab diet is marked with circles.

### Slide 22
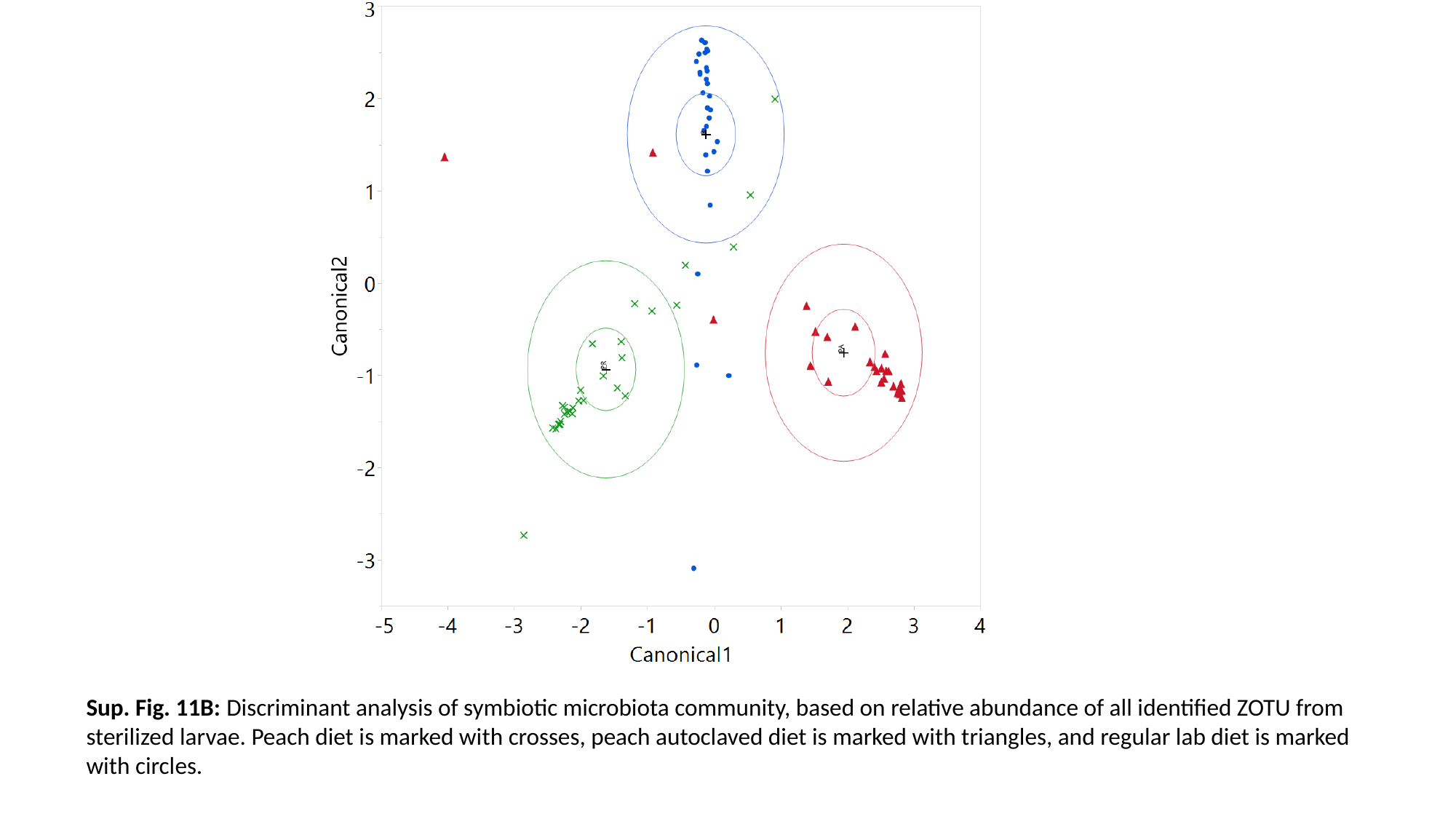

Sup. Fig. 11B: Discriminant analysis of symbiotic microbiota community, based on relative abundance of all identified ZOTU from sterilized larvae. Peach diet is marked with crosses, peach autoclaved diet is marked with triangles, and regular lab diet is marked with circles.

### Slide 23
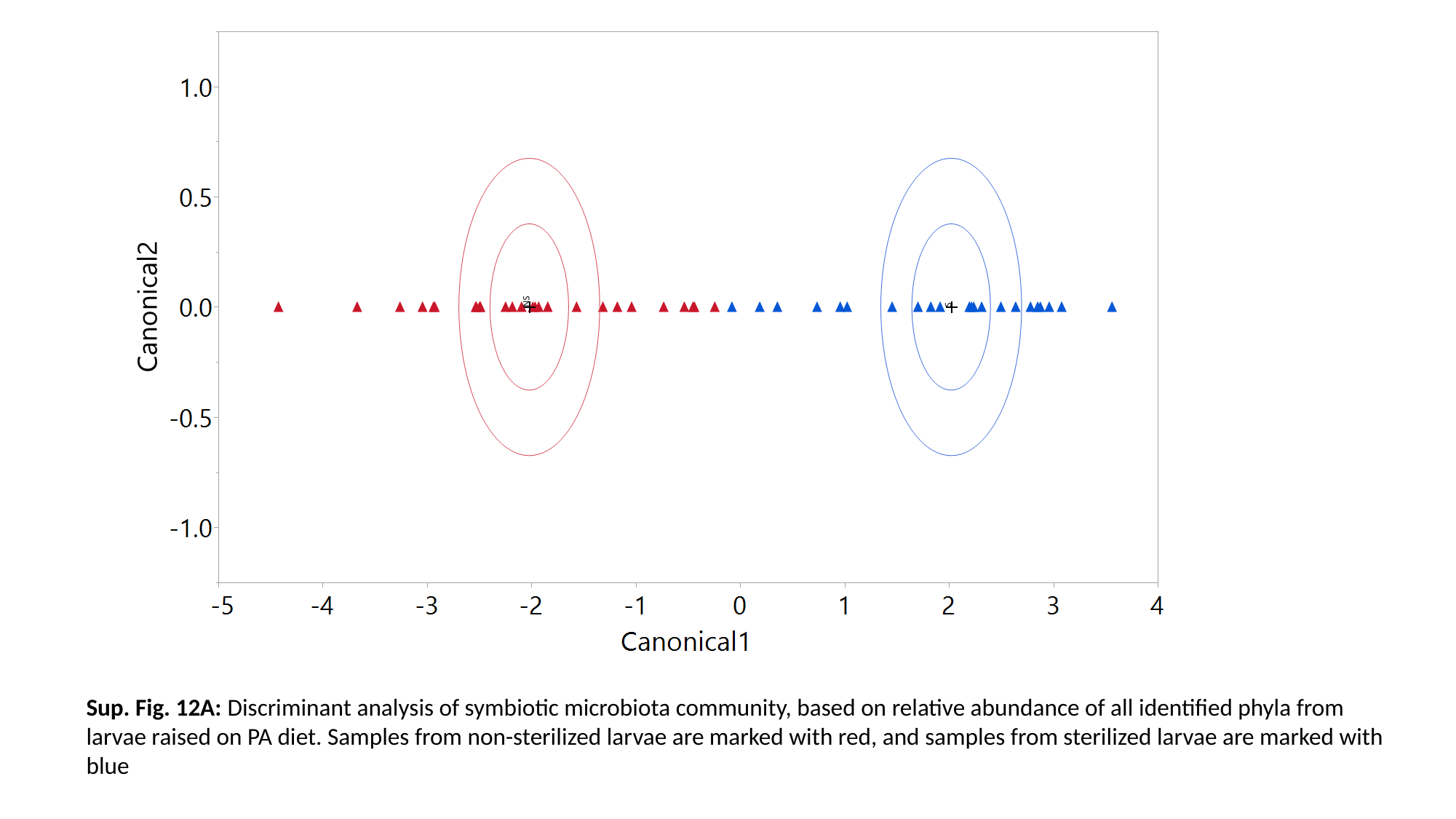

Sup. Fig. 12A: Discriminant analysis of symbiotic microbiota community, based on relative abundance of all identified phyla from larvae raised on PA diet. Samples from non-sterilized larvae are marked with red, and samples from sterilized larvae are marked with blue

### Slide 24
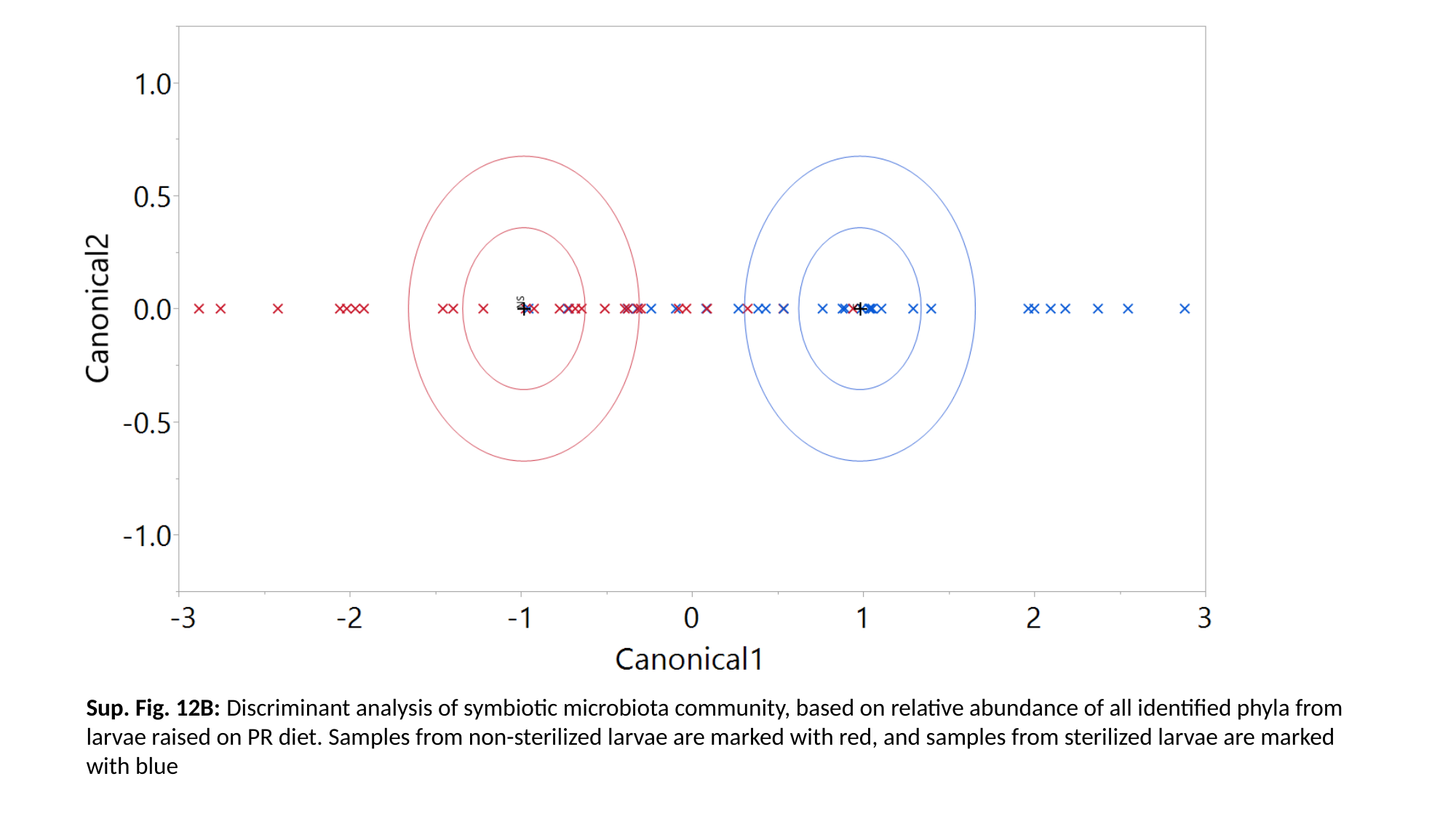

Sup. Fig. 12B: Discriminant analysis of symbiotic microbiota community, based on relative abundance of all identified phyla from larvae raised on PR diet. Samples from non-sterilized larvae are marked with red, and samples from sterilized larvae are marked with blue

### Slide 25
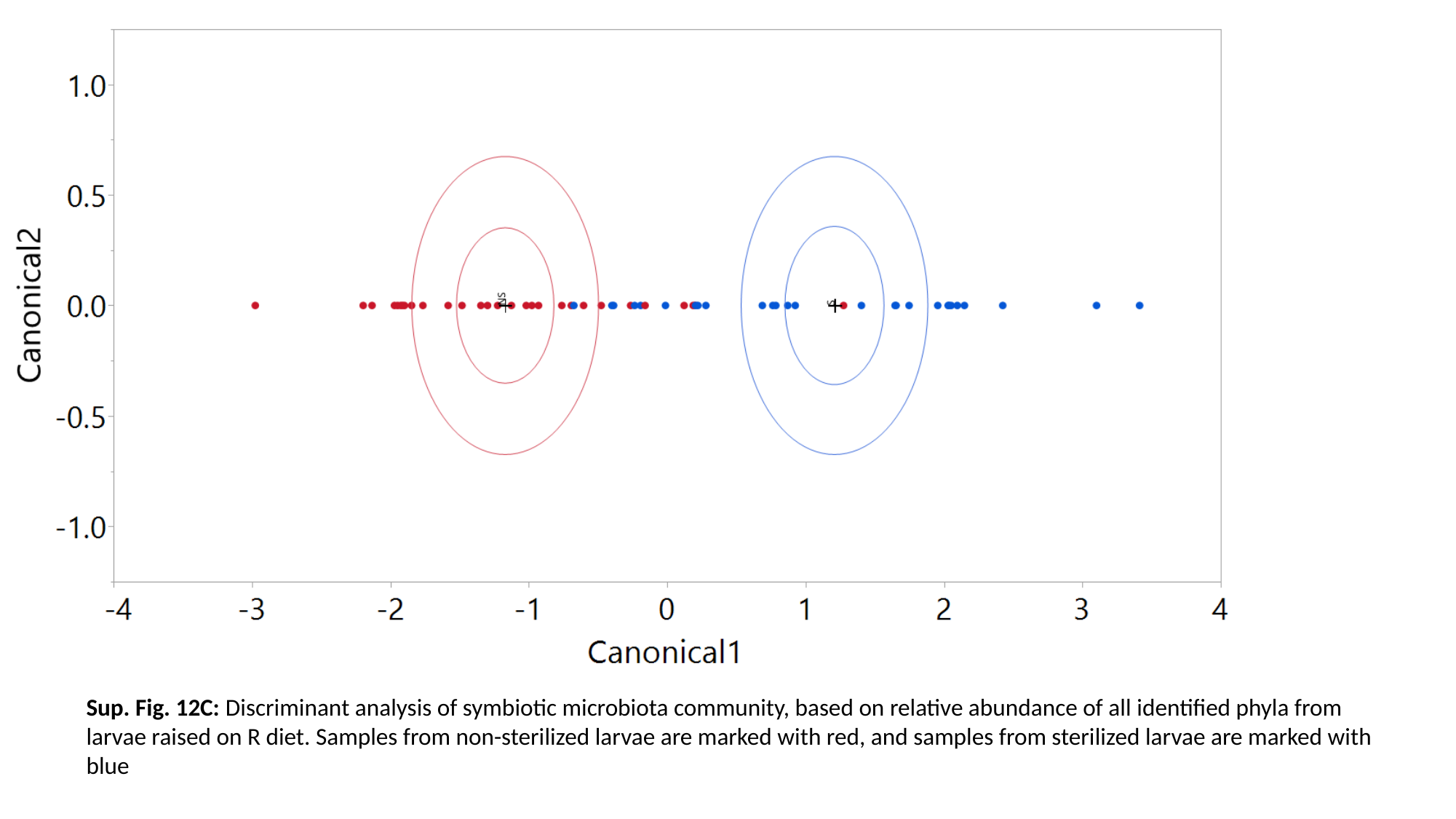

Sup. Fig. 12C: Discriminant analysis of symbiotic microbiota community, based on relative abundance of all identified phyla from larvae raised on R diet. Samples from non-sterilized larvae are marked with red, and samples from sterilized larvae are marked with blue

### Slide 26
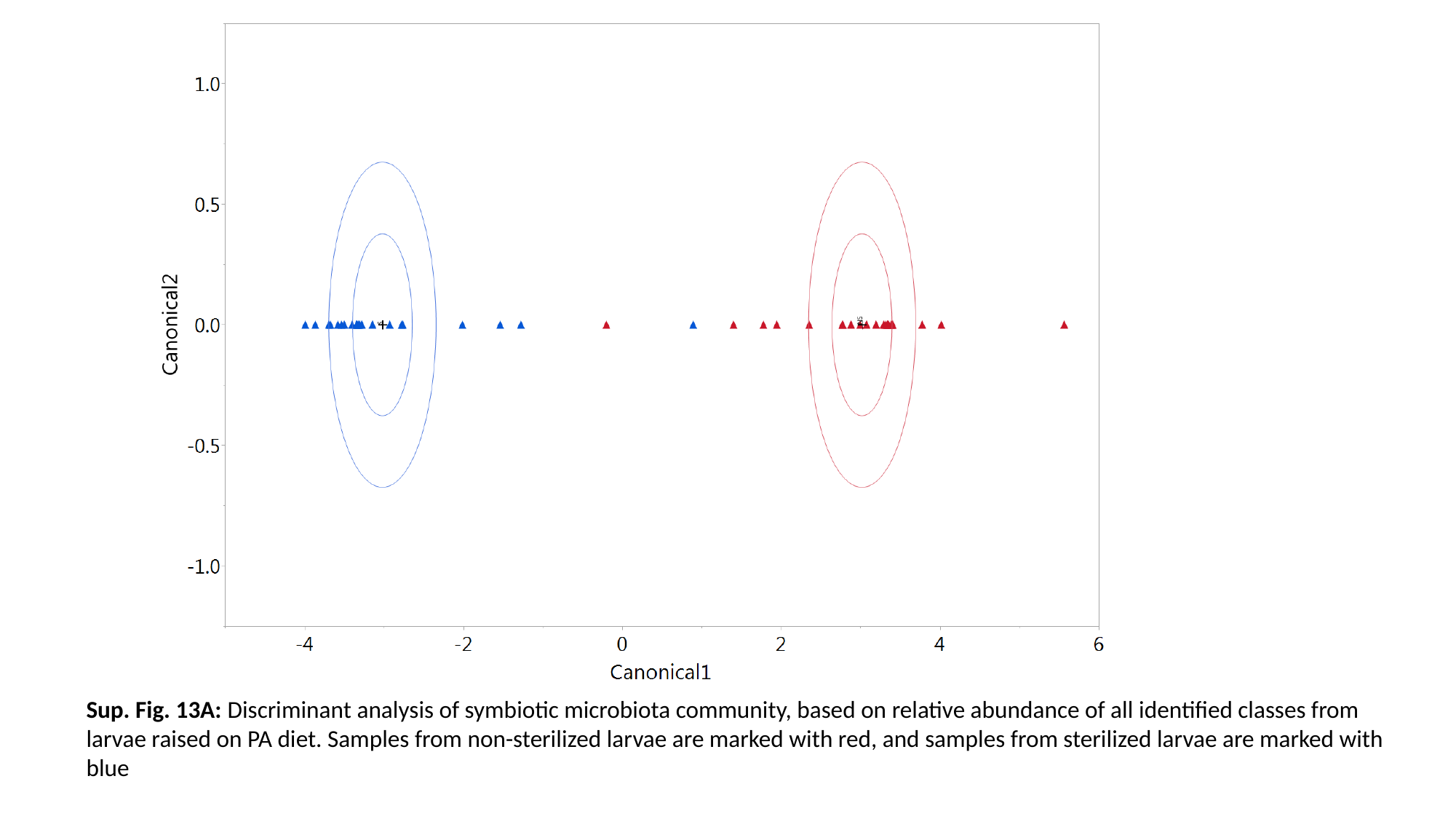

Sup. Fig. 13A: Discriminant analysis of symbiotic microbiota community, based on relative abundance of all identified classes from larvae raised on PA diet. Samples from non-sterilized larvae are marked with red, and samples from sterilized larvae are marked with blue

### Slide 27
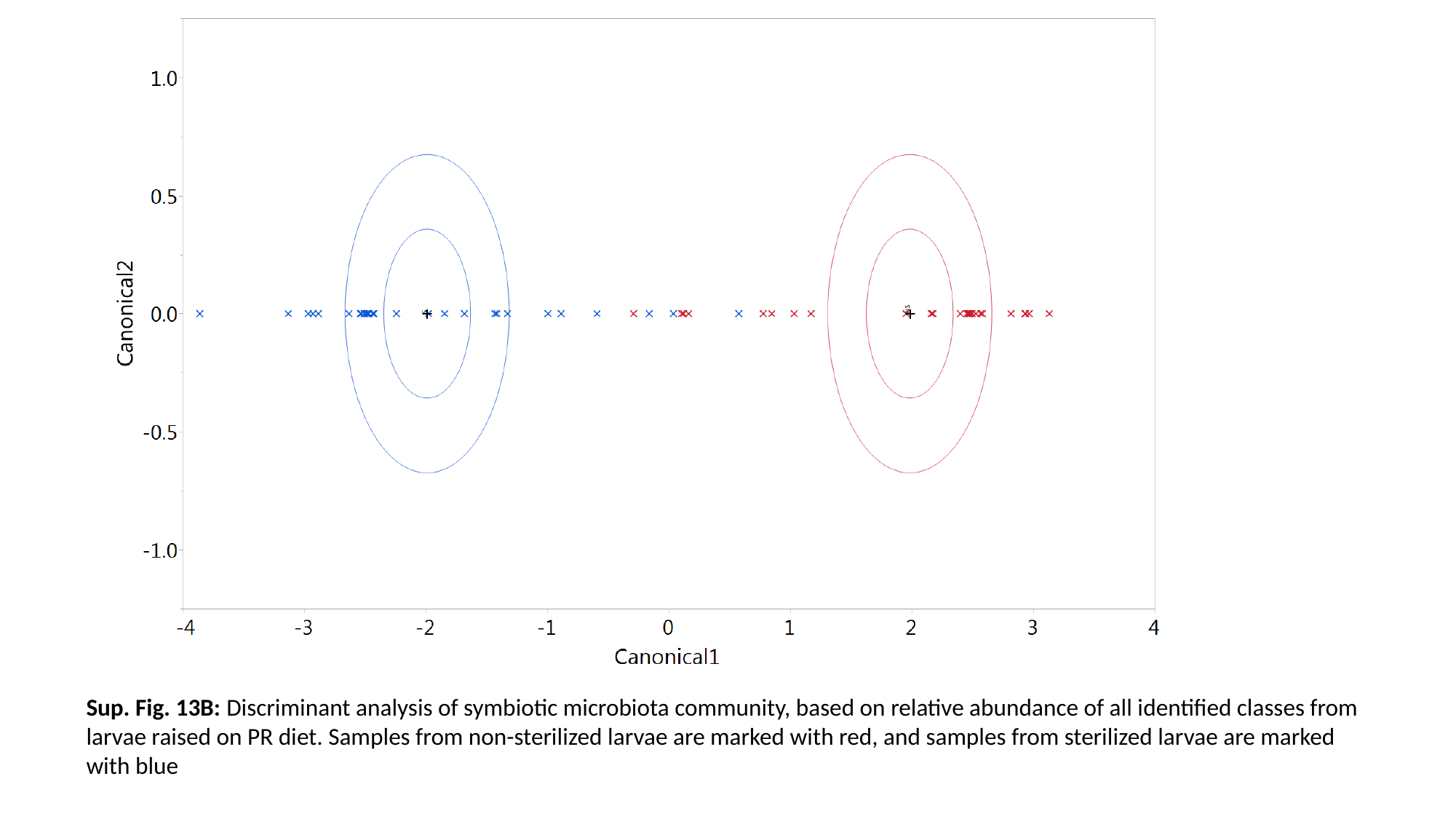

Sup. Fig. 13B: Discriminant analysis of symbiotic microbiota community, based on relative abundance of all identified classes from larvae raised on PR diet. Samples from non-sterilized larvae are marked with red, and samples from sterilized larvae are marked with blue

### Slide 28
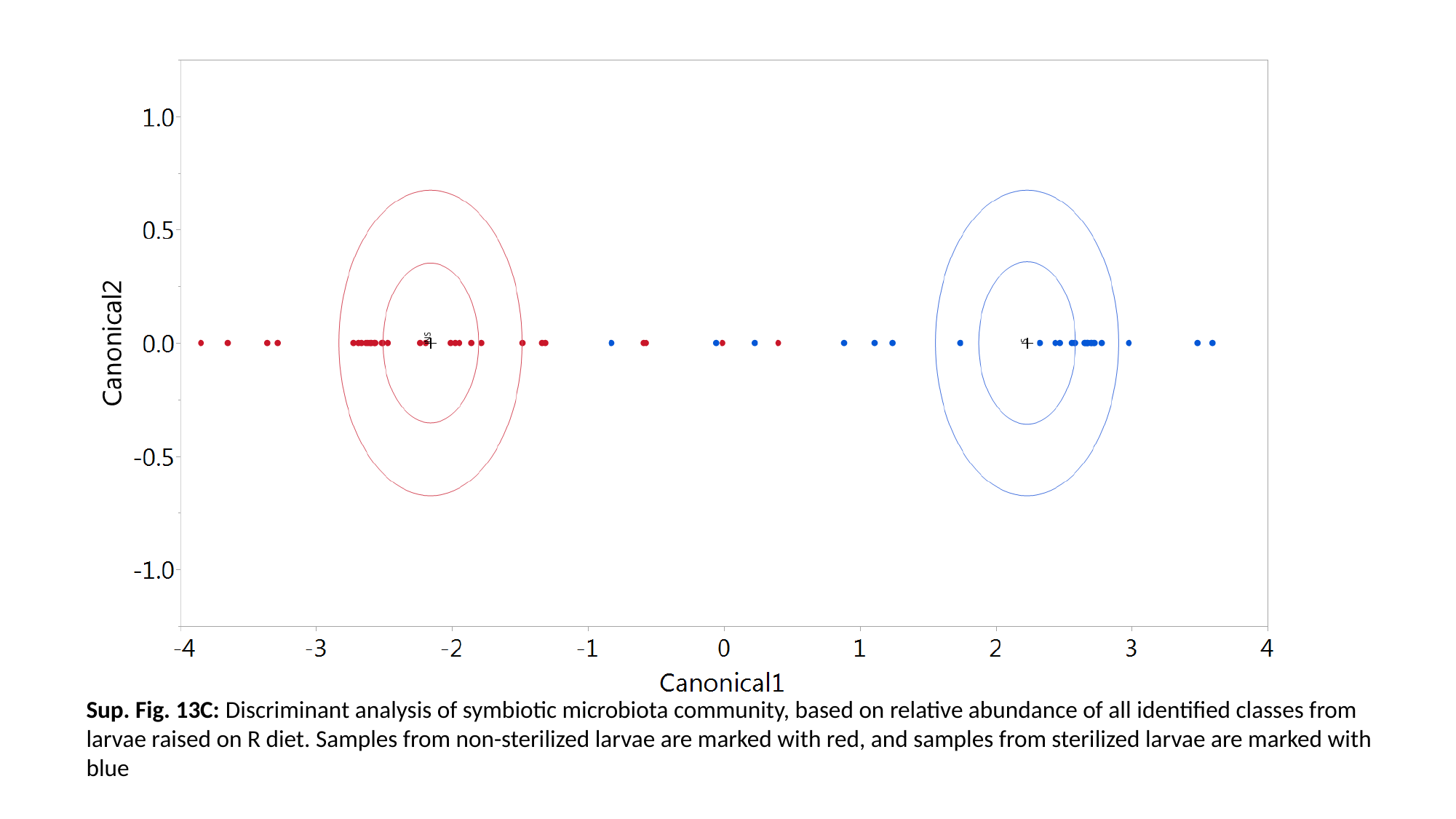

Sup. Fig. 13C: Discriminant analysis of symbiotic microbiota community, based on relative abundance of all identified classes from larvae raised on R diet. Samples from non-sterilized larvae are marked with red, and samples from sterilized larvae are marked with blue

### Slide 29
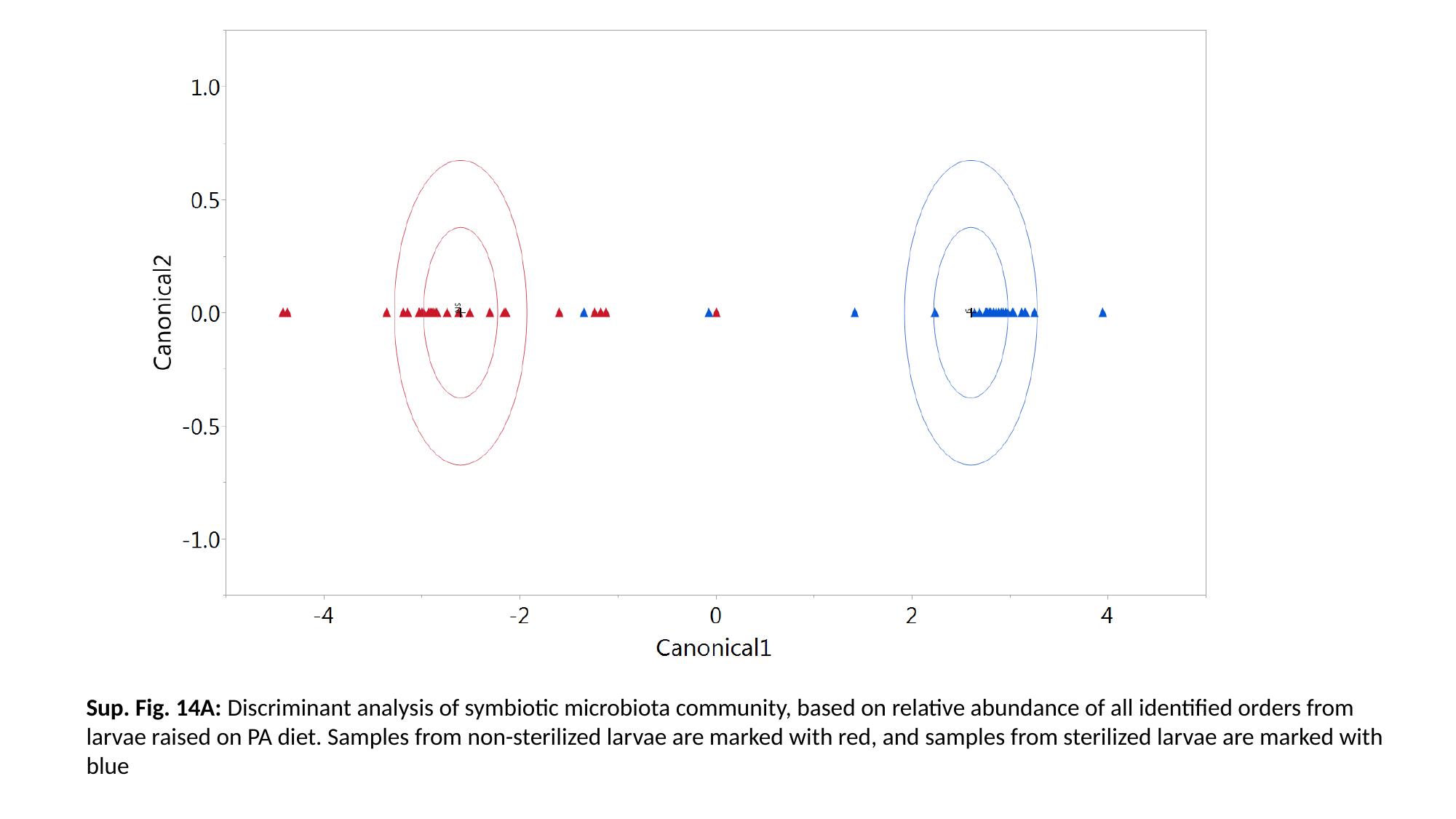

Sup. Fig. 14A: Discriminant analysis of symbiotic microbiota community, based on relative abundance of all identified orders from larvae raised on PA diet. Samples from non-sterilized larvae are marked with red, and samples from sterilized larvae are marked with blue

### Slide 30
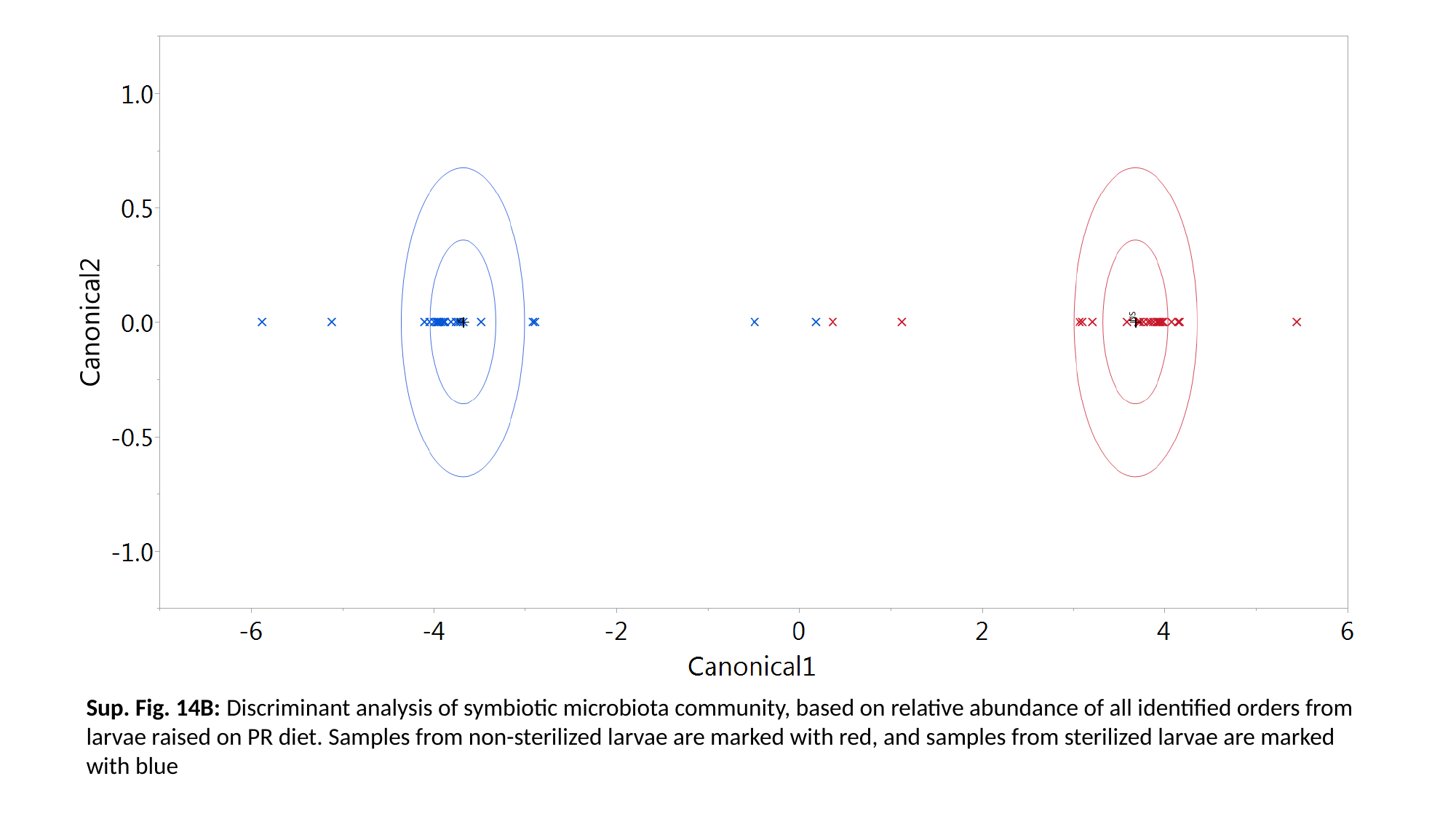

Sup. Fig. 14B: Discriminant analysis of symbiotic microbiota community, based on relative abundance of all identified orders from larvae raised on PR diet. Samples from non-sterilized larvae are marked with red, and samples from sterilized larvae are marked with blue

### Slide 31
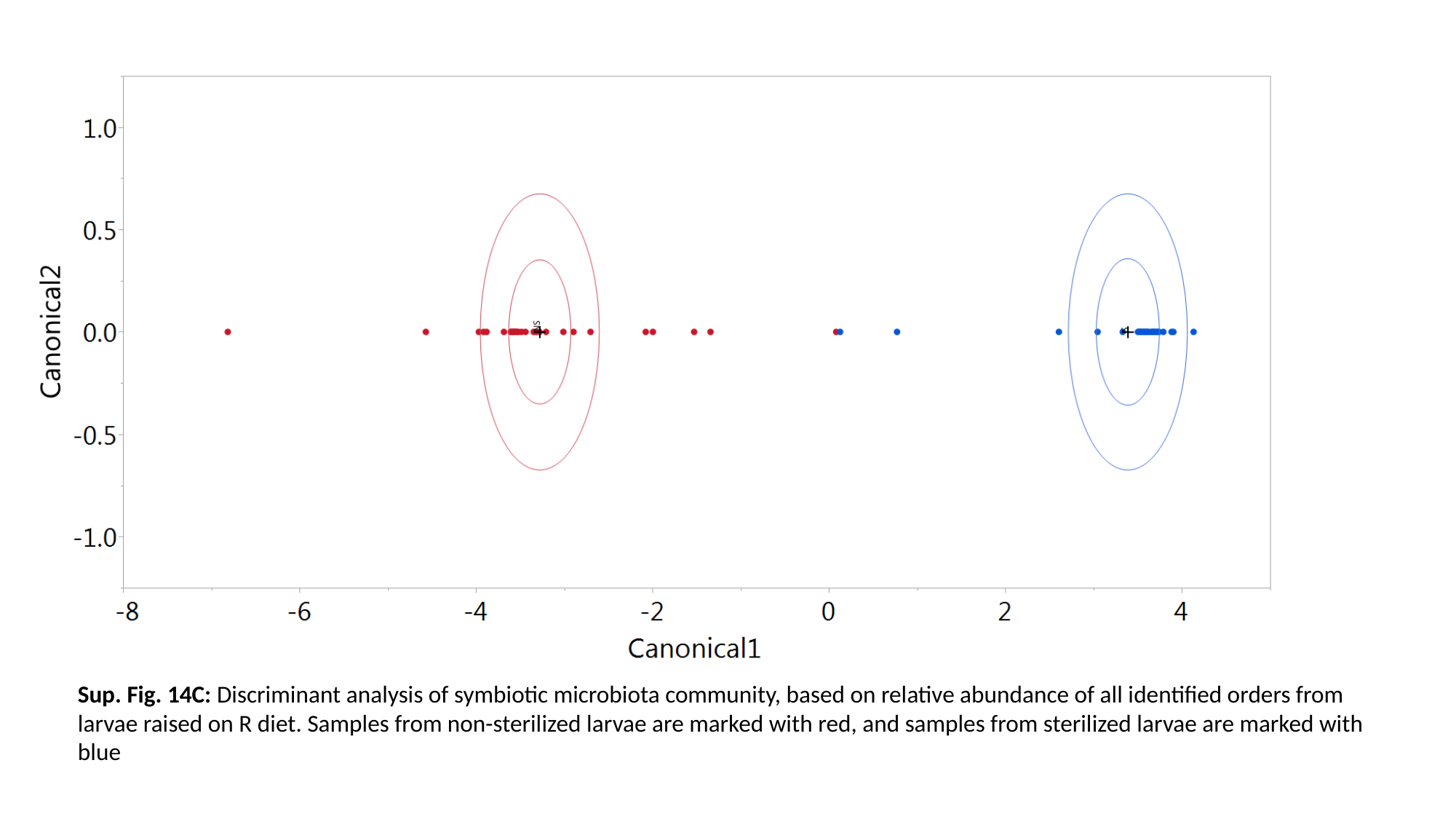

Sup. Fig. 14C: Discriminant analysis of symbiotic microbiota community, based on relative abundance of all identified orders from larvae raised on R diet. Samples from non-sterilized larvae are marked with red, and samples from sterilized larvae are marked with blue

### Slide 32
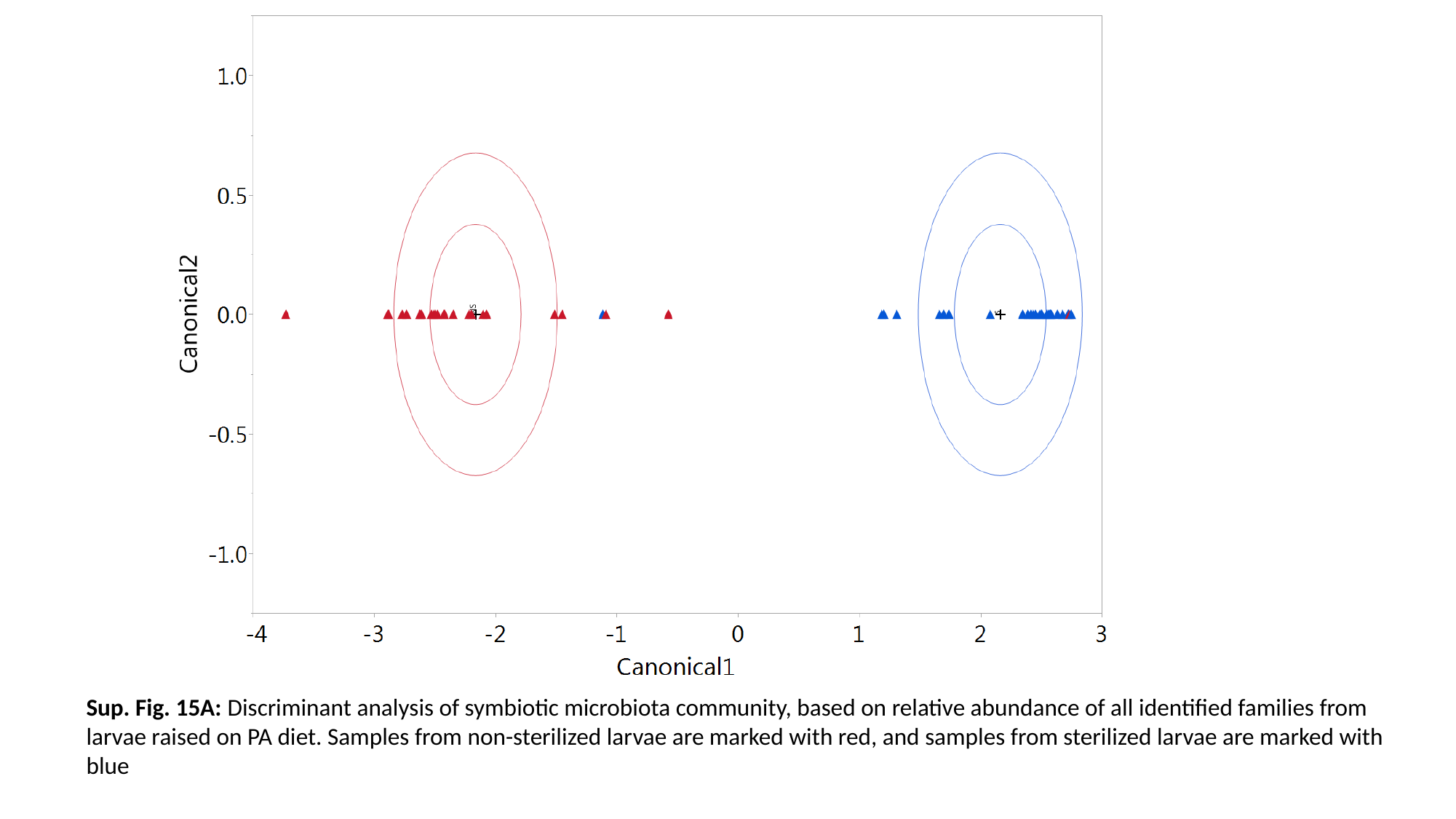

Sup. Fig. 15A: Discriminant analysis of symbiotic microbiota community, based on relative abundance of all identified families from larvae raised on PA diet. Samples from non-sterilized larvae are marked with red, and samples from sterilized larvae are marked with blue

### Slide 33
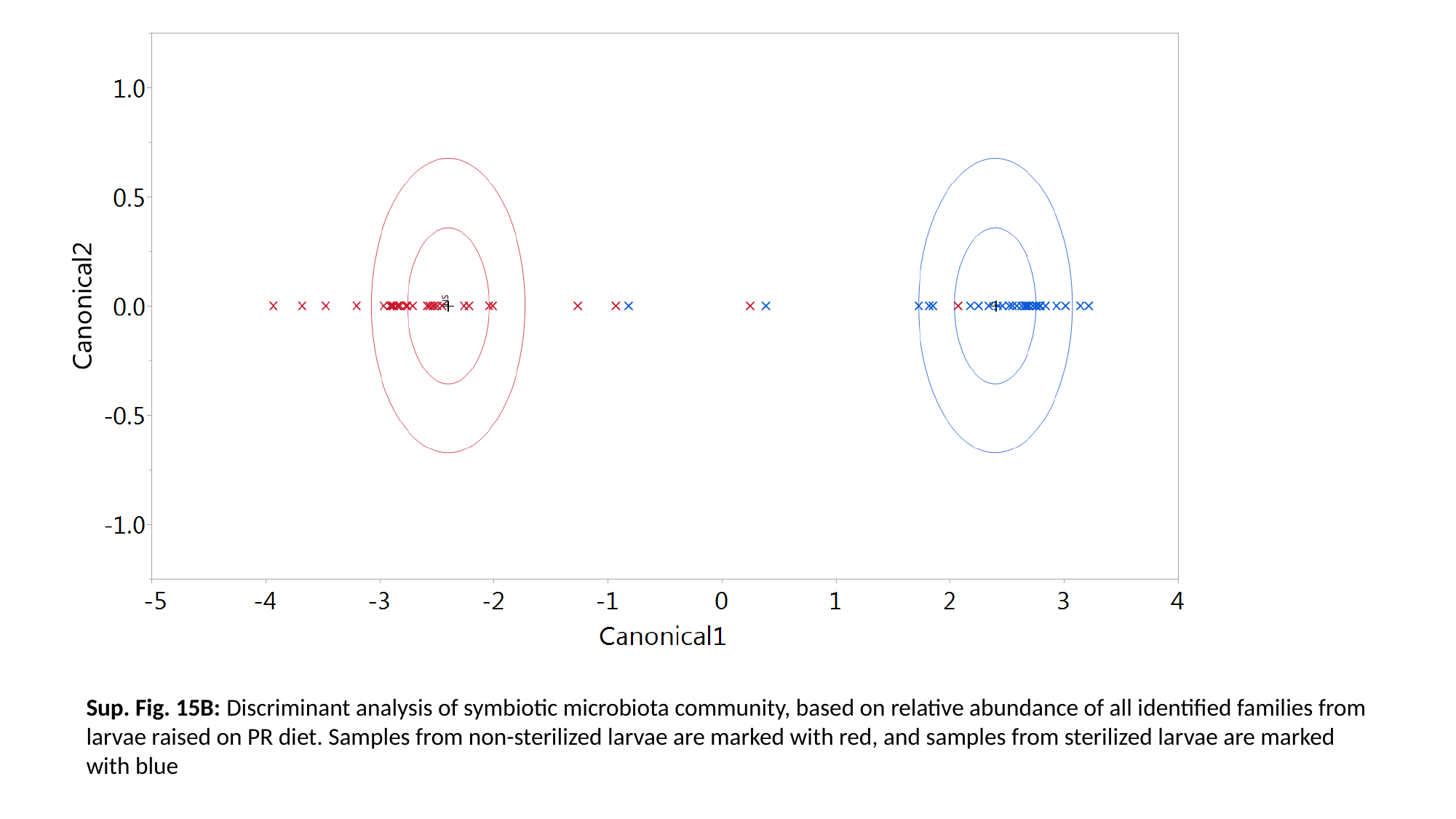

Sup. Fig. 15B: Discriminant analysis of symbiotic microbiota community, based on relative abundance of all identified families from larvae raised on PR diet. Samples from non-sterilized larvae are marked with red, and samples from sterilized larvae are marked with blue

### Slide 34
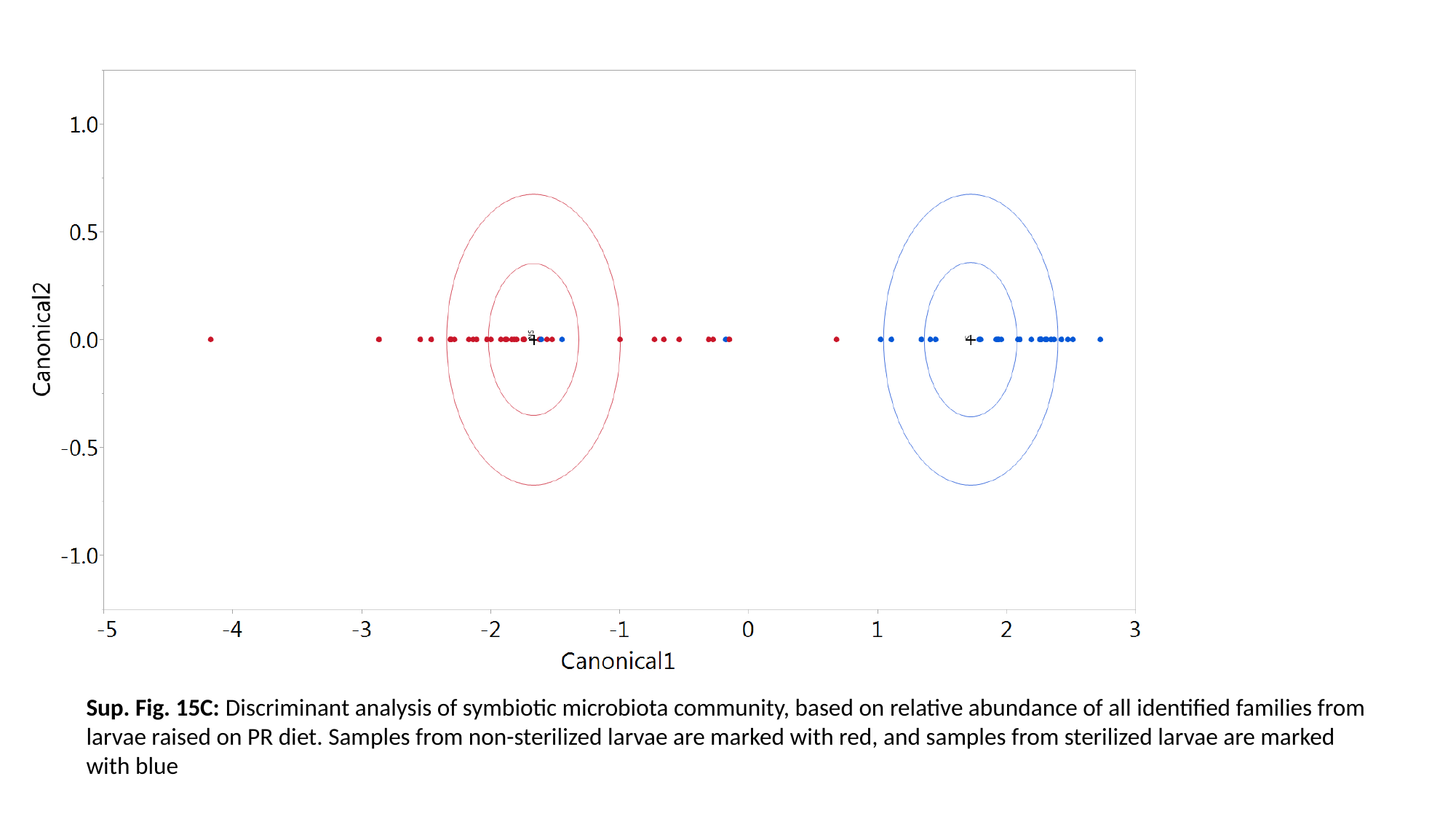

Sup. Fig. 15C: Discriminant analysis of symbiotic microbiota community, based on relative abundance of all identified families from larvae raised on PR diet. Samples from non-sterilized larvae are marked with red, and samples from sterilized larvae are marked with blue

### Slide 35
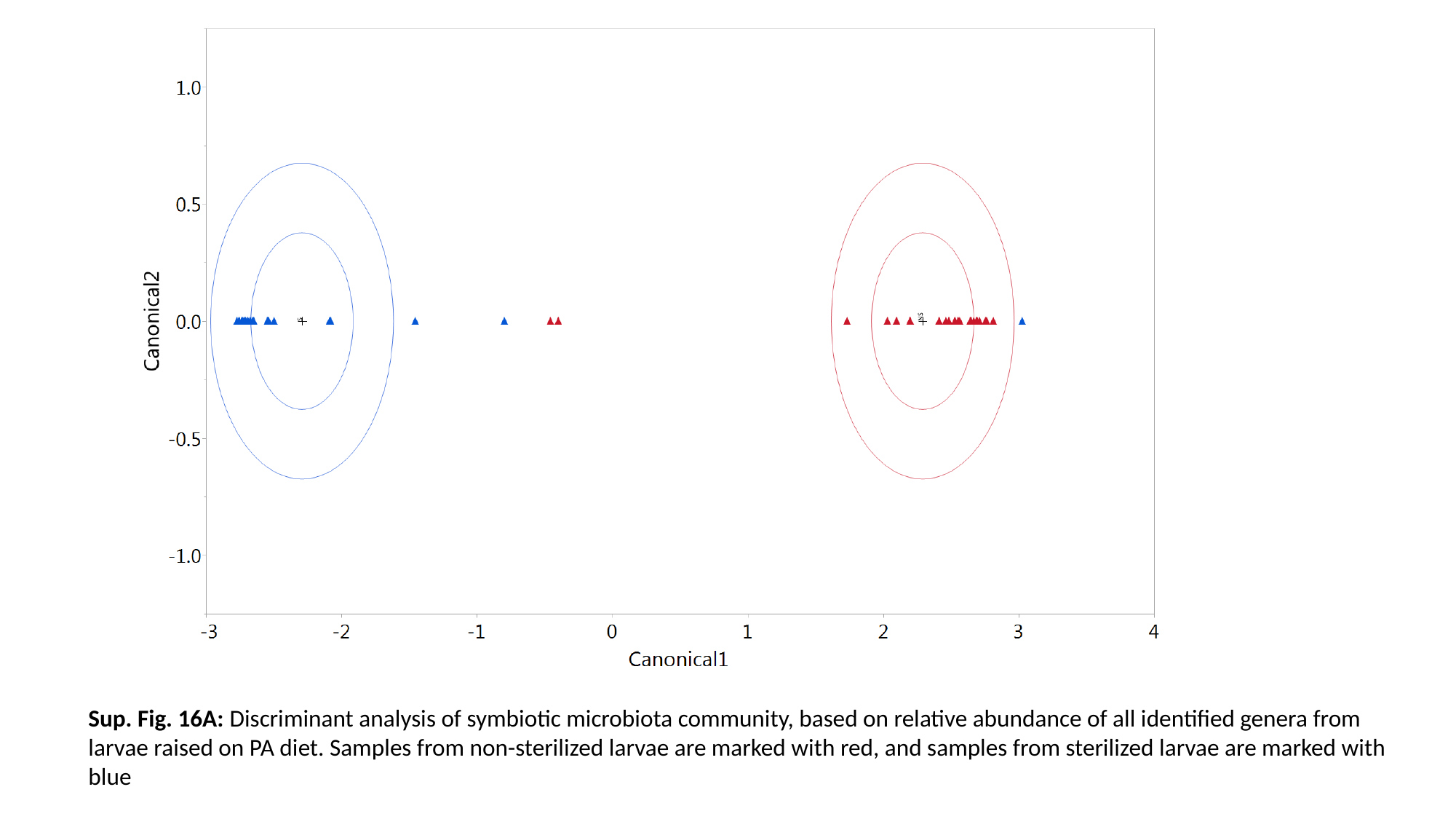

Sup. Fig. 16A: Discriminant analysis of symbiotic microbiota community, based on relative abundance of all identified genera from larvae raised on PA diet. Samples from non-sterilized larvae are marked with red, and samples from sterilized larvae are marked with blue

### Slide 36
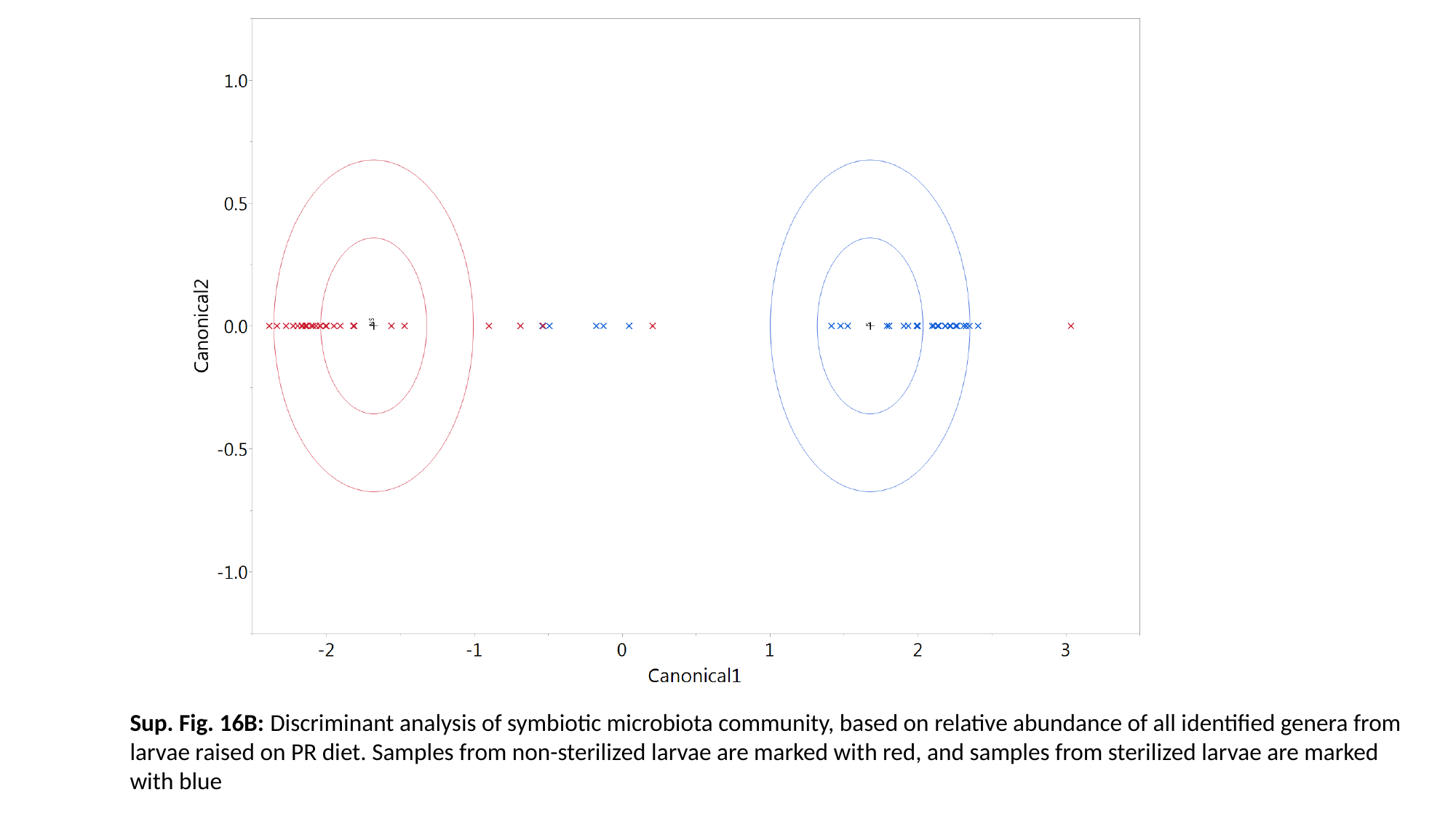

Sup. Fig. 16B: Discriminant analysis of symbiotic microbiota community, based on relative abundance of all identified genera from larvae raised on PR diet. Samples from non-sterilized larvae are marked with red, and samples from sterilized larvae are marked with blue

### Slide 37
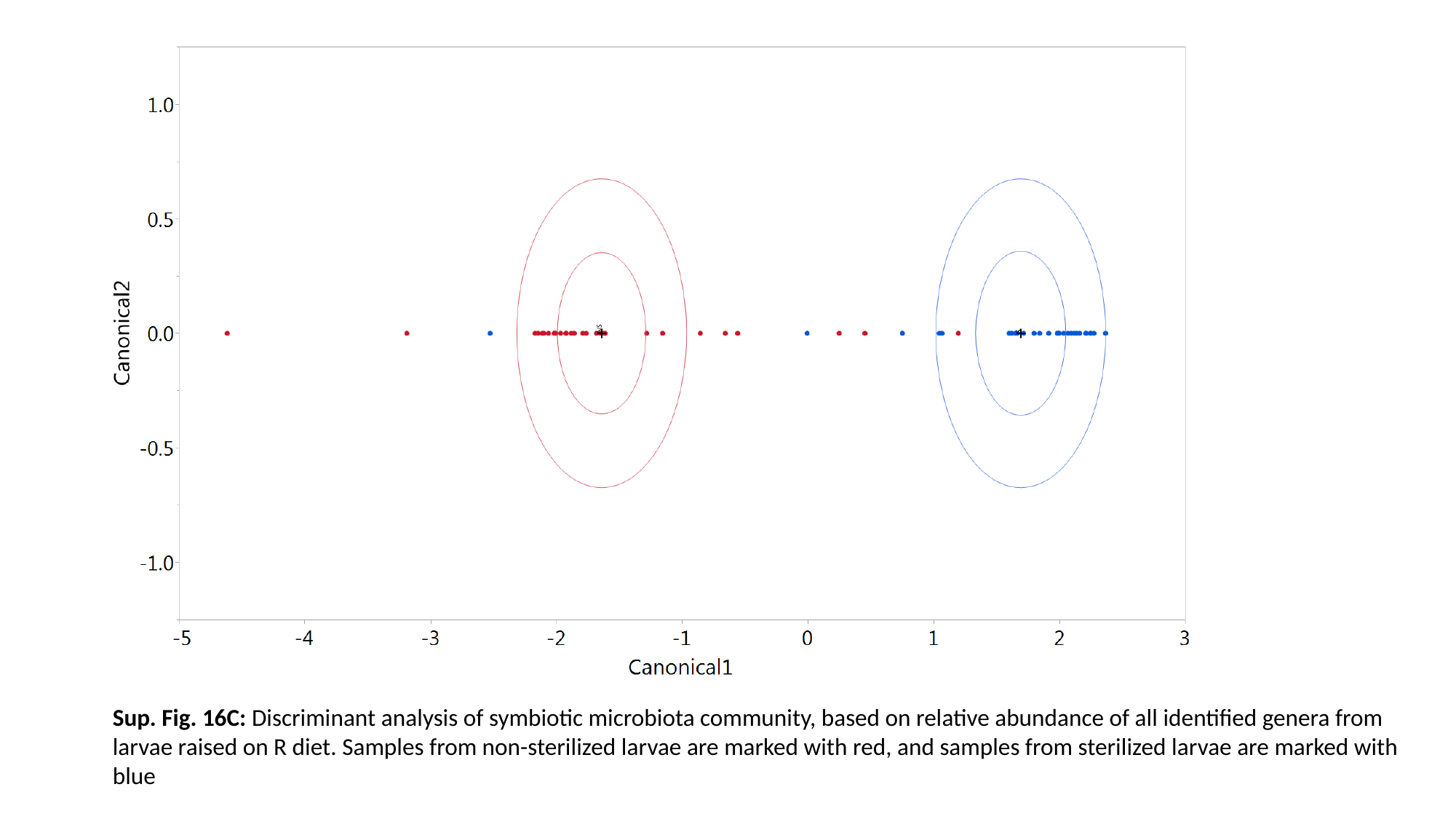

Sup. Fig. 16C: Discriminant analysis of symbiotic microbiota community, based on relative abundance of all identified genera from larvae raised on R diet. Samples from non-sterilized larvae are marked with red, and samples from sterilized larvae are marked with blue

### Slide 38
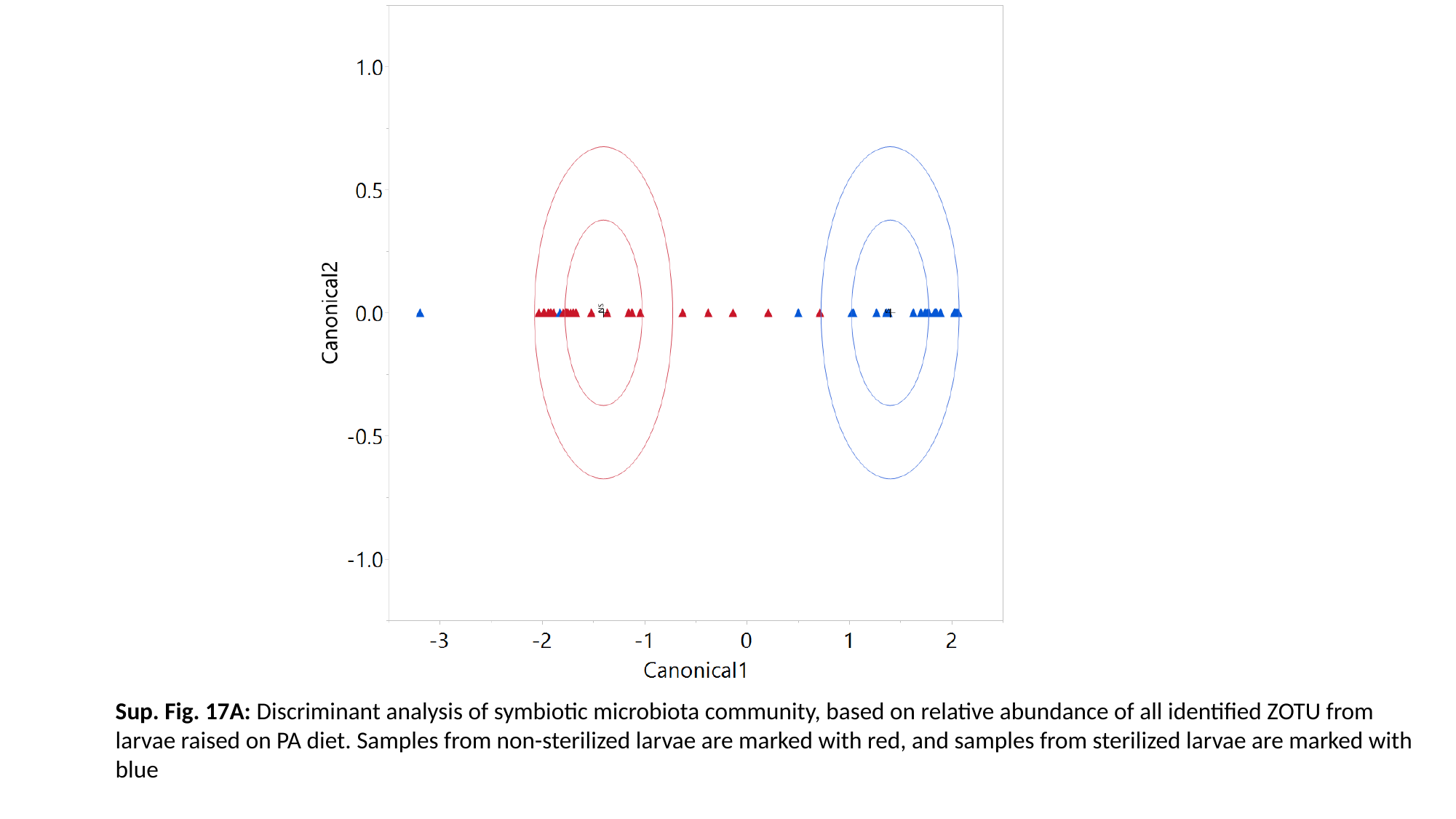

Sup. Fig. 17A: Discriminant analysis of symbiotic microbiota community, based on relative abundance of all identified ZOTU from larvae raised on PA diet. Samples from non-sterilized larvae are marked with red, and samples from sterilized larvae are marked with blue

### Slide 39
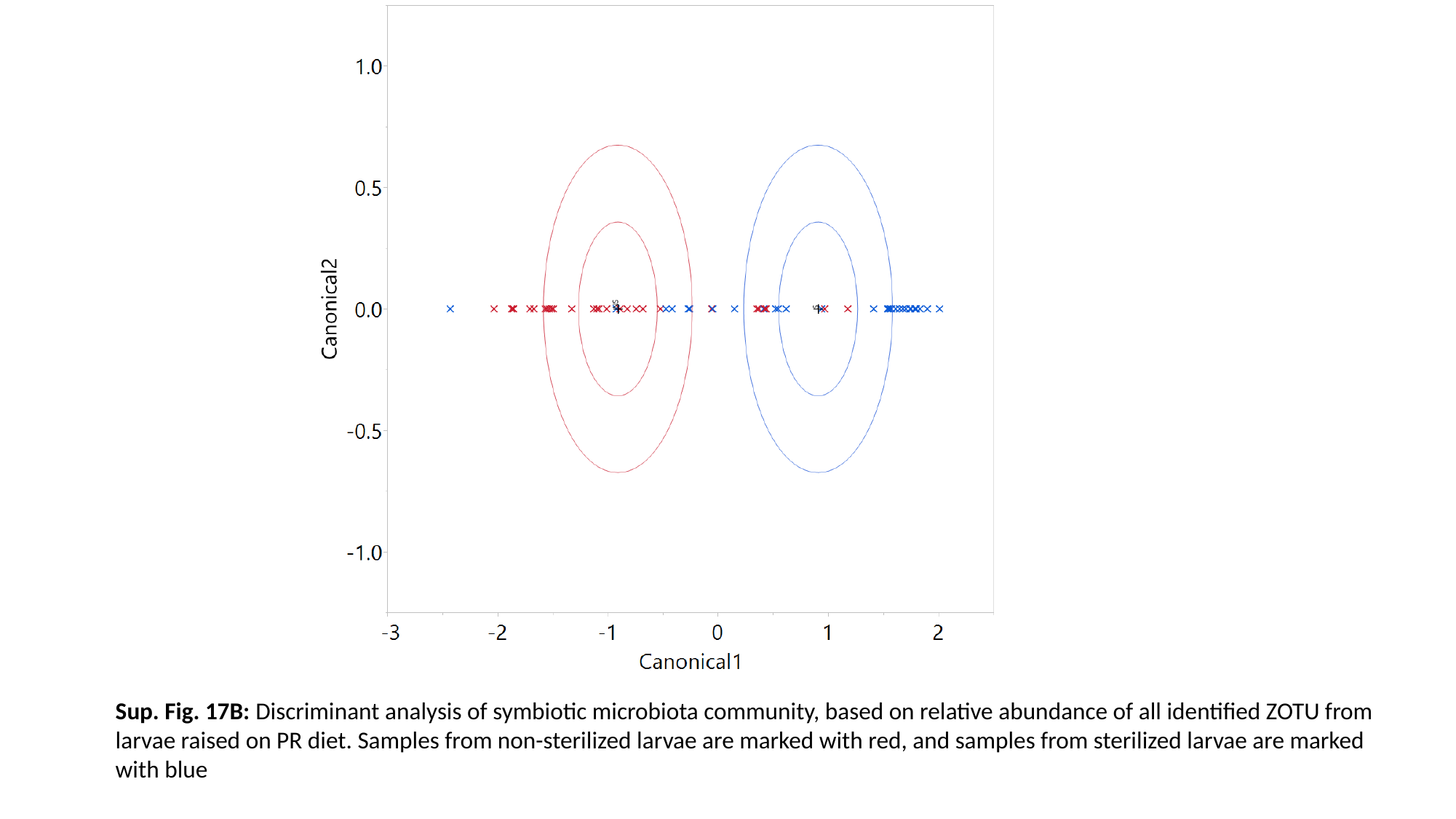

Sup. Fig. 17B: Discriminant analysis of symbiotic microbiota community, based on relative abundance of all identified ZOTU from larvae raised on PR diet. Samples from non-sterilized larvae are marked with red, and samples from sterilized larvae are marked with blue

### Slide 40
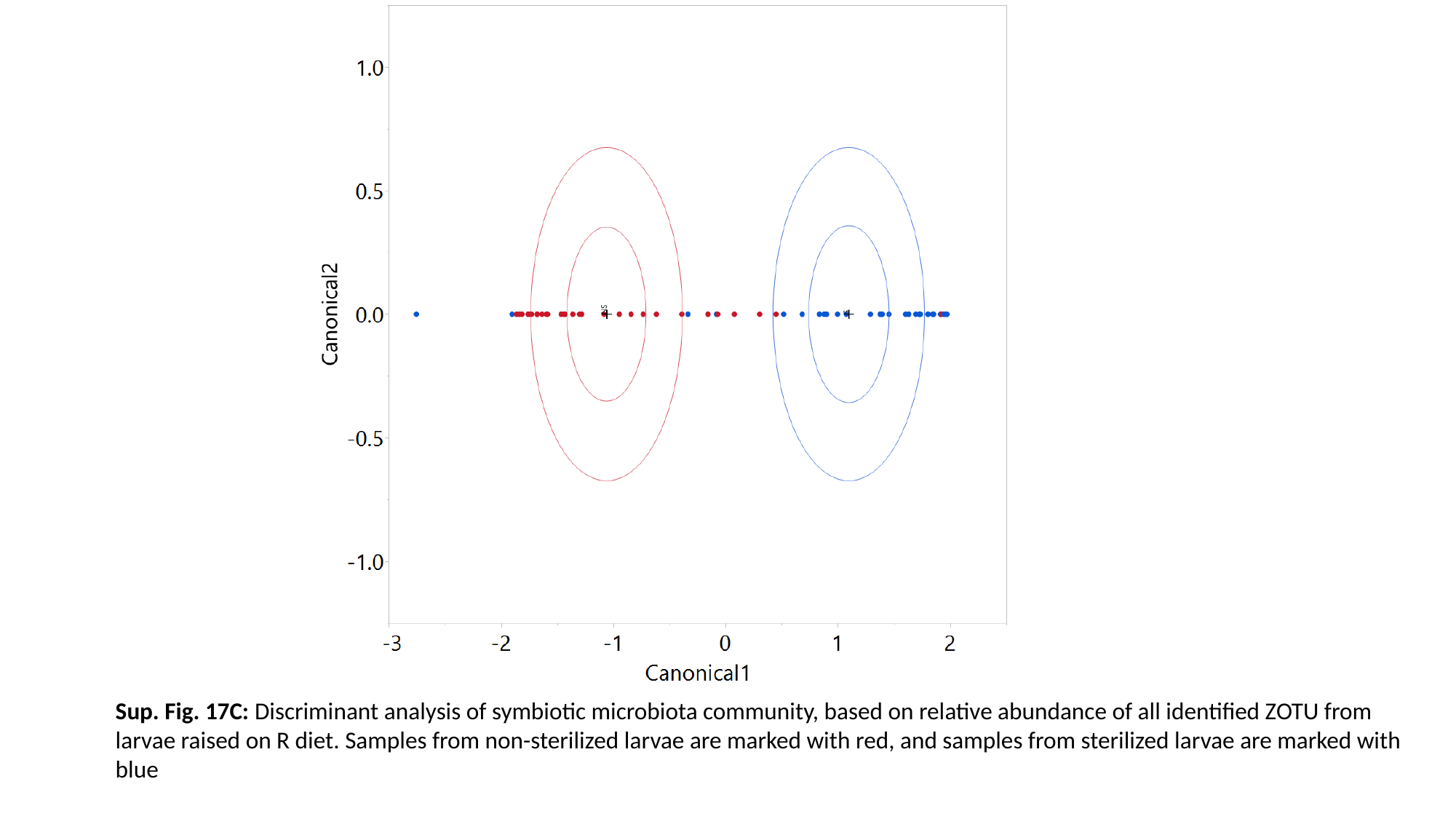

Sup. Fig. 17C: Discriminant analysis of symbiotic microbiota community, based on relative abundance of all identified ZOTU from larvae raised on R diet. Samples from non-sterilized larvae are marked with red, and samples from sterilized larvae are marked with blue

### Slide 41
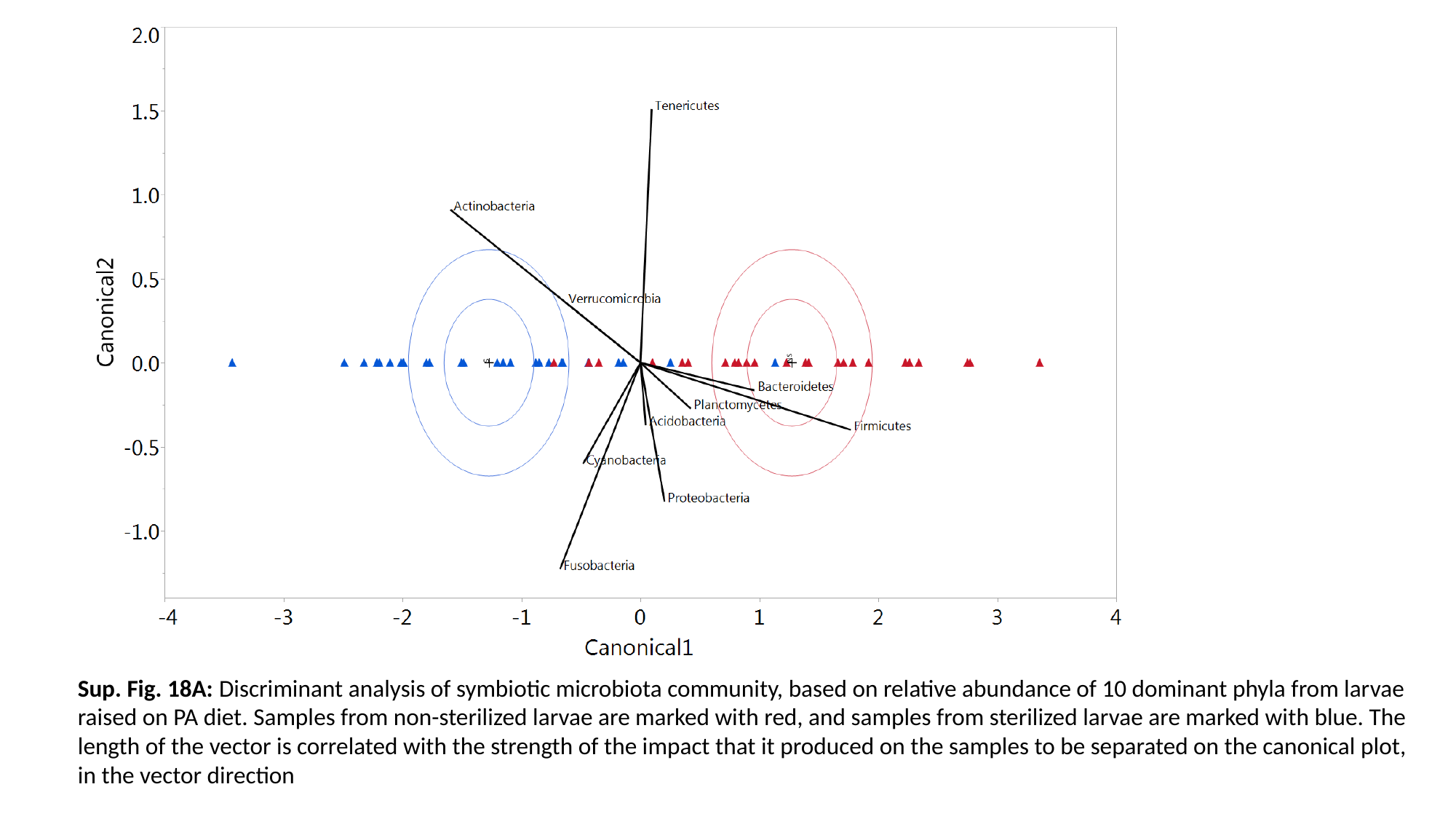

Sup. Fig. 18A: Discriminant analysis of symbiotic microbiota community, based on relative abundance of 10 dominant phyla from larvae raised on PA diet. Samples from non-sterilized larvae are marked with red, and samples from sterilized larvae are marked with blue. The length of the vector is correlated with the strength of the impact that it produced on the samples to be separated on the canonical plot, in the vector direction

### Slide 42
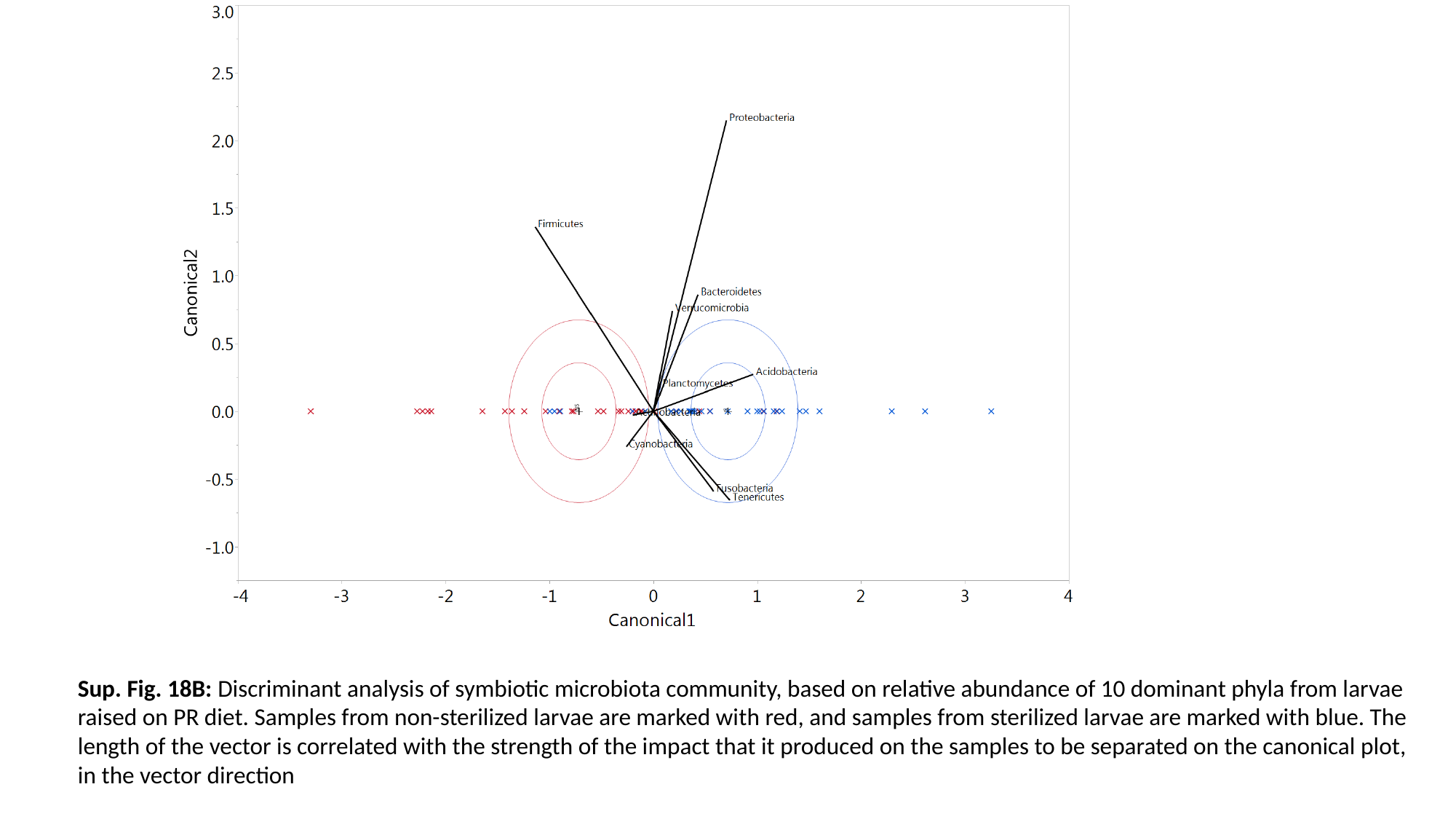

Sup. Fig. 18B: Discriminant analysis of symbiotic microbiota community, based on relative abundance of 10 dominant phyla from larvae raised on PR diet. Samples from non-sterilized larvae are marked with red, and samples from sterilized larvae are marked with blue. The length of the vector is correlated with the strength of the impact that it produced on the samples to be separated on the canonical plot, in the vector direction

### Slide 43
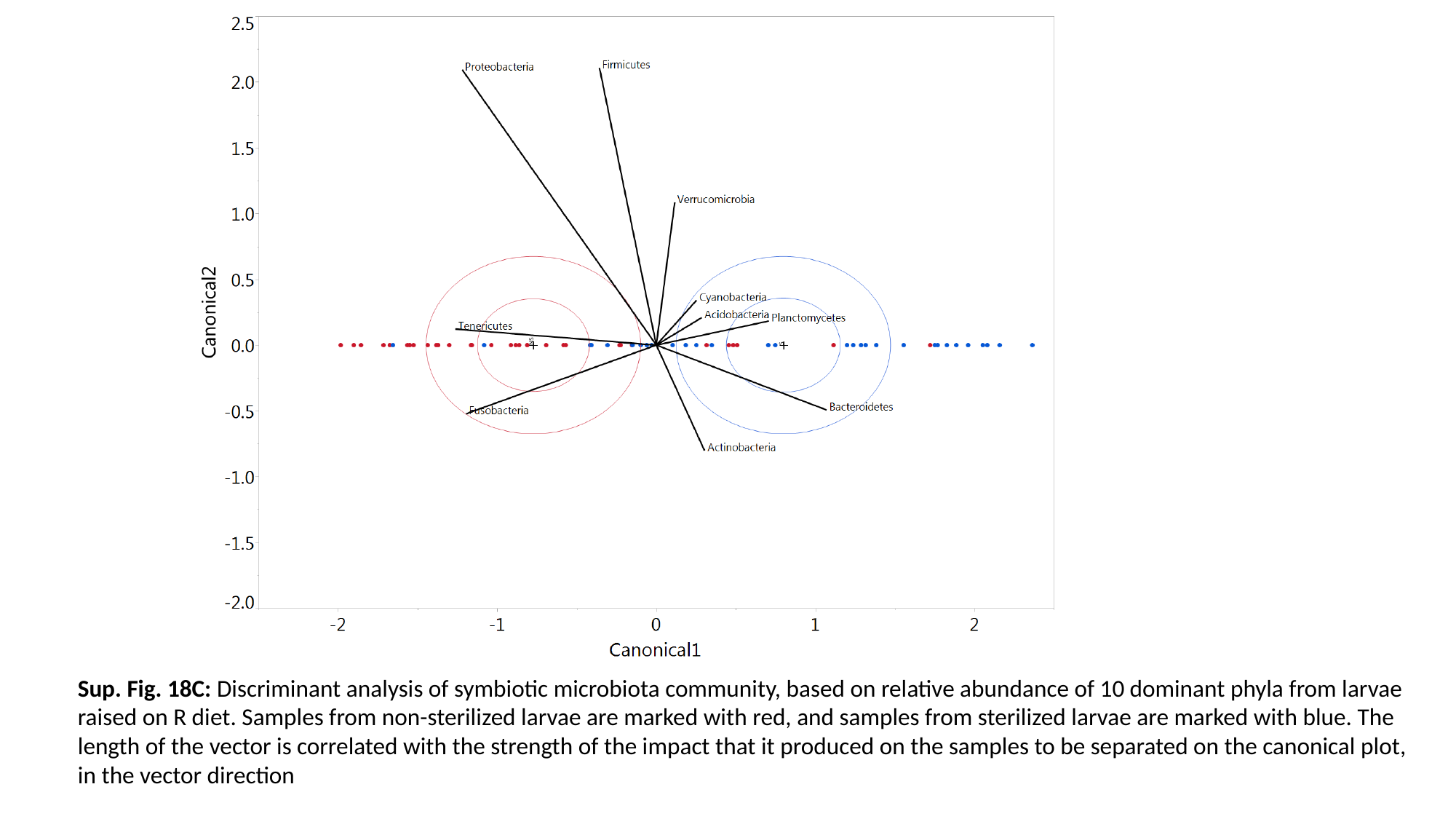

Sup. Fig. 18C: Discriminant analysis of symbiotic microbiota community, based on relative abundance of 10 dominant phyla from larvae raised on R diet. Samples from non-sterilized larvae are marked with red, and samples from sterilized larvae are marked with blue. The length of the vector is correlated with the strength of the impact that it produced on the samples to be separated on the canonical plot, in the vector direction

### Slide 44
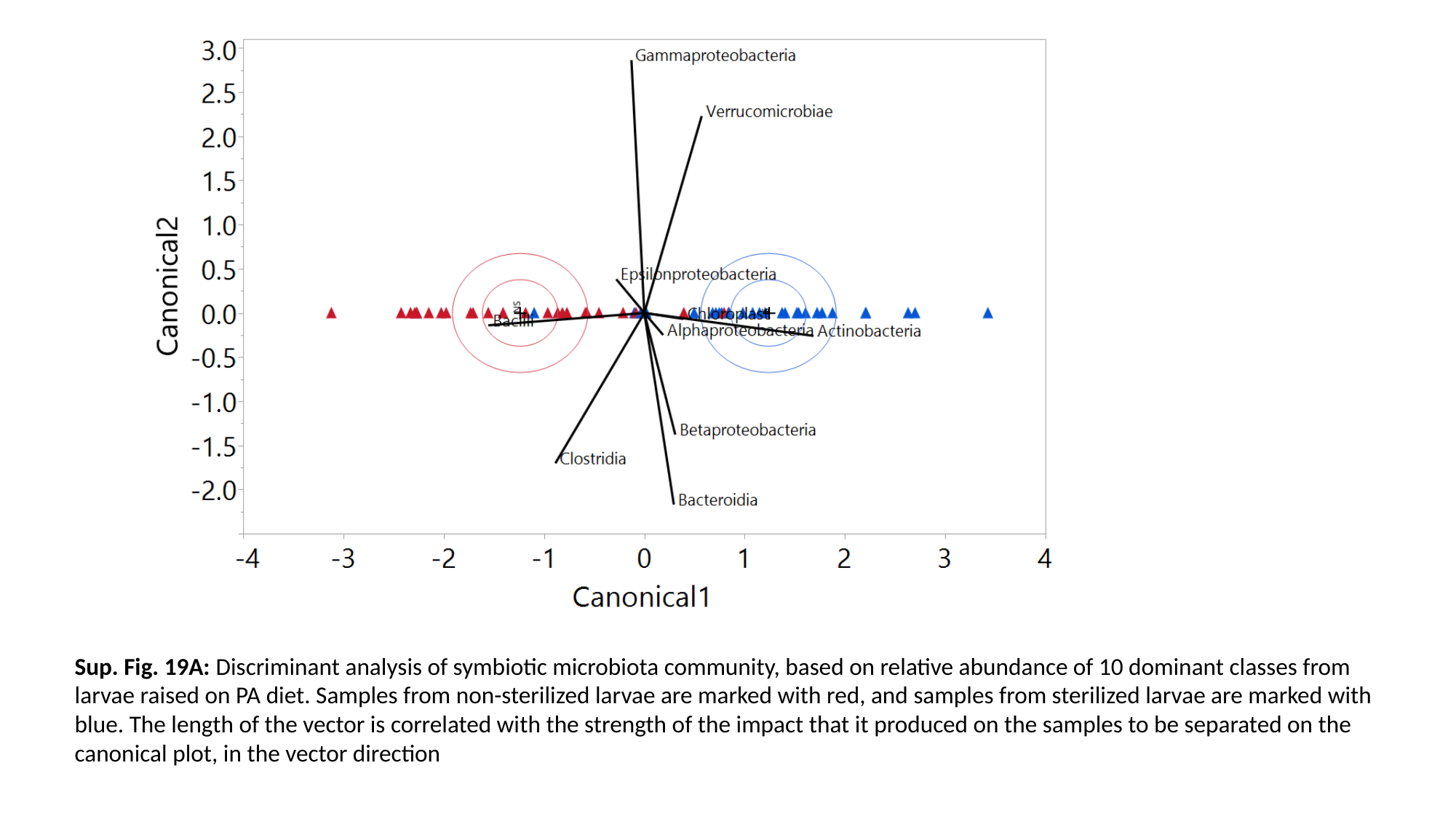

Sup. Fig. 19A: Discriminant analysis of symbiotic microbiota community, based on relative abundance of 10 dominant classes from larvae raised on PA diet. Samples from non-sterilized larvae are marked with red, and samples from sterilized larvae are marked with blue. The length of the vector is correlated with the strength of the impact that it produced on the samples to be separated on the canonical plot, in the vector direction

### Slide 45
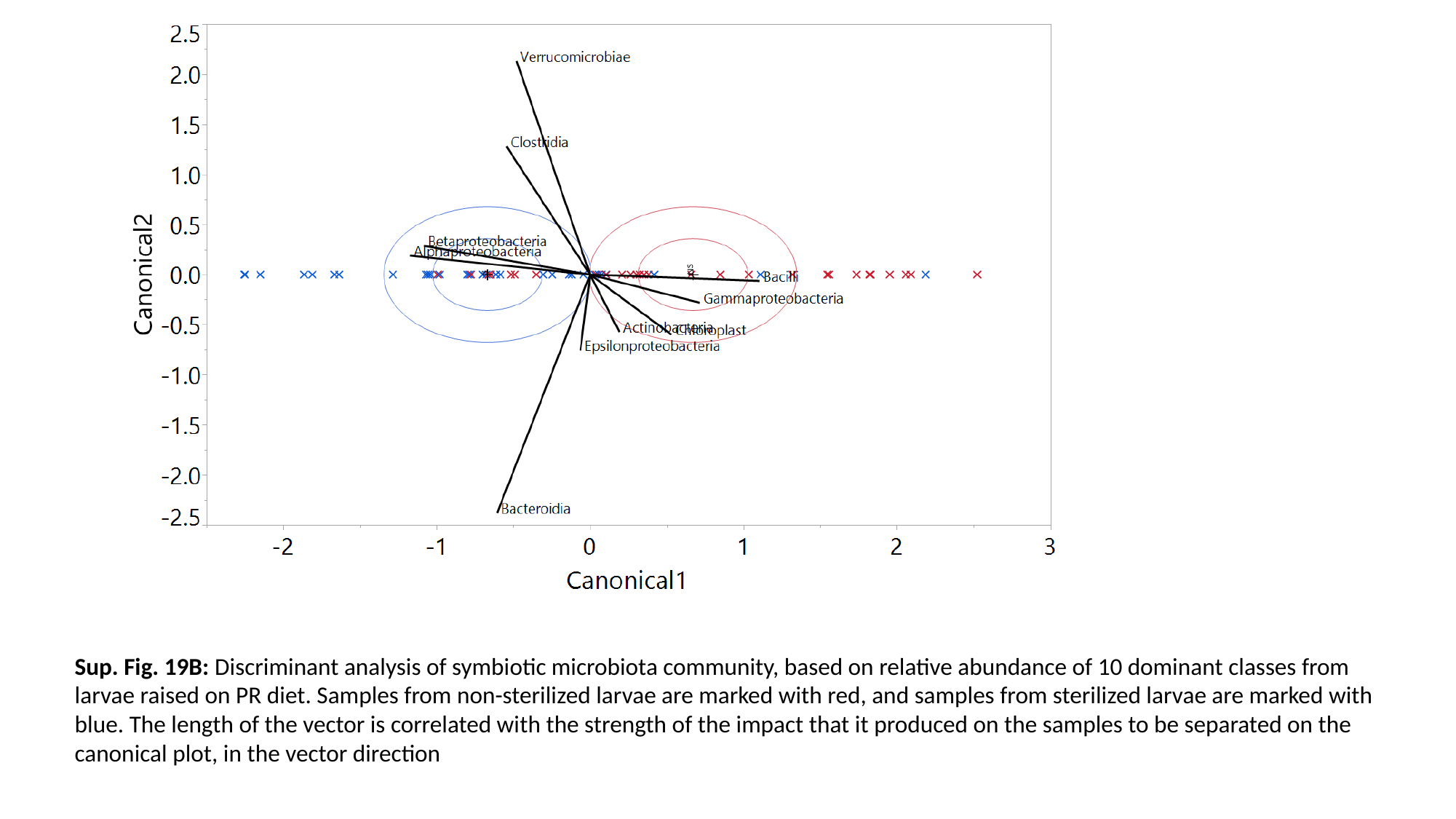

Sup. Fig. 19B: Discriminant analysis of symbiotic microbiota community, based on relative abundance of 10 dominant classes from larvae raised on PR diet. Samples from non-sterilized larvae are marked with red, and samples from sterilized larvae are marked with blue. The length of the vector is correlated with the strength of the impact that it produced on the samples to be separated on the canonical plot, in the vector direction

### Slide 46
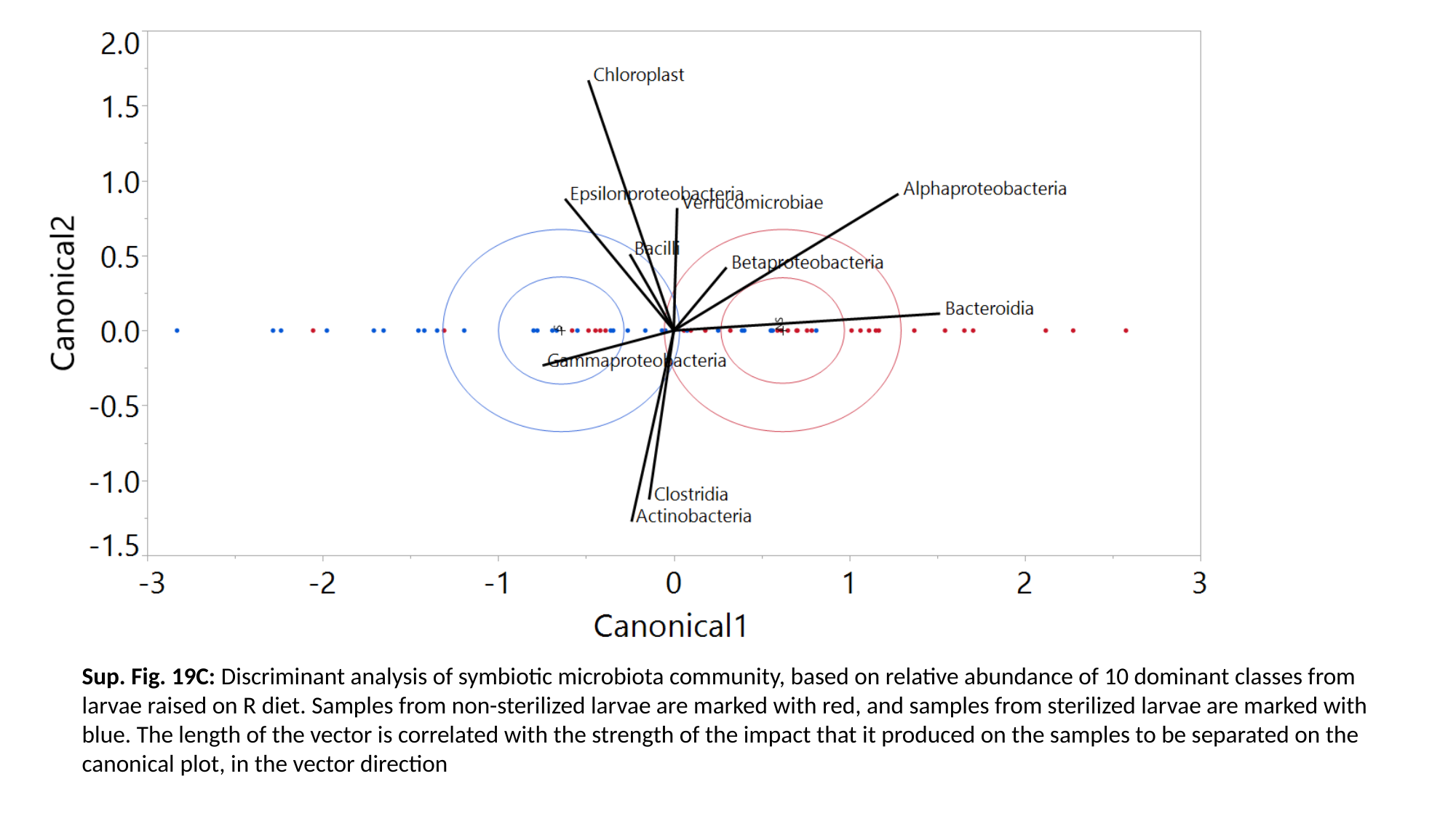

Sup. Fig. 19C: Discriminant analysis of symbiotic microbiota community, based on relative abundance of 10 dominant classes from larvae raised on R diet. Samples from non-sterilized larvae are marked with red, and samples from sterilized larvae are marked with blue. The length of the vector is correlated with the strength of the impact that it produced on the samples to be separated on the canonical plot, in the vector direction

### Slide 47
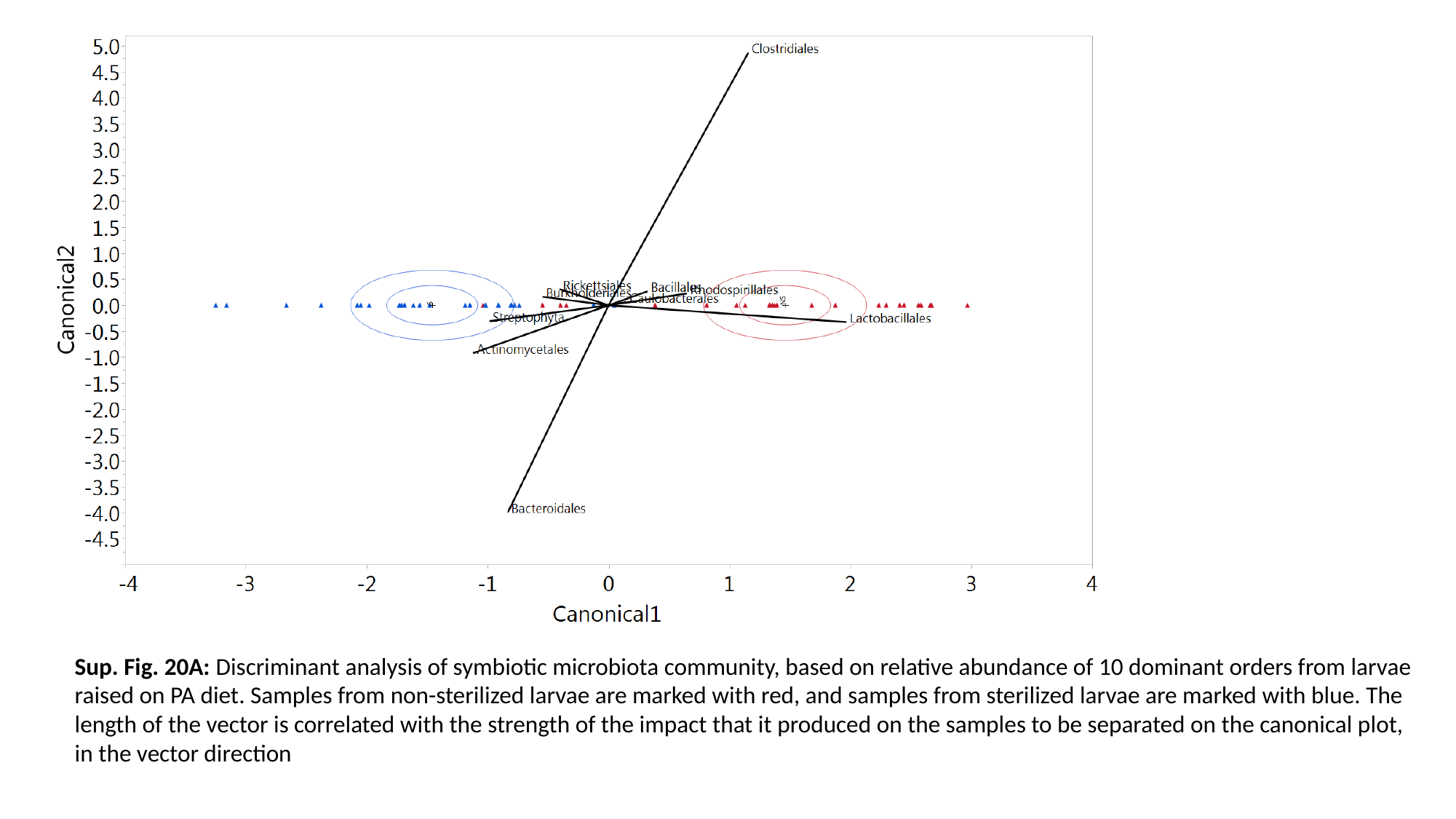

Sup. Fig. 20A: Discriminant analysis of symbiotic microbiota community, based on relative abundance of 10 dominant orders from larvae raised on PA diet. Samples from non-sterilized larvae are marked with red, and samples from sterilized larvae are marked with blue. The length of the vector is correlated with the strength of the impact that it produced on the samples to be separated on the canonical plot, in the vector direction

### Slide 48
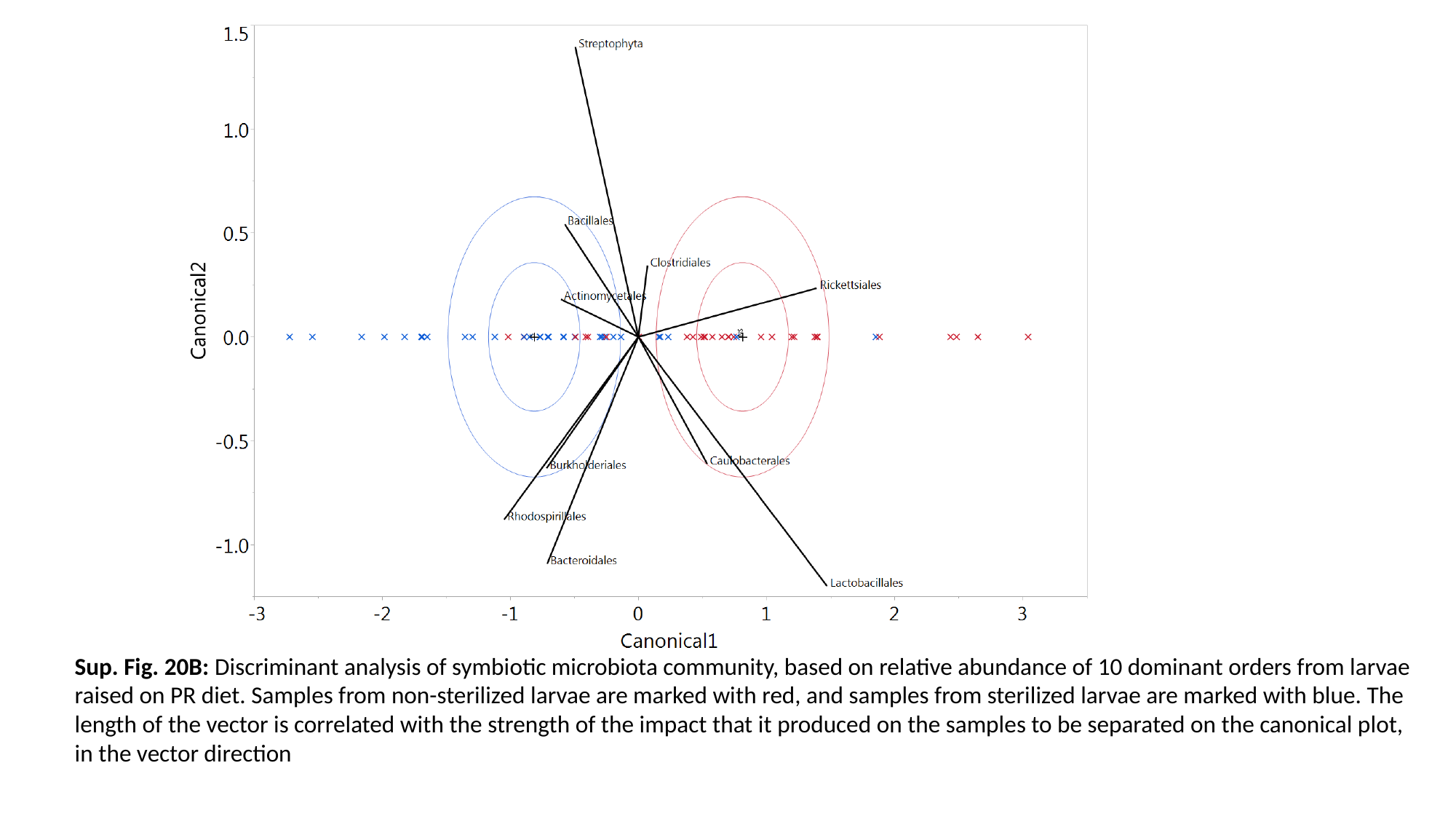

Sup. Fig. 20B: Discriminant analysis of symbiotic microbiota community, based on relative abundance of 10 dominant orders from larvae raised on PR diet. Samples from non-sterilized larvae are marked with red, and samples from sterilized larvae are marked with blue. The length of the vector is correlated with the strength of the impact that it produced on the samples to be separated on the canonical plot, in the vector direction

### Slide 49
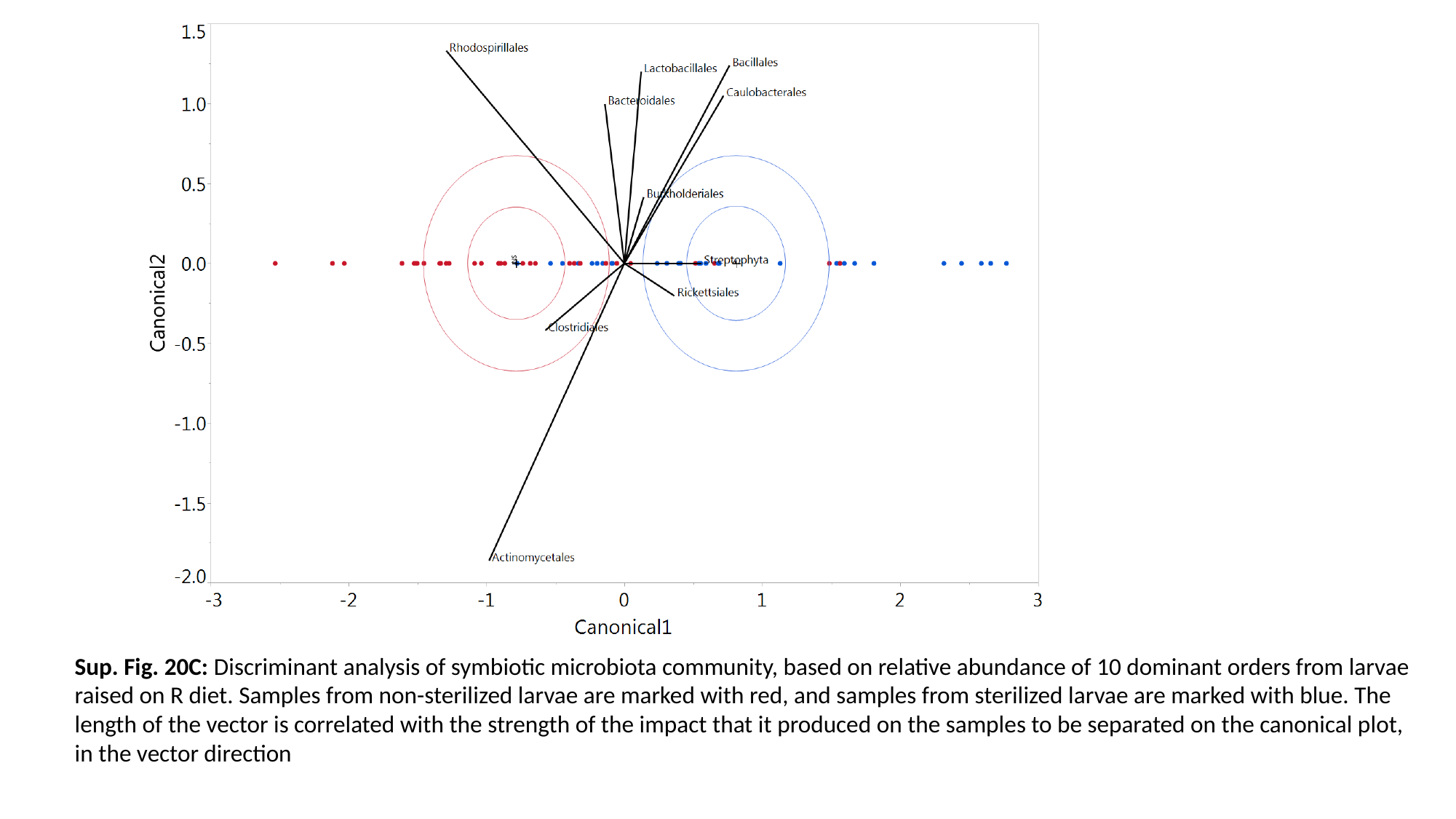

Sup. Fig. 20C: Discriminant analysis of symbiotic microbiota community, based on relative abundance of 10 dominant orders from larvae raised on R diet. Samples from non-sterilized larvae are marked with red, and samples from sterilized larvae are marked with blue. The length of the vector is correlated with the strength of the impact that it produced on the samples to be separated on the canonical plot, in the vector direction

### Slide 50
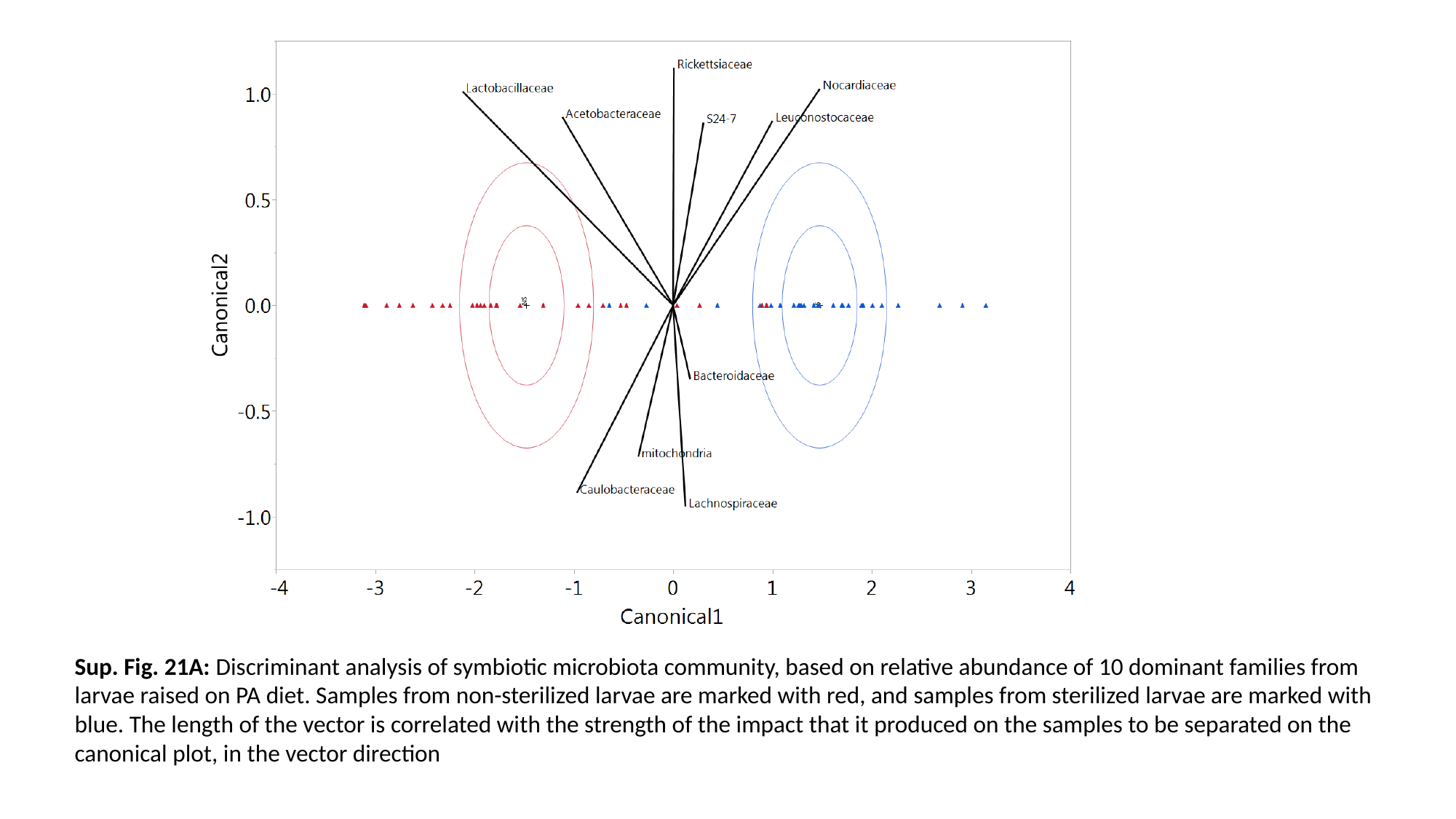

Sup. Fig. 21A: Discriminant analysis of symbiotic microbiota community, based on relative abundance of 10 dominant families from larvae raised on PA diet. Samples from non-sterilized larvae are marked with red, and samples from sterilized larvae are marked with blue. The length of the vector is correlated with the strength of the impact that it produced on the samples to be separated on the canonical plot, in the vector direction

### Slide 51
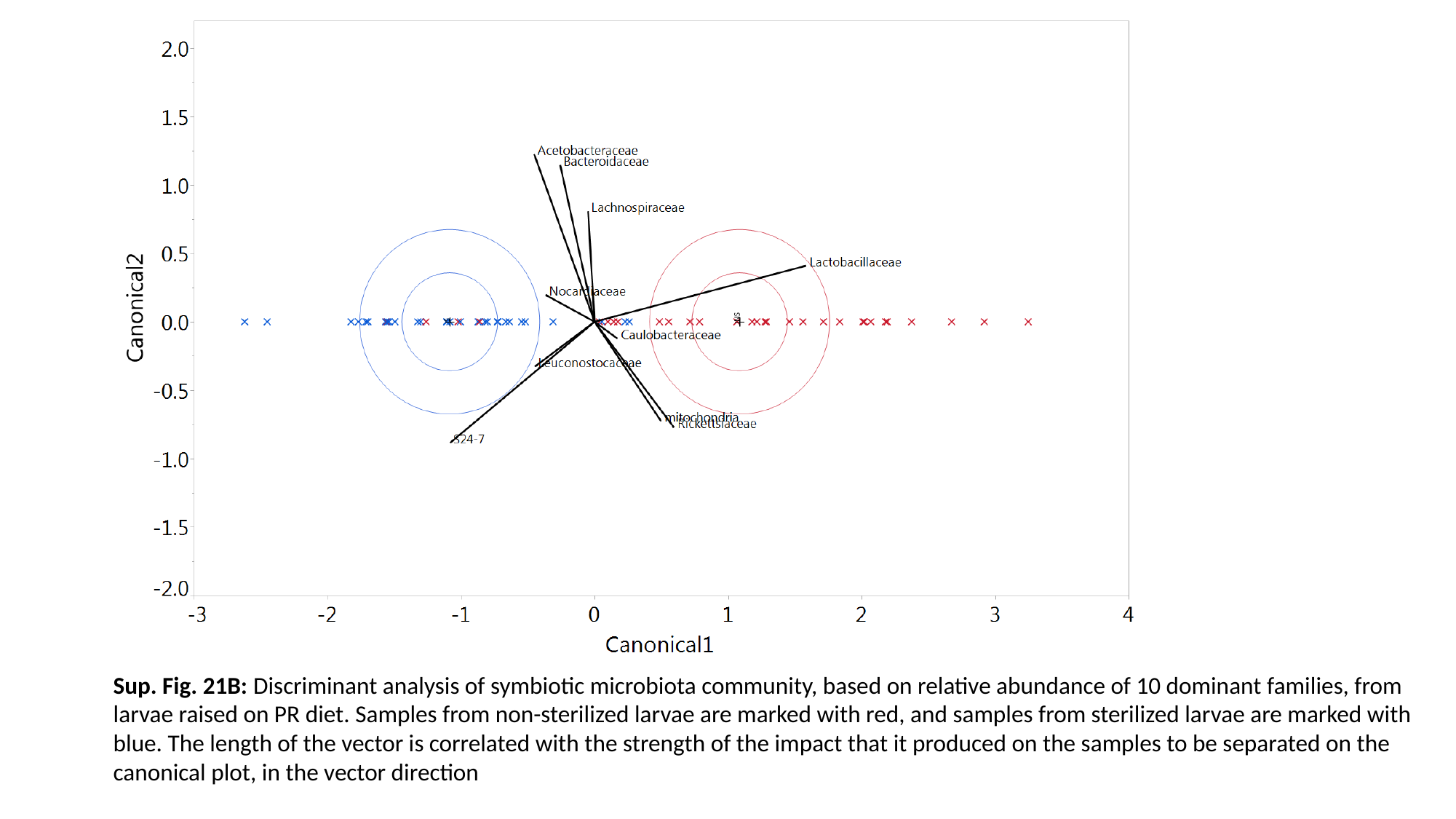

Sup. Fig. 21B: Discriminant analysis of symbiotic microbiota community, based on relative abundance of 10 dominant families, from larvae raised on PR diet. Samples from non-sterilized larvae are marked with red, and samples from sterilized larvae are marked with blue. The length of the vector is correlated with the strength of the impact that it produced on the samples to be separated on the canonical plot, in the vector direction

### Slide 52
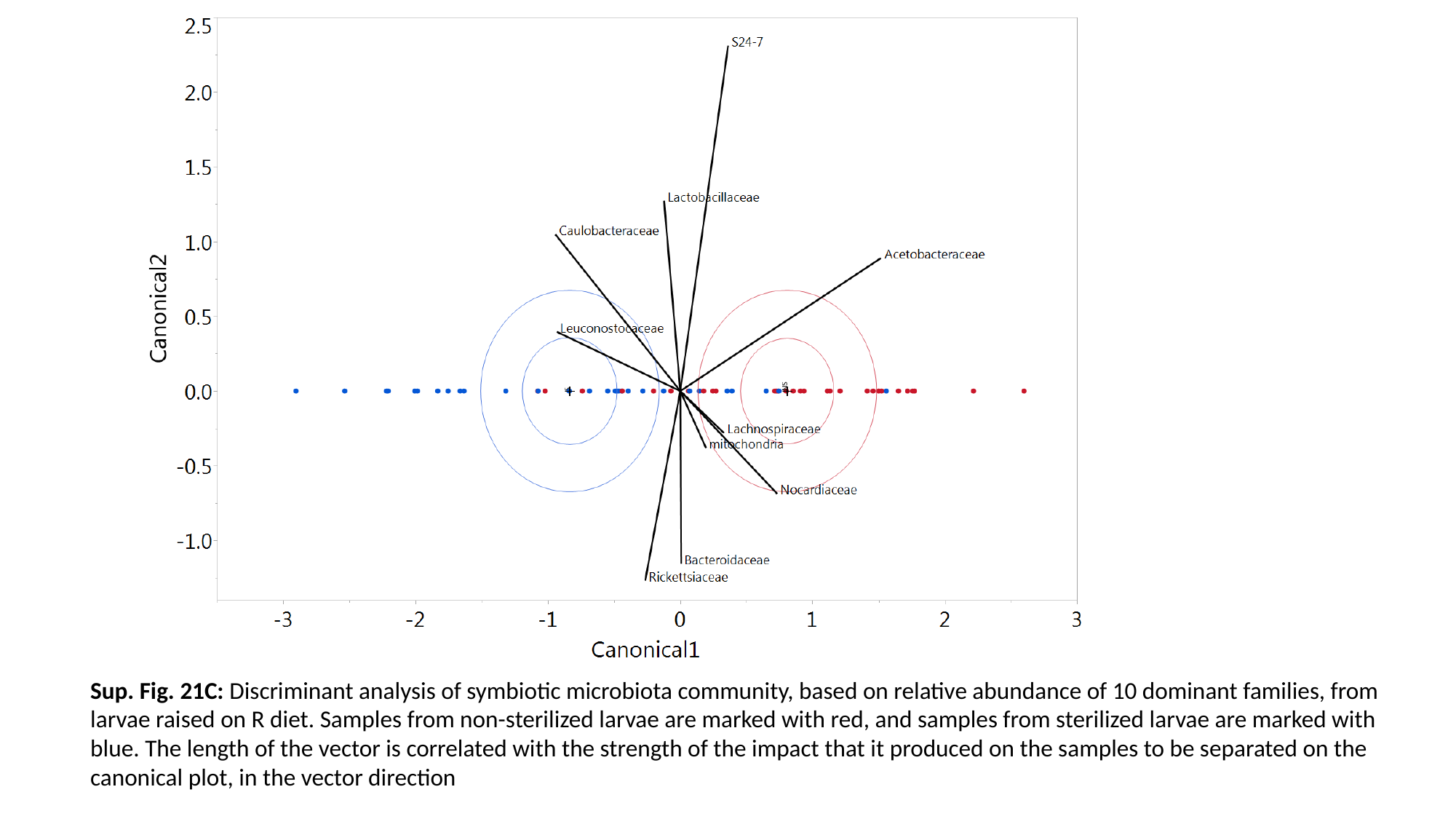

Sup. Fig. 21C: Discriminant analysis of symbiotic microbiota community, based on relative abundance of 10 dominant families, from larvae raised on R diet. Samples from non-sterilized larvae are marked with red, and samples from sterilized larvae are marked with blue. The length of the vector is correlated with the strength of the impact that it produced on the samples to be separated on the canonical plot, in the vector direction

### Slide 53
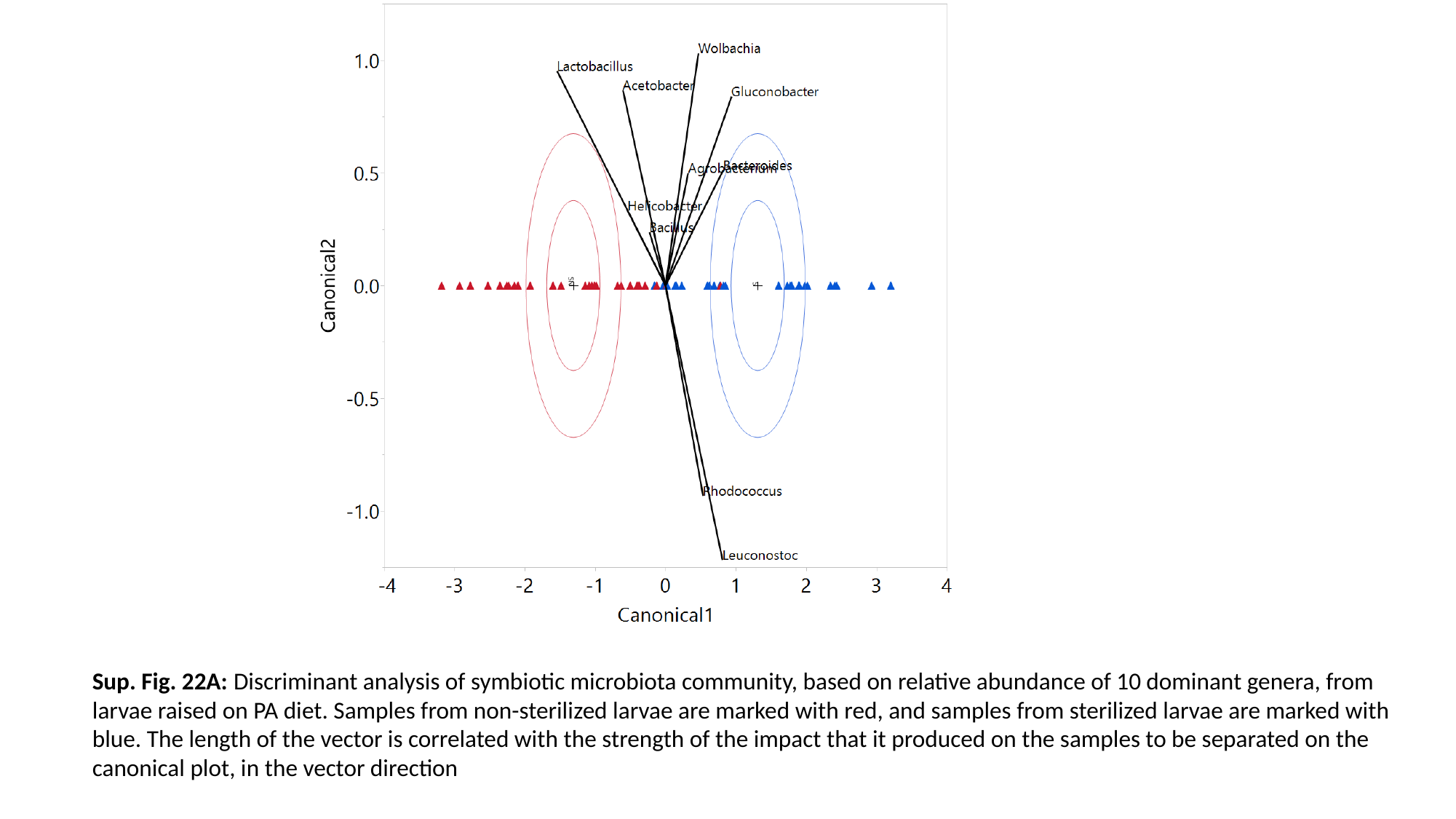

Sup. Fig. 22A: Discriminant analysis of symbiotic microbiota community, based on relative abundance of 10 dominant genera, from larvae raised on PA diet. Samples from non-sterilized larvae are marked with red, and samples from sterilized larvae are marked with blue. The length of the vector is correlated with the strength of the impact that it produced on the samples to be separated on the canonical plot, in the vector direction

### Slide 54
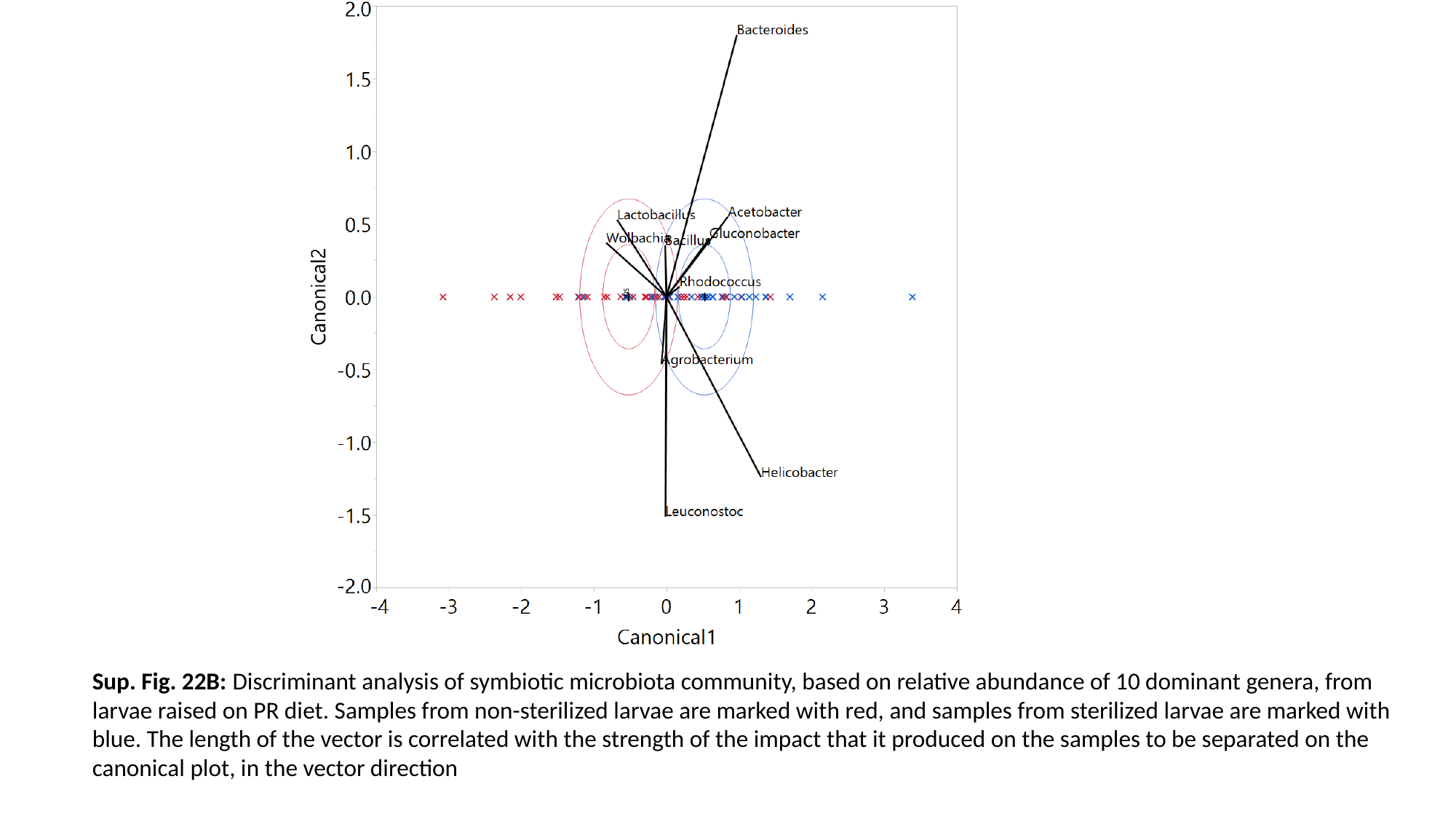

Sup. Fig. 22B: Discriminant analysis of symbiotic microbiota community, based on relative abundance of 10 dominant genera, from larvae raised on PR diet. Samples from non-sterilized larvae are marked with red, and samples from sterilized larvae are marked with blue. The length of the vector is correlated with the strength of the impact that it produced on the samples to be separated on the canonical plot, in the vector direction

### Slide 55
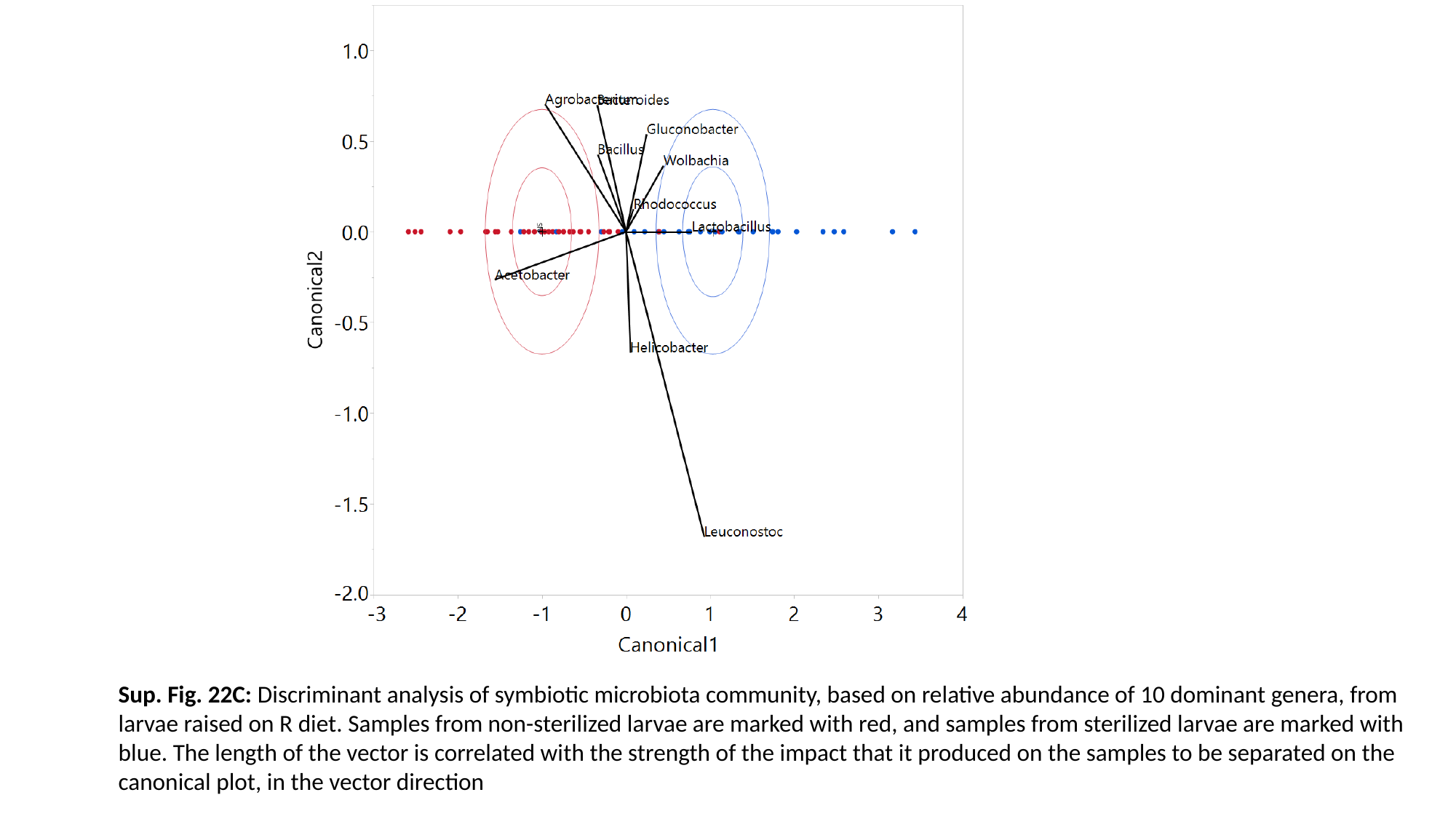

Sup. Fig. 22C: Discriminant analysis of symbiotic microbiota community, based on relative abundance of 10 dominant genera, from larvae raised on R diet. Samples from non-sterilized larvae are marked with red, and samples from sterilized larvae are marked with blue. The length of the vector is correlated with the strength of the impact that it produced on the samples to be separated on the canonical plot, in the vector direction

### Slide 56
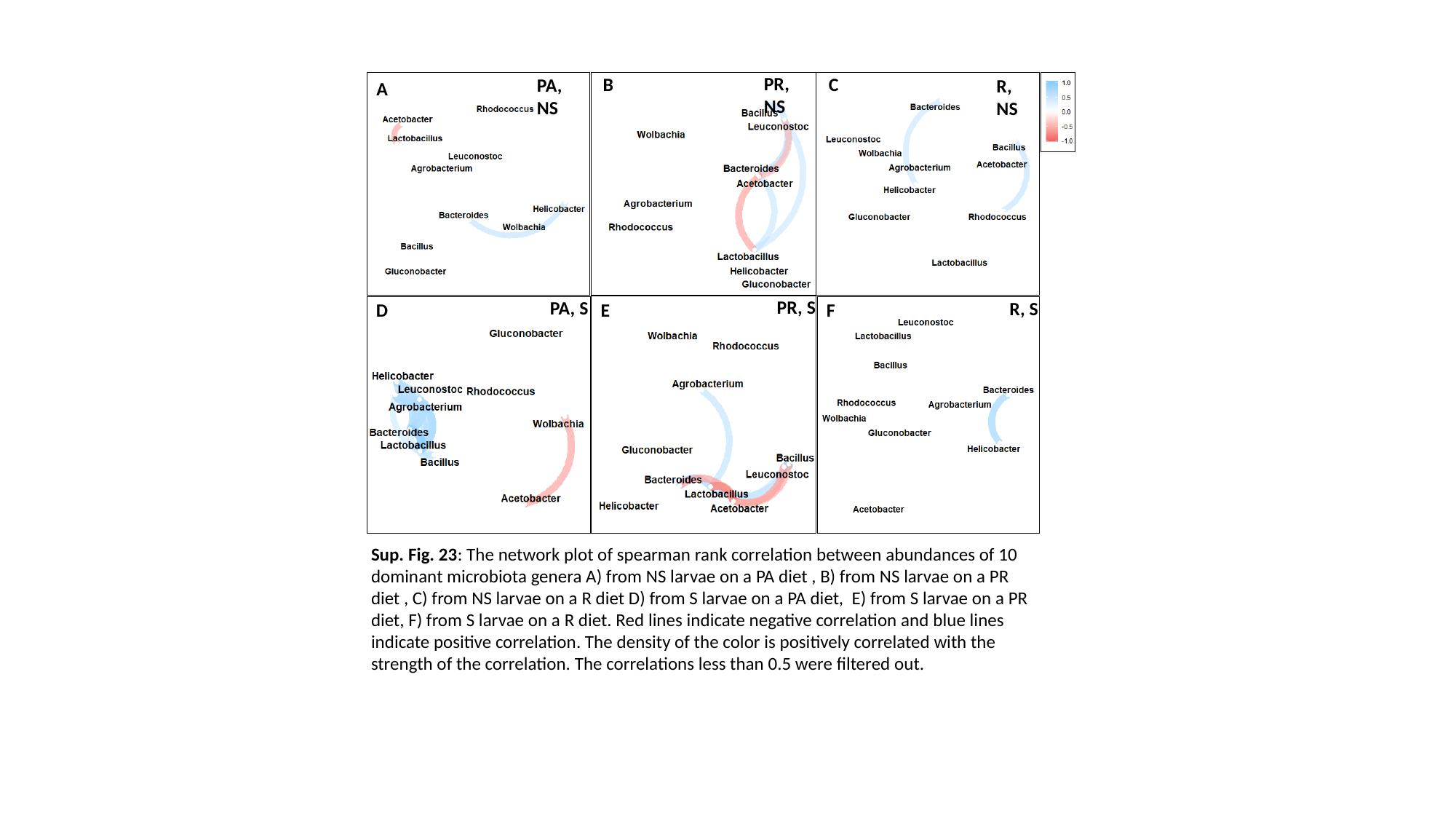

PR, NS
B
C
PA, NS
R, NS
A
PR, S
PA, S
R, S
D
E
F
Sup. Fig. 23: The network plot of spearman rank correlation between abundances of 10 dominant microbiota genera A) from NS larvae on a PA diet , B) from NS larvae on a PR diet , C) from NS larvae on a R diet D) from S larvae on a PA diet, E) from S larvae on a PR diet, F) from S larvae on a R diet. Red lines indicate negative correlation and blue lines indicate positive correlation. The density of the color is positively correlated with the strength of the correlation. The correlations less than 0.5 were filtered out.

### Slide 57
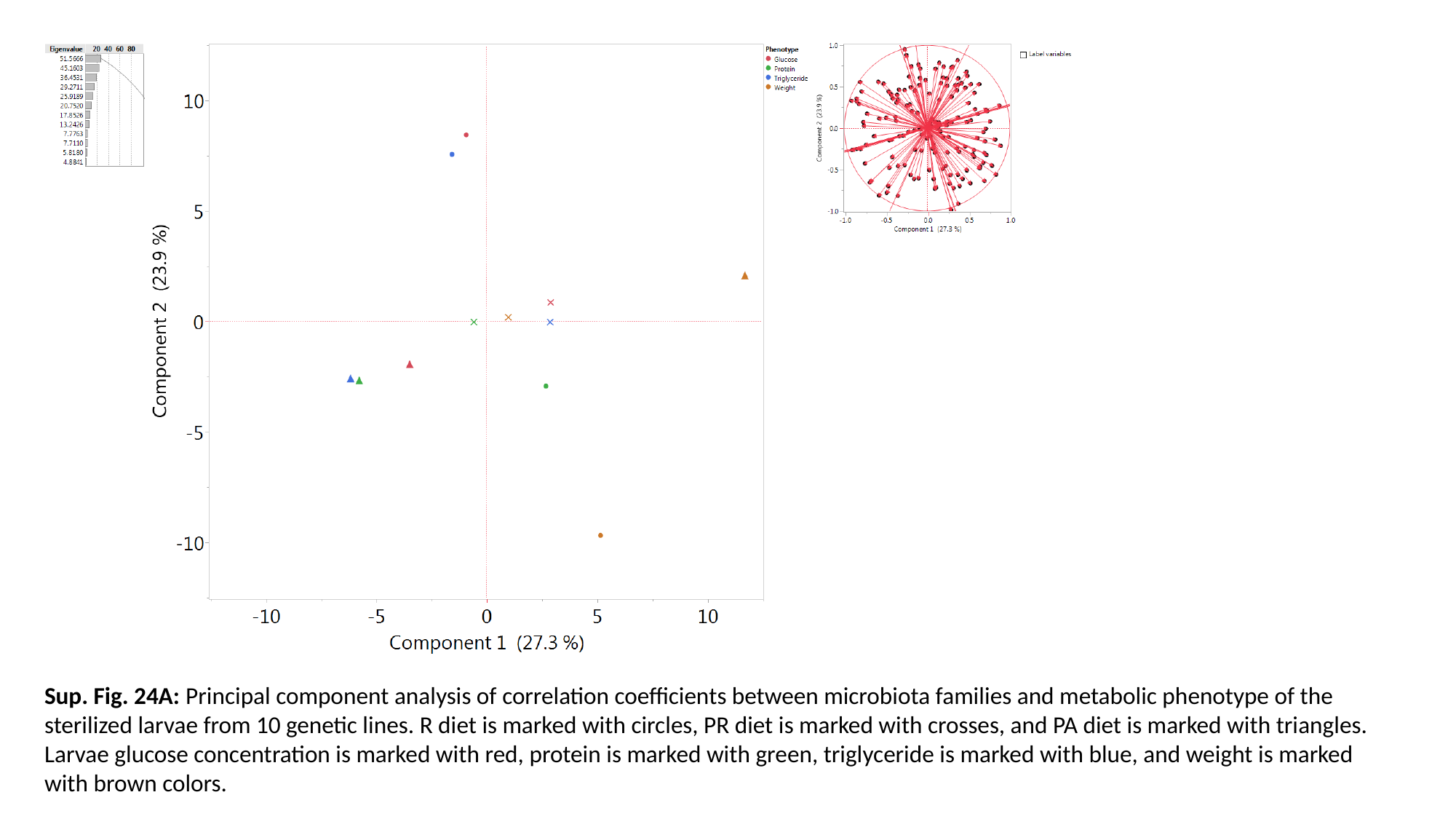

Sup. Fig. 24A: Principal component analysis of correlation coefficients between microbiota families and metabolic phenotype of the sterilized larvae from 10 genetic lines. R diet is marked with circles, PR diet is marked with crosses, and PA diet is marked with triangles. Larvae glucose concentration is marked with red, protein is marked with green, triglyceride is marked with blue, and weight is marked with brown colors.

### Slide 58
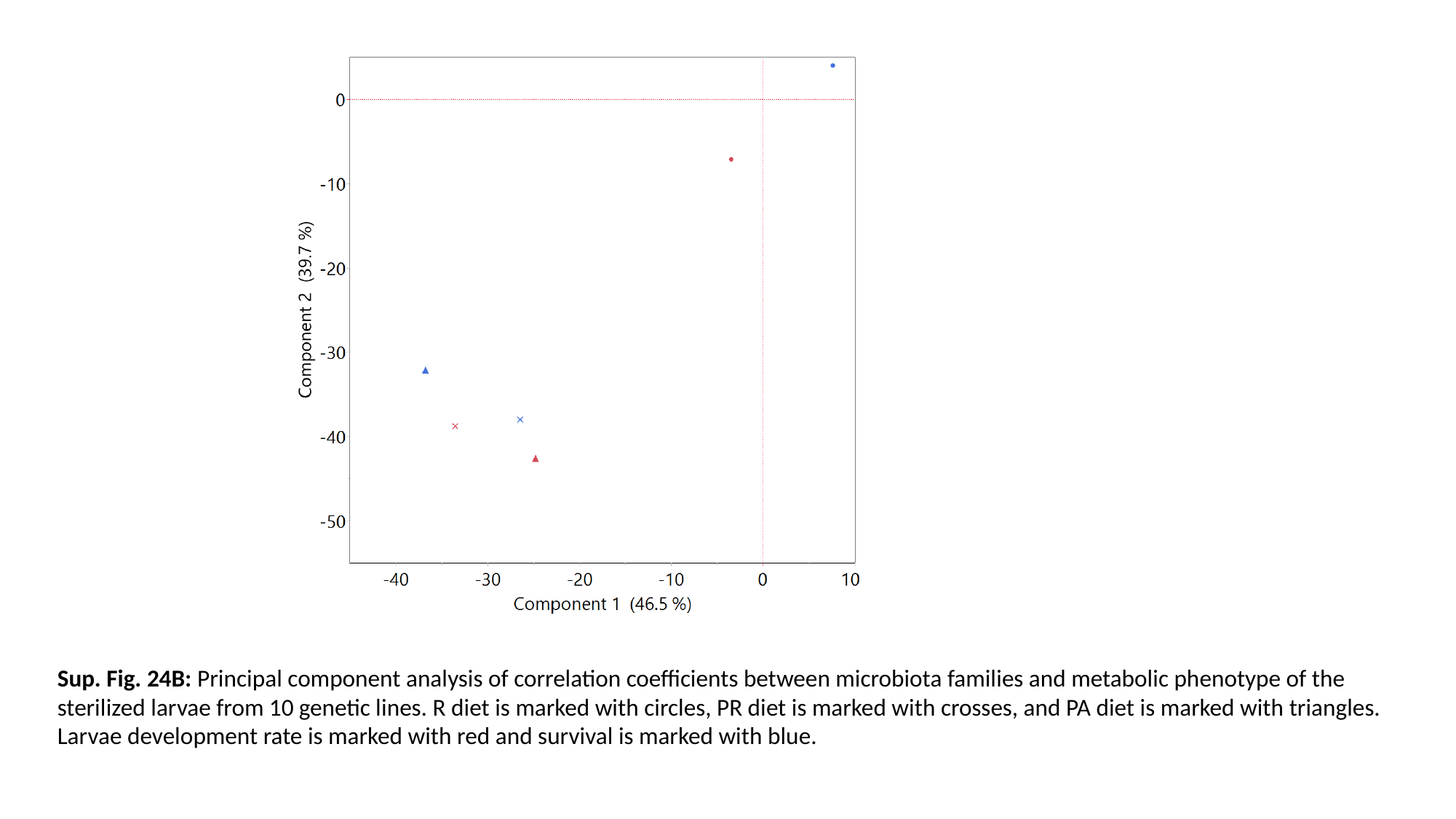

Sup. Fig. 24B: Principal component analysis of correlation coefficients between microbiota families and metabolic phenotype of the sterilized larvae from 10 genetic lines. R diet is marked with circles, PR diet is marked with crosses, and PA diet is marked with triangles. Larvae development rate is marked with red and survival is marked with blue.
